# Substrate traits shape the structure of microbial community engaged in metabolic division of labor

**DOI:** 10.1101/2020.11.18.387787

**Authors:** Miaoxiao Wang, Xiaoli Chen, Yue-Qin Tang, Yong Nie, Xiao-Lei Wu

## Abstract

Metabolic division of labor (MDOL) is widespread in nature, whereby a complex metabolic pathway is shared between different strains within a community for mutual benefit. However, little is known about how the mutual interactions in the microbial community engaged in MDOL are regulated. We hypothesized that when degradation of an organic compound is carried out via MDOL, the substrate traits (i.e., concentration and its toxicity) modulate the benefit allocation between the two microbial populations, thus affecting the structure of this community. We tested this hypothesis by combining mathematical modelling with experiments using engineered synthetic microbial consortia. Numerous modelling analyses suggested that the proportion of the population executing the first metabolic step can be simply estimated by Monod-like formulas governed by substrate traits. The model and the proposed formula quantitatively predicted the structure of our synthetic consortia composed of two strains degrading salicylate through MDOL. Individual-based modelling and colony pattern formation assays further indicated that our rule is also applicable to estimating community structure in spatially structured environments. Our results demonstrate that the structure of the microbial communities can be quantitatively predicted from simple environmental factors, such as substrate concentration and its toxicity, which provides novel perspectives on understanding the assembly of natural communities, as well as insights into how to manage artificial microbial systems.

## Introduction

In natural environments, microorganisms rarely live autonomously; instead, they interact with other individuals to form complex communities, in which they secrete a variety of toxins to compete with each other, or share metabolites to mutually benefit their survival. Among diverse modes of microbial interaction, metabolic division of labor (MDOL) is one of the most widespread phenomena, where distinct populations perform different but complementary steps of the same metabolic pathway [1-4]. MDOL controls numerous ecologically and environmentally important biochemical processes. One important aspect of microbial metabolism implemented by MDOL is the degradation of a variety of complex organic compounds, including PAHs [5, 6], pesticide [7-10], plastics [11], antibiotics [12], or polysaccharides [13, 14]. Bacterial degradation of these complex substrates is usually mediated by long metabolic pathways via a number of intermediates. While these pathways often remain intact within one population, they are frequently found segregated across different members within a community in a MDOL manner. Typical examples include syringate degradation via sequential cross-feeding between *Acetobacterium woodii* and *Pelobacter acidigallici* [5], phenanthrene degradation between *Marinobacter* sp. N4 and other PAH-degrading microbes in marine environments [6], as well as atrazine degradation through MDOL within four bacterial species [9]. However, little is known about how microbial communities engaged in MDOL are regulated [15].

The substrate whose concentration spatially and temporally fluctuates in the marine [16], soil [17] and wastewater [18] environments, acts as one of the most important conditions that govern the performance of the microbial communities [19-21]. Firstly, the concentration of substrates regulates the growth of microbial populations according to the Monod equation [22]. Secondly, many substrates such as PAHs [23, 24], pesticide [7-10], and antibiotics [12], are toxic to bacterial cells, inhibiting their growth. Increasing substrate concentration enhances resource availability of a population that benefit its growth, but also potentially increases the toxic effects of substrate that harms its growth (e.g., growth kinetics may follow the equations integrated with toxic terms [25]). Thus, concentration and toxicity of substrate profoundly affect the fitness of its microbial degraders [24, 26, 27]. However, it still remains ill-defined how substrate straits affect the relative fitness of different strains involved in a community, and thus govern the structure of the community. As structure of a community is fundamental to determine its functioning [28, 29], revealing this question is fundamental for managing such microbial systems for the removal of serious pollutants.

Distinct from the pure culture, the effects of substrate on different populations involved in a MDOL community may vary quite a lot. Firstly, asymmetric benefit allocation exists between different populations in the MDOL community. In MDOL communities that degrade organic compounds, only the population performing the last steps can produce the growth resources (such as small organic acids) that support the bacterial growth (Supplementary Figure 1). Therefore, the population performing the last steps can preferentially acquire and privatize these nutrients (which we henceforth call *product privatization*), thus acquiring the greater benefit, while the other members have to collect nutrients leaked from this population (Figure 1; left of the first row). This uneven allocation of limited resources generally benefits the population that executes the last steps (we henceforth named this population the ‘Embezzler’, analogous to a human worker responsible for the final step of an assembly line, who pockets the final product and fails to share profits with other workers). This phenomenon has been observed in many recent studies [7, 10, 30]. Increasing substrate concentration would enhance the flux of metabolites [31, 32]. Because the Embezzler only have a limited capacity of consuming the final product, increased metabolic flux causes more product released from the Embezzler cells, in turn facilitating the growth of the other population (Figure 1; Right of the first row). Secondly, substrate toxicity exerts different influences on different members. The population performing the first step transforms the toxic substrate to the intermediates (we named it the ‘Detoxifier’ henceforth), which helps it possess a lower intracellular concentration of the toxic substrate (Figure 1; The second row), resulted in that the toxic substrate is less harmful to the Detoxifier than to Embezzler. Accordingly, Detoxifier is favored when the substrate is toxic.

**Figure 1.**
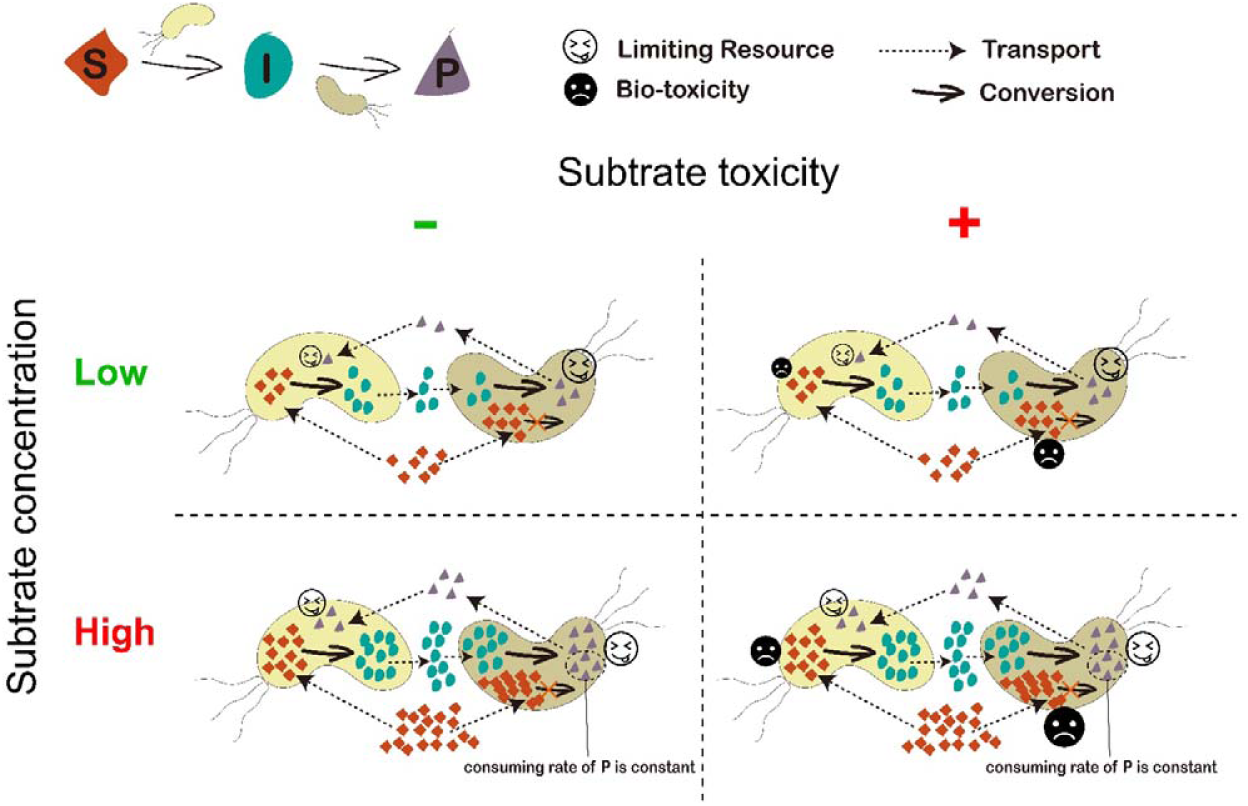
Hypothesis for how substrate concentration and toxicity govern the structure of community engaged in MDOL. In a community degrading an organic compound through metabolic division of labor (MDOL), final product was assumed to be the sole resource and was synthesized by the strain performing the second step. Therefore, this strain will obtain more nutrients (denoted as bigger ‘smiling face’), while the other strain has to collect product released from this population (denoted as smaller ‘smiling face’). Thus, the last population was named ‘Embezzler’. However, increasing the concentration of the substrate (vertical axis) improves the flux of the pathway. Since the P consuming ability of Embezzler cells is limited (dashed box), increasing the concentration will lead to higher final product leakiness, favoring the growth of the first population. Moreover, introducing substrate biotoxicity (horizontal axis) also favors the first population, because it converts this toxic substrate (denoted as smaller sad face), resulting in lower intracellular substrate concentration compared to that of the Embezzler cells (denoted as bigger sad face). Thus, the first population was named ‘Detoxifier’.

It is important to reveal the effects of substrate concentration and toxicity on the structure of the MDOL community. To test the above two hypotheses and reveal how substrate traits shape the structure of microbial community engaged in MDOL, in this study, we combined mathematical modelling and experimentation using a synthetic microbial community. We also tested whether these effects are different when the community grows in spatially well-mixed and structured environments.

## Results

### Testing of our hypotheses in a well-mixed system

#### An ODE model for modelling the dynamics of a community engaged in MDOL

To test our hypotheses on the effects of substrate concentration and its toxicity, we built a mathematical model to simulate the dynamics of a community engaged in metabolic division of labor (MDOL) in a well-mixed system. The dimensionless form of this model is composed of 11 ordinary differential equations (ODEs; Eqn. [4] - Eqn. [13] in Methods). As summarized in Figure 2A, we considered the degradation of an organic substrate (S) into an intermediate metabolite (I), before being degraded to the final product (P). We assumed that two strains carry out this pathway via MDOL, with the first strain only executing the first step, and the second only executing the second. Initially, only S was supplied and the initial concentration was parameterized by *s*_*0*_ (nondimensional). Importantly, based on our hypothesis of ‘Embezzler behavior’, we assumed that, P, which is synthesized by the second strain, is the sole available resource for the growth of both strains. As a result, the second strain possesses the advantage of preferentially acquiring the resource, while the first strain only obtains those growth-limiting resource that is leaked from the second strain. Therefore, the second strain behaves as an ‘Embezzler’. Moreover, biotoxicity of the substrate was imposed (Supplementary Table 3;[25]) to the growth function, and the toxic strength was mediated by parameter θ. Thus, for the scenarios where substrate is assumed to be toxic, the strain executing the first step behaves as a ‘Detoxifier’. Details about the model are described in Supplementary Information S1.

**Figure 2.**
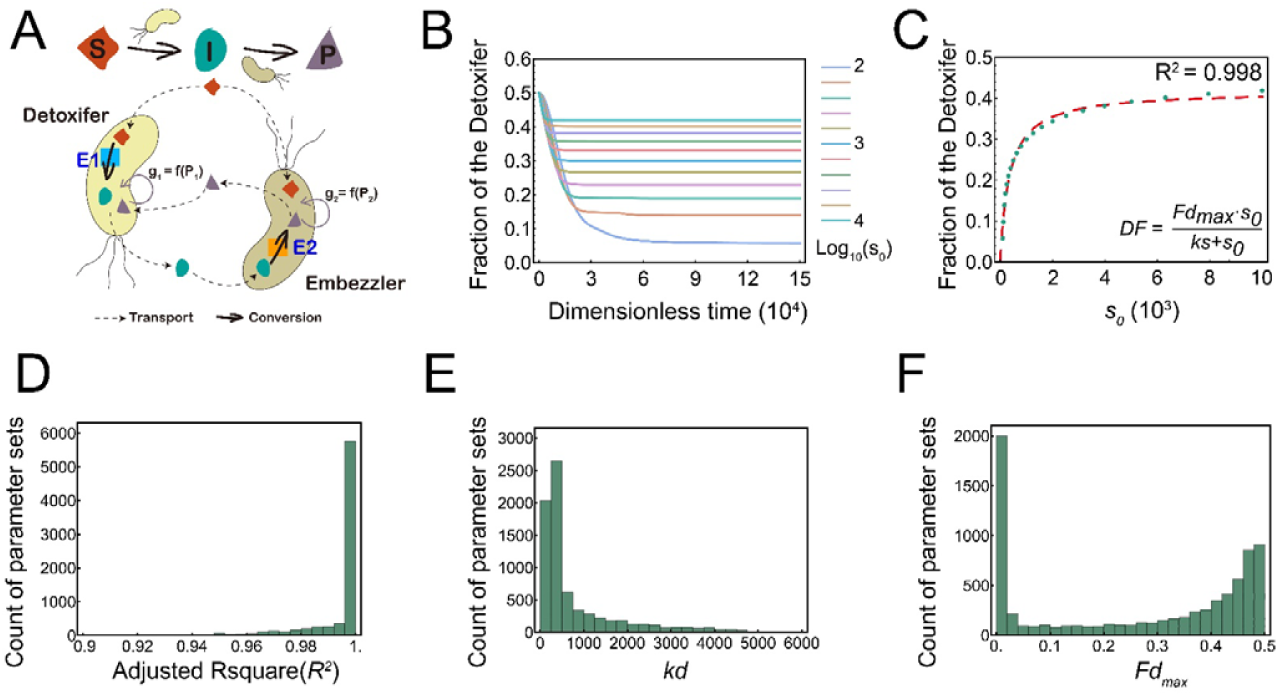
Simulation of the ordinary differential equation (ODE) model excluding substrate toxicity. (A) Schematic diagram showing the basic assumptions of our ODE model without including substrate toxicity. (B-C) A representative case shows how substrate concentration affects the structure of a MDOL community. The simulation dynamics of the fraction of Detoxifier population with the conditions of different initial substrate concentrations are shown in (B). The relationship between substrate concentration and stead-state fraction of Detoxifier is shown in (C). In (C), the green dots denote the simulated stead-state fraction of Detoxifier, and the red dashed line shows the plot of the best fitting function using Eqn. [1]. Parameter values used in these simulations: *y* =10^−4^, *Cp* = 10, *bg* = 1, *a*_*1*_ = 10000, *a*_*2*_ = 1000, *β*_*2*_ = 1, *γ*_*s*_ = 1, *γ*_*i*_ =1, *γ*_*p*_ = 1, *ρ* =10^−2^. The best fitting value of *ks* in this case is 35.3, and that of *Fd*_*max*_ is 0.417. (D-F) Distributions of Adjusted R^2^ (D) of the fitting functions, best fitting value of *ks* (E) and *Fd*_*max*_ (F) in the second-round simulations that does not include substrate toxicity, using 7776 parameter value combinations of the five key parameters (*a*_*1*_, *γ*_*s*_, *γ*_*i*_, *γ*_*p*_, and *Cp*).

#### Analysis of the ODE model indicates initial substrate concentration affects the structure of a MDOL community

To test our first hypothesis stating that substrate concentration affects the structure of the community, we analyzed our ODE model omitting substrate toxicity (Figure 2A). As the dimensionless model contains 11 independent parameters (Supplementary Table 4) that may affect the structure of the community, we performed a first round of numerical simulations using 885,735 parameter sets considering realistic value ranges of all the parameters (Supplementary Information S1.3; Supplementary Table 4). Our analysis showed that the Embezzler population dominated the steady-state community in all these simulations (Supplementary Figure 2; no toxic scenarios, i.e., steady-state frequencies of Detoxifier are lower than 0.5), which was in agreement with our basic assumption of product privatization. Multivariate regression analyses further suggested that six key parameters played vital roles in shaping the structure of MDOL community (Supplementary Table 4; Supplementary Figure 3A; *p* < 0.01 and the fitting coefficient values over 0.01). Notably, *s*_*0*_ was second most important according to the absolute value of the fitting coefficient. *s*_0_ positively correlated with the steady-state proportion of the Detoxifier population, suggesting that a higher initial substrate concentration favors the Detoxifier, consistent with our first hypothesis.

Through the second round of simulations (Supplementary Information S1.3), We found that when all other five key parameters were kept constant, the steady-state proportion of the Detoxifier population (*DF*) increased with an increase of the initial substrate concentration (Figure 2B and 2C), and can be estimated by a Monod-like formula using *s*_*0*_ as the function argument (Figure 2C),

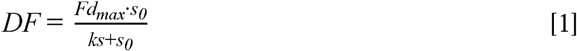

Here, *Fd*_*max*_ represents the maximum proportion of the Detoxifier populations when substrate is non-toxic; *ks* represents the half-saturation constant. Our analysis indicated that the simulation results of all tested parameter sets can be accurately fitted to Eqn. [1] (Figure 2D, values of Adjusted R^2^ mostly over 0.95), although the best fitting of *Fd*_*max*_ and *ks* were affected by the values of other five key parameters (Figure 2E and 2F; Supplementary Information S1.3; Supplementary Table 5; Supplementary Figure 4-5). Together, these results suggest that, in the absence of substrate toxicity, the proportion of the Detoxifier population increases nonlinearly with the increase of the initial substrate concentration, and maintains a maximum value.

To investigate why substrate concentration governs the structure of a community, we next analyzed the intracellular and extracellular concentration of final product of the two populations. We found that with the increase of initial substrate concentration, the fraction of final product released by the Embezzler population increased (Supplementary Figure 6A-H; Supplementary Figure 6I, Red dots). As a consequence, the Detoxifier obtained more product from the environment, resulting in a higher intracellular product concentration, gradually approaching that of the Embezzler. Moreover, based on the first hypothesis, the intracellular product concentration of the Detoxifier should never exceed that of the Embezzler, even if the substrate concentration was elevated to high levels. This prediction was confirmed by our analyses (Supplementary Figure 6A-H; Supplementary Figure 6I, blue dots). As a result, Embezzler cells still maintained their advantage from privatizing final product. This result suggests that in the absence of substrate toxicity, the benefit from product privatization obtained by the Embezzler population cannot be completely eliminated by simply increasing the substrate concentration. This observation matched with our result that the maximum proportion of the Detoxifier population (*Fd*_*max*_) never exceeded 0.5 (Figure 2F; Supplementary Figure 5). In summary, these results suggest that substrate concentration affects the structure of the community engaged in MDOL by affecting the amount of the final product released by Embezzler (Figure 1; the first row).

#### Analysis of the ODE model indicates that substrate toxicity affects the structure of a MDOL community

To test our second hypothesis, we next employed an ODE model that included the parameter of substrate toxicity (Figure 3A). Applying similar simulation and analysis method as used in the above section (Supplementary Information S1.3), we found that the toxic strength (*θ*) of substrate also played a significant role in structure the MDOL community. *θ* exhibited a significantly positive relationship with the final proportion of the Detoxifier population (Figure 3B; Supplementary Figure 2-3; Supplementary Table 4), in agreement with our second hypothesis. We then upgraded Eqn. [1] to collectively consider the effects of substrate concentration and its toxicity (Figure 3C), as follow

**Figure 3.**
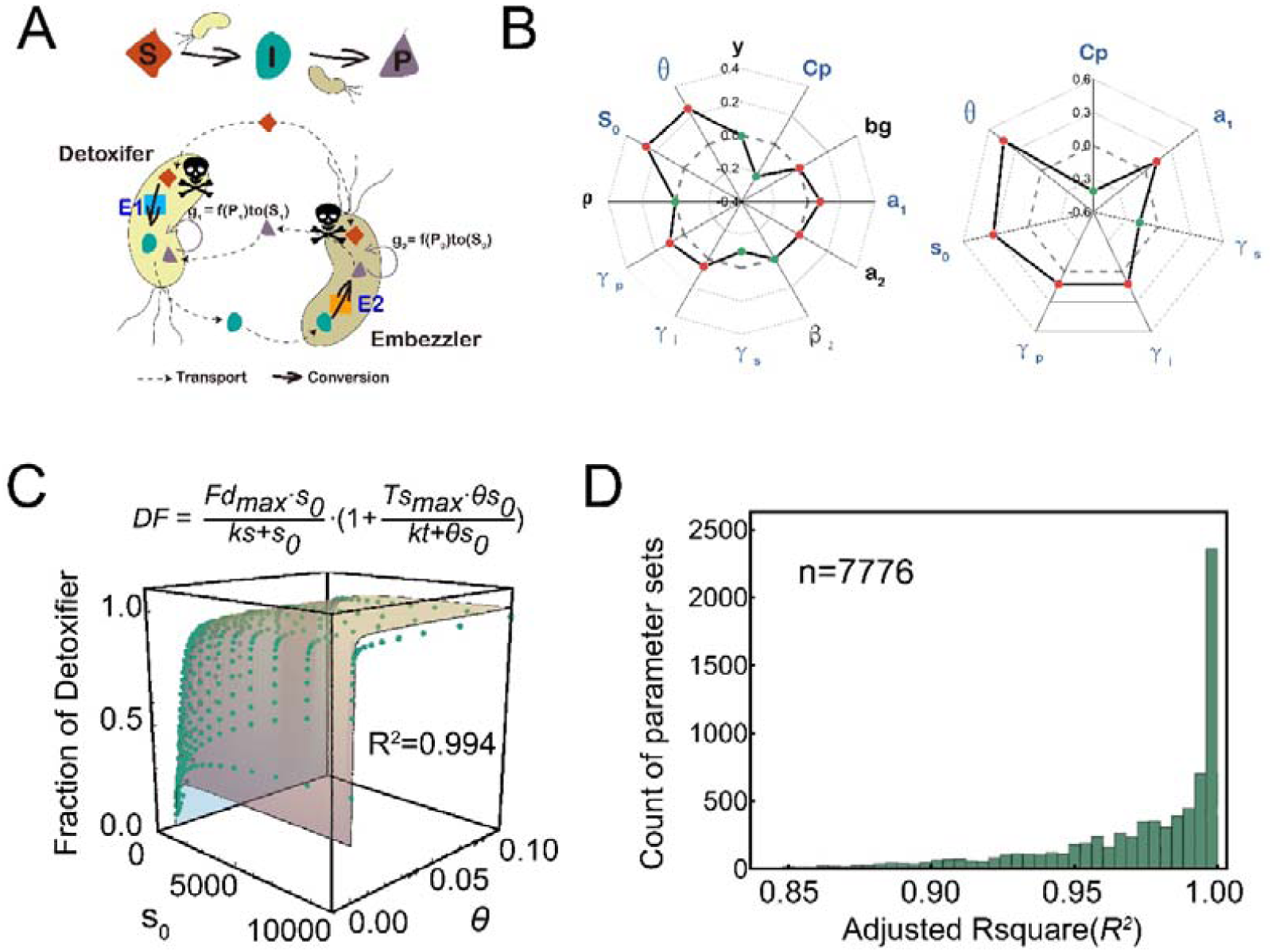
Simulation of the ordinary differential equation (ODE) model that includes substrate toxicity, suggesting that both substrate concentration and its toxicity collectively affect the structure of a community engaged in MDOL. (A) Schematic diagram showing the basic assumptions of our ODE model that includes substrate toxicity. (B) Multiple linear regression analysis of the simulation results of the ODE model showed how the parameters included in the model affect the structure of the MDOL community. Left: results from the first-round simulations that considered all the twelve parameters are shown. Blue font denotes the identified key parameters. Right: results from the second-round simulations that only considered the seven key parameters. The axis of the radar plot denotes the values of fitting coefficients of the parameters from multiple linear regression analyses. Red dots denote the steady-state fraction of Detoxifier is positively correlated with corresponding parameter, while the green dots represent the negative correlation. The origin axis (0) is highlighted by dash line to emphasize the fact that the closer a value is to zero, the smaller the effect on the community structure by the corresponding parameter. The data are also listed in Supplementary Table 4 and Supplementary Table 5. In this analysis, the toxic effects of substrate on population growth were assumed to follow a reciprocal relationship. Results considering other relationships are shown in Supplementary Figure 3. (C) A representative case shows how both substrate concentration and its toxicity collectively affect the stead-state proportion of Detoxifier cells. The green dots denote the simulated stead-state fraction of Detoxifier, and the surface shows the plot of the best fitting function using Eqn. [2]. Parameter values used in these simulations: *y* =10^−4^, *Cp* = 10, *bg* = 1, *a* = 10000, *a* = 1000, *β*_*2*_ = 1, *γ*_*s*_ = 1, *γ* _i_=1, *γ*_*p*_= 1, *ρ =*10^−2^. The best fitting value of *ks, Fd*_*max*_, *kt*, and *TS*_*max*_ in this case are 48.9, 0.423, 0.848, 3.39, respectively. (D) Distributions of Adjusted R^2^ of the fitting functions in the second-round simulations that includes substrate toxicity, using 7776 parameter value combinations of the five key parameters (*a*_*1*_, *γ*_*s*_, *γ*_*i*_, *γ*_*p*_, and *Cp*).

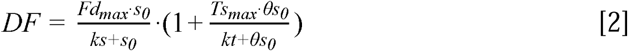

In Eqn. [2], we use term 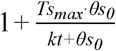 to describe the effect of substrate toxicity on the proportion of the Detoxifier populations. *Ts*_*max*_ represents the maximum fold increase of Detoxifier proportion benefiting from the substrate toxicity; *ks* represents the half-saturation constant of this toxic effect. This term is positively affected by the toxic strength (*θ*) and substrate concentration (*s*_*0*_), since increasing either toxic strength or substrate concentration harms population growth (see Eqn. [12]-[13] in Methods and Supplementary Table 3). Our analyses further indicated that the *DF* values derived from numerical simulations accurately fitted to the values predicted by Eqn. [2] (Figure 3D; values of Adjusted R^2^ mostly over 0.90; see Supplementary Table 5, and Supplementary Figure 7-10 for parameter sensitive analyses). These results suggest that when substrate toxicity was taken into account, the proportion of the Detoxifier population increased with both the initial concentration and the toxic strength of the substrate.

To address why substrate toxicity affects structure of the community, we next analyzed the intracellular and extracellular concentration of both S and P of the two populations. As shown in Supplementary Figure 11, the fraction of final product released by the Embezzler population largely agrees with the result derived from those non-toxic scenarios, suggesting that the presence of substrate toxicity does not change the leakiness of final product from the Embezzler. Our analysis of the S concentration showed that the Detoxifier population generally maintained a lower intracellular concentration level of S than that of the Embezzler (Supplementary Figure 12), due to its conversion of S, thus possessing a growth advantage over the Embezzler population. Based on this mechanism, higher speed of the first reaction, or lower S transport rate, appears to favor the Detoxifier population since these two conditions assist Detoxifier in maintaining a lower intracellular S concentration. Consistent with this corollary, *Ts*_*max*_ was significantly positively correlated with *α*_*1*_ and significantly negatively correlated with *γ*_*s*_ (Supplementary Table 5; Supplementary Figure 10). Overall, these results indicated that the difference in intracellular concentration of substrate is the main reason why substrate toxicity favors the Detoxifier population (Figure 1; second row).

When we assessed the community structure at different conditions of substrate traits, we found that Detoxifier population dominated the community when the substrate concentration and substrate toxicity were sufficiently high (its relative proportion exceeded 50% of the community; Figure 3C; Supplementary Figure 2), suggesting that the benefit from product privatization of the Embezzler can be neutralized by higher substrate concentration and toxicity. This phenomenon is quantitively characterized by Eqn. [2]: the maximum Detoxifier proportion (*Fd*_*max*_) never exceed 0.5 in the absence of substrate toxicity (Supplementary Figure 8), but substrate toxicity can assist Detoxifier in breaking through this constraint, as quantified by the term 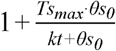.

In summary, our simulations clearly showed that when a compound degradation pathway is executed through MDOL in a community, both increasing substrate concentration and toxicity of the substrate favor the Detoxifier population, resulting in substrate traits to shape the structure of the community.

#### Experimental evaluation of our rule using a liquid culture of a synthetic microbial consortium engaged in MDOL

To experimentally test the prediction from our ODE model, we engineered a synthetic consortium composed of two *P. stutzeri* strains, which cooperatively degrade an organic compound, salicylate, via MDOL (Figure 4A). In this synthetic consortium, strain *P. stutzeri* AN0010 only retained its ability to convert toxic substrate, salicylate to the intermediate catechol [33], behaving as the ‘Detoxifier’; the second strain, *P. stutzeri* AN0001, was only able to metabolize catechol, but possessed the preferential access to the final product, i.e., pyruvate and acetyl-CoA (Figure 4A), the direct carbon source of both strains, thus behaving as the ‘Embezzler’. Details about the strain construction are described in Supplementary Information S3. For simplicity, we henceforth refer to our community as ‘SMC-mdol’.

**Figure 4.**
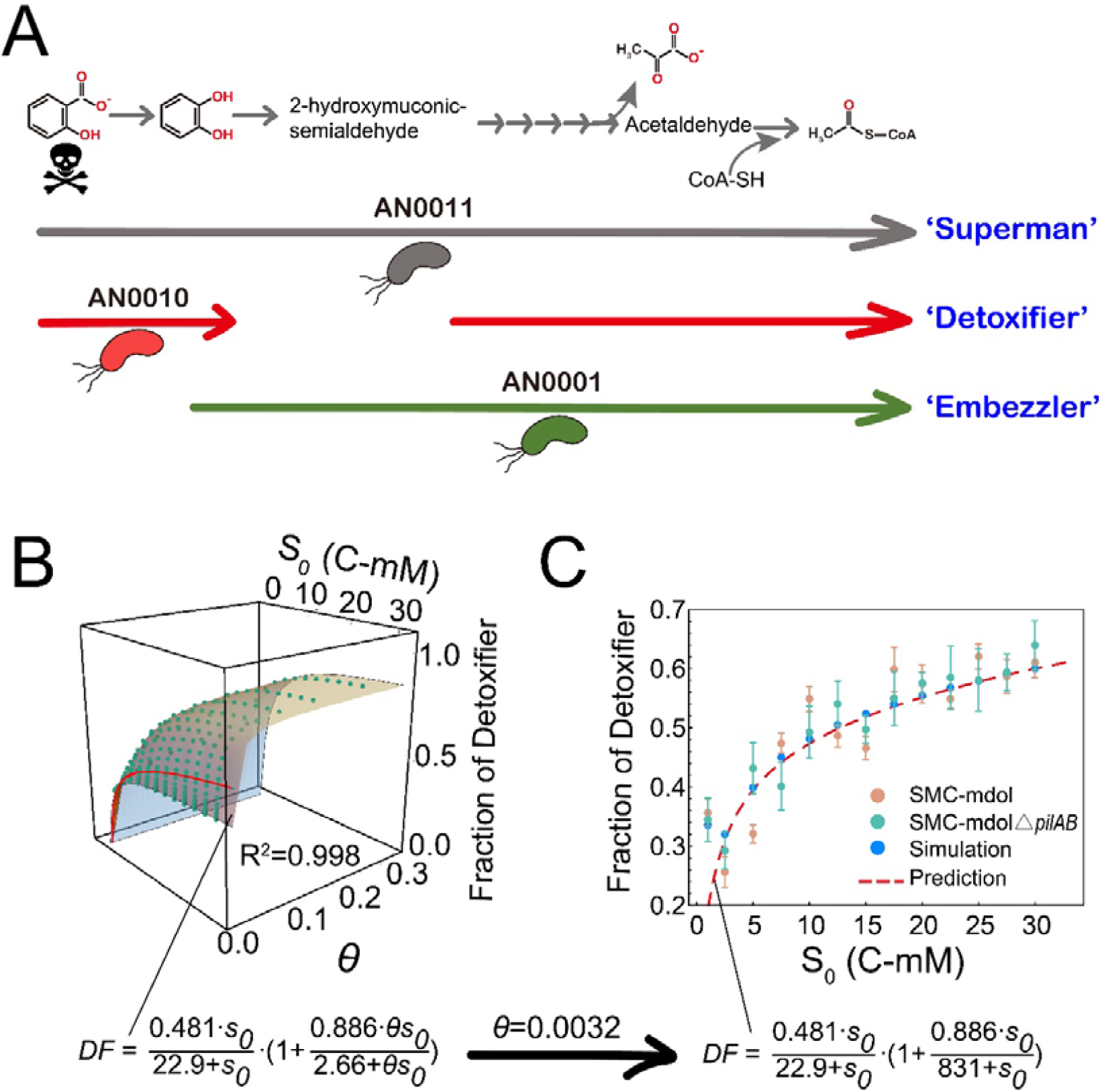
Structure of SMC-mdol in a spatially unstructured system governed by different substrate traits. (A) Design of the SMC-mdol. Shown are the pathway of salicylate degradation in ‘Superman’ strain *P. stutzeri* AN0011, as well as partial pathways carried out by Detoxifier strain AN0010 and Embezzler strain AN0001. Skull marks that salicylate is toxic. (B) Predicting the structure of the synthetic consortium using our ODE model, as well as the derived predictive function using Eqn. [2]. The relationship between the steady-state fraction of the Detoxifier population and substrate concentration (*s*_*0*_), as well as substrate toxic strength (*θ*), was built from our mathematical model using parameters consistent with our experiemental system. Each green dot shows the steady-state fraction of Detoxifier obtained by one simulation accociated with the specific parameter set. The surface diagram shows distribution of the steady-state fraction of Detoxifier predicted by our proposed simple formula. The Red line in the surface denotes the scenarios *θ*=0.0032, which is the toxic strength of salicylate obtained from experiemental measurements. (C) The experimental measured steady-state fractions of Detoxifier in cultures with different salicylate concentrations is consistent with those from mathematical predictions. Note that in the plots, substrate concentrations are shown in dimentional form (*S*_*0*_, Cmmol/L), but in the predictive functions, the fitting analysis was performed using its dimensionless form (*s*_*0*_).

We first derived a function to predict the structure of our synthetic consortium based on our model using experimentally measured or previously reported parameters (Figure 4B; Supplementary Table 6; Supplementary Information S1.3). We quantified the toxicity of salicylate (see Supplementary Information S3.4 for measurement details), and the measured dimensionless value of toxic strength (*θ*) of salicylate was 0.0032 (Supplementary Figure 13). Accordingly, we mathematically predicted the effects of substrate traits on the structure of SMC-mdol, as indicated by the red line in Figure 4B and 4C. In the liquid minimal medium supplemented with different concentrations of salicylate, SMC-mdol exhibited similar dynamics to that from our corresponding ODE simulations (Supplementary Figure 14). The steady-state proportion of Detoxifier population increased from 25.6% ± 2.5% to 61.1% ± 2.6% as a function of initial salicylate concentration (Figure 4C). Moreover, our prediction function accurately estimated the steady-state structure of SMC-mdol, with a predictive power (Adjusted R^2^) of 0.983. Importantly, when the substrate concentration reached high levels, the Detoxifier population dominated the community (i.e., its relative fraction over 50 %), suggesting that substrate toxicity considerably affected the structure of our consortium. Together, these experiments confirmed our simple rule proposed from mathematical modelling, and suggested that the structure of microbial community engaged in MDOL are governed by concentration and toxicity of the substrate.

### Testing our hypotheses in spatially structured environments

In the above modeling and experiments, we investigated how substrate traits affect the structure of a MDOL community, principally by assuming that the substances and cells were well-mixed in the system. However, microorganisms frequently grow in spatially structured environments [34-36]. Previous studies reported that different physical characteristics between the well-mixed and spatially structured systems significantly affected the structure of a community [37-40]. Therefore, we set out to test whether our rule derived from the assumption of a well-mixed system can be expanded to estimate the structure of a MDOL community in spatially structured environments.

#### Individual-based modelling of the dynamics of a MDOL community

To develop a mathematical framework to simulate the dynamics of MDOL community in spatially structured environment, we built an individual-based (IB) model. The basic configuration of our IB model was identical to the framework of our ODE model. Moreover, we assumed that the diffusion of S, I, and P was limited in the IB model, and mediated by their diffusion coefficients (*D*_*s*_, *D*_*i*_, and *D*_*p*_). Details about the IB model are described in Supplementary Information S2.

To test our hypotheses, we ran the IB model using the parameters consistent with our experimental system (Supplementary Table 7), but varied the toxic strength (*θ*) and initial concentration of the substrate (*s*_*0*_). We found that during the colony growth, cell lineages of Detoxifier and Embezzler segregated at frontiers, forming adjacent red and green cell sectors (Figure 5A; Supplementary video 1-4). Analysis of the spatial distribution of S, I, and P suggested that the development of this colony characteristic was mainly attributed to the ‘active layer effect’ reported previously [41]. As S is generally supplied from the outside of the colony, a thin active cell layer formed depending on the penetration of S, I and P (Supplementary video 1-4). Consequently, community structures in the inoculating and expanding regions may differ. Accordingly, we separately analyzed the structures in the inoculating region and expanding region of the colonies (Supplementary Figure 15). We found that with the growth of colony, community structures in the inoculating region changed little, while the community structures in the expanding region shifted over time, gradually approaching a steady-state (Supplementary Figure 16). Therefore, we next investigated how substrate traits affect the steady-state structures of the MDOL community in the expanding regions. the community structure in the expanding region was significantly affected by substrate traits, and can be well estimated by the rule (Eqn. [2]) that we proposed for a well-mixed system (Figure 5B; Supplementary Figure 17). This result indicated that the structure of the MDOL community in spatially structured environments can also be estimated by the proposed simple formula governed by substrate traits.

**Figure 5.**
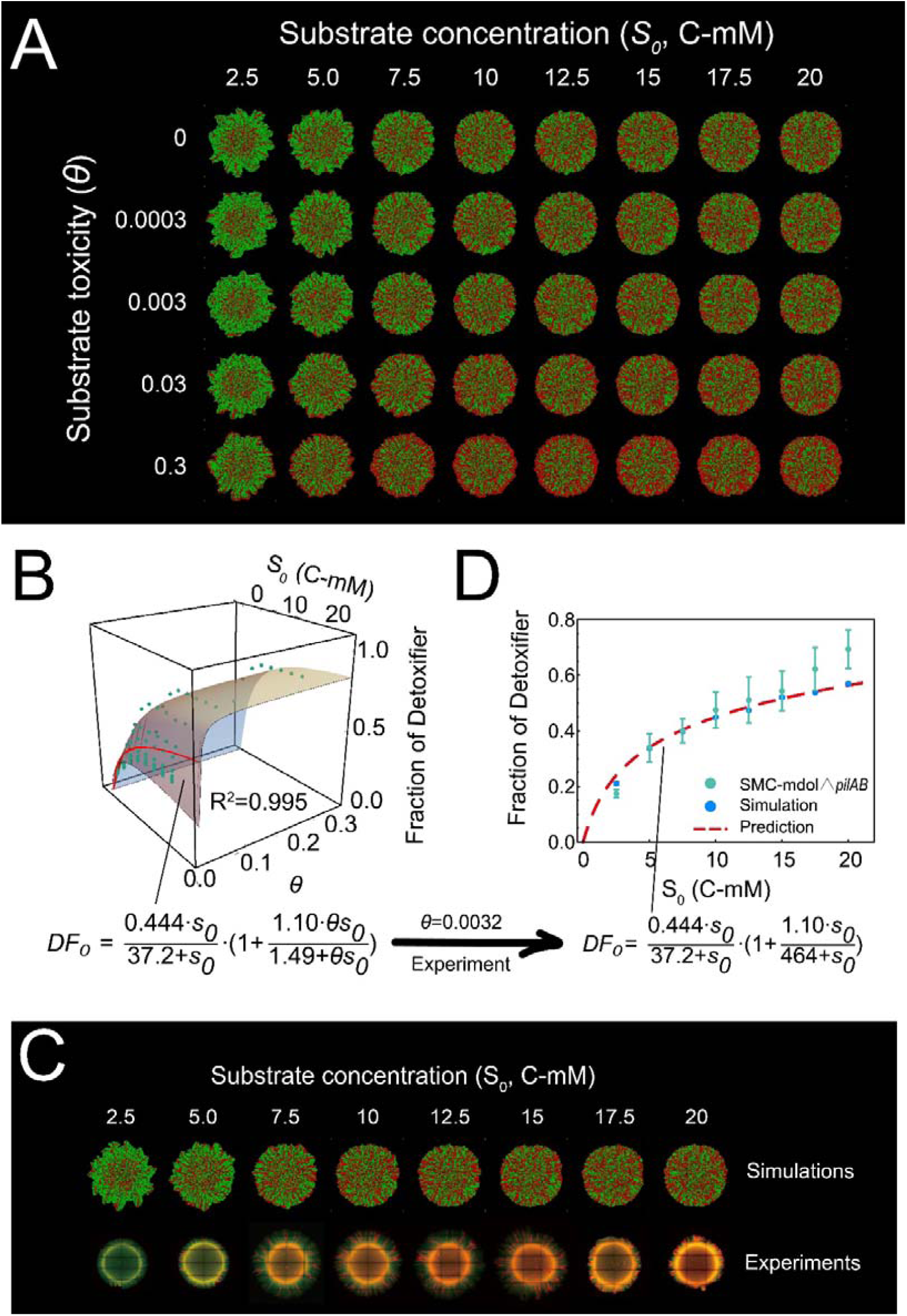
Substrate traits governing the structure of a microbial community engaged in metabolic division of labor (MDOL) in a spatially structured environment. (A) Representative colony patterns from Individual-based (IB) modelling initialized with different substrate traits. Detoxifier cells are shown in red, while Embezzler cells are shown in green. (B) Analysis of community composition in the expanding region of the colonies from IB simulations across eight kinds of initial substrate concentrations and five different toxic strength. Plot shows how both substrate concentration and its toxicity collectively affect the stead-state proportion of Detoxifier. The green dots denote the simulated stead-state fraction of Detoxifier. The surface shows the plot of the best fitting function using Eqn. [2]. The Red line in the surface denotes the scenarios θ=0.0032, which is the toxic strength of salicylate obtained from experimental measurements. (C) Representative colony patterns from the pattern formation assays of SMC-mdol△*pilAB*, as well as the IB simulations using the parameters matched with our synthetic system (Supplementary Table 7), across eight different initial substrate concentrations. (D) The experimental measured steady-state fractions of Detoxifier in the expanding region of these colonies is consistent with those from mathematical predictions. Note that in the plots, substrate concentrations are shown in dimensional form (*S*_*0*_, Cmmol/L), but in the predictive functions, the fitting analyses were performed using its dimensionless form (*s*_*0*_).

We also found that increasing substrate concentration assisted Detoxifier to obtain more product from the environment, thus retaining higher intracellular product concentrations (Supplementary Figure 18). Furthermore, Detoxifier cells possessed a lower intracellular concentration level of S than that of the Embezzler cells in our IB simulations (Supplementary Figure 19); higher speed of the first reaction, or lower S transport rate, also significantly increased the maximum benefit (*Ts*_*max*_) that Detoxifier cells can obtained from substrate toxicity (Supplementary Figure 20; correlation analysis *p*<0.0001), same as our results from ODE modelling. Therefore, same mechanisms as in the well-mixed system are also applicable to explain why substrate traits affects the structure of MDOL community in spatially structured environments.

#### Experimental evaluation of our rule by culturing our synthetic microbial consortium in spatially structured environment

We next experimentally tested our hypotheses in spatially structured environments. Several studies have reported that type IV pilus may affected the microbial colony patterns [42-44]. To directly focus on the effects of substrate traits and avoid the effects of pili, we deleted the *pilA* and *pilB* genes of the both strains involved in our synthetic consortium. This design follows other studies that performed patterning experiments using non-motile strains [45-48]. The derived consortium was named as SMC-mdol△*pilAB*. As shown in Fig. 4C, this strain modification did not change the effects of substrate traits on the structures of the consortium in well-mixed system, as well as the salicylate toxicity to the strains (Supplementary Figure 13).

To test our hypotheses, we cultured SMC-mdol△*pilAB* on an agarose surface to which salicylate was added at different concentrations. The experimentally observed colony patterns were very similar to those observed in the simulations (Figure 5C). We next separately assessed the structures of the consortium in both the inoculating region and expanding region of the colonies. We found that the proportion of Detoxifier population slightly shifted from 40.9% ± 3.5% to 60.0% ± 6.0% in the inoculating region (Supplementary Figure 21), but it largely varied from 17.4% ± 1.5% to 69.0% ± 7.0% in the expanding region (Figure 5D). Importantly, the experimental results of expanding region accurately fitted to our derived prediction function (Figure 5D) with a predicting power (Adjusted R^2^) of 0.982. Together, our simulations and experiments demonstrated that our rules on how substrate traits shape the structure of MDOL community were applicable when this community grew in a spatially structured environment.

#### The effects of substance diffusivity on the structure of the MDOL community

Although the structure of MDOL community in spatially structured and well-mixed environments can both be estimated by Eqn. [2], the estimated parameter values in the prediction functions derived from ODE and IB model are slightly different (Figure 4 and 5), even if we applied identical parameters and equations in these two models (Supplementary Information S2.3). Through mathematical modelling, we revealed that limited mass diffusion is one of the major reasons that lead to this difference (see Supplementary Information 2.2 for detail). Our analyses suggested that higher level of P diffusion favors the Detoxifier (Supplementary Figure 22-23), whereas increasing the diffusion level of I harms the Detoxifier (Supplementary Figure 24-25).

In addition, we found that the diffusion level of substrate has two opposing effects on the structure of MDOL community. On the one hand, higher diffusion level of S benefits Detoxifier (Figure 6A, first row), through thickening the cell’s ‘active layer’ (Figure 6B; [48]), and thus increasing production and secretion of the final product by Embezzler cells. On the other hand, higher diffusion level of S also decreases the fitness of the Detoxifier cells by modifying the concentration gradient of S around the two types of cells, and thus changing relative toxic level of S (Figure 6A, second row; Supplementary Figure 26). Combining these two effects, we formulated a new formula to estimate the structure of MDOL community

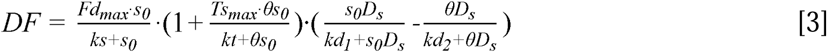

**Figure 6.**
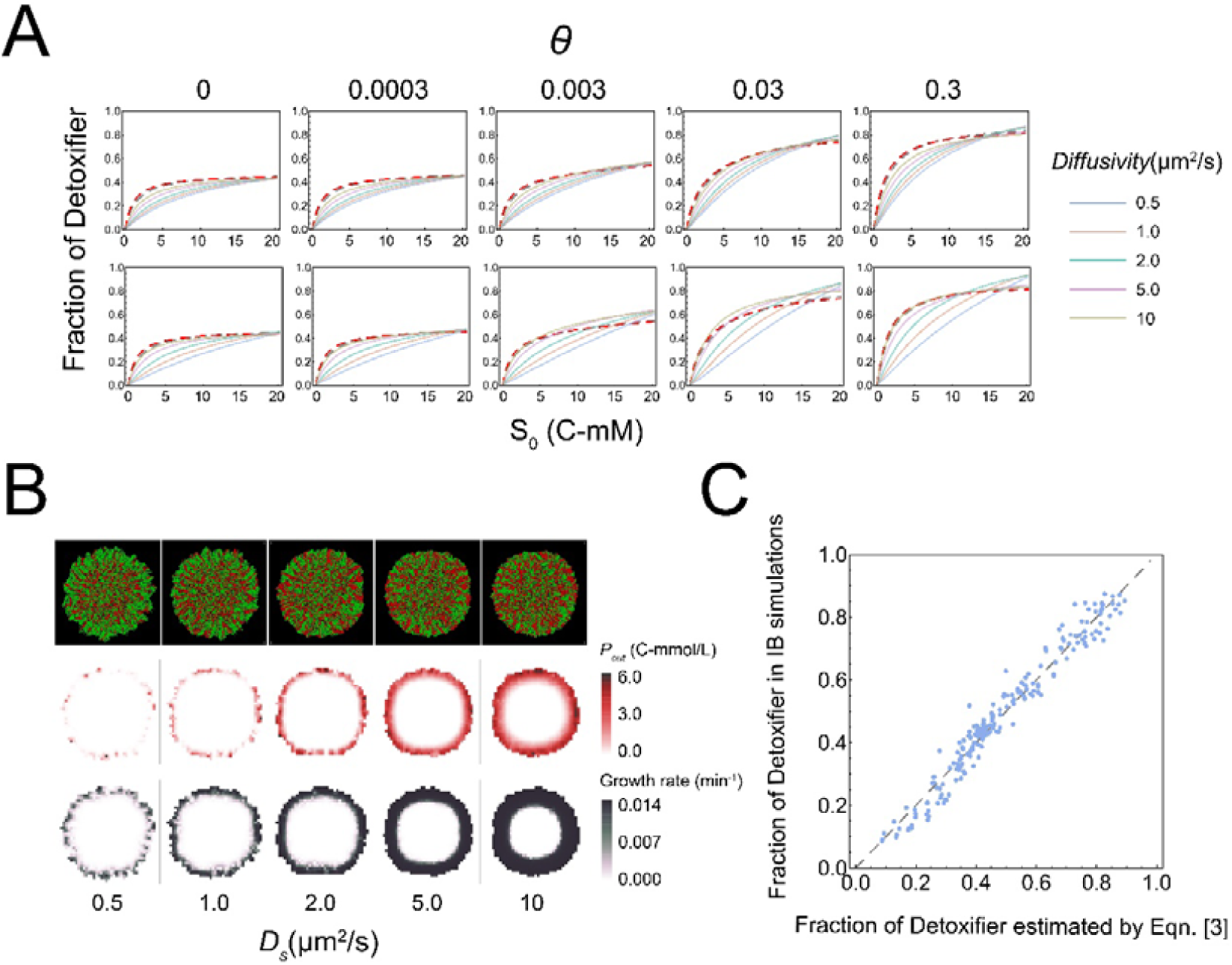
The effects of the diffusion level of substrate, intermediate and product on the structure of MDOL community. (A) The relationship between initial substrate concentration (*S*_*0*_) with the steady-state proportion of Detoxifier cells in the expanding region of the colonies, across different substance diffusion level (denoted by different curve colors) and different strength of substrate toxicity (*θ*, denoted by five subgraphs). First row: diffusion levels of S, I and P (that is *D*_*i*_, *D*_*i*_, and *D*_*p*_) were set to be identical and simultaneously modulated in the simulations. Second row: Diffusion levels of I and P (*D*_*i*_ and *D*_*p*_) were set as default values shown in Supplementary Table 7, while diffusion levels of S were solely modulated. Other parameters in these simulations were initialized with the default values shown in Supplementary Table 7. The simulation data were then fitted to Eqn. [2] to obtain the curves shown in the plot. The Adjust R^2^ values for these fitting analyses range from 0.994 to 0.997. (B) Diffusion levels of substrate affected the thickness of cell ‘active layer’. Representative colony images (first row), the corresponding distributions of final product (second row), as well as the distributions of cell growth rates (third row) in the 2D plane at steady-state, obtained from individual-based simulations initialized with different diffusion level of substrate. Shown are the results in which *S*_*0*_ was set to 10 C-mol/L and *θ* was 0 (not include substrate toxicity). In the colony images, Detoxifier cells are shown in red, while Embezzler cells are shown in green. Thickness of cell ‘active layer’ is reflected by thickness of the cell layer that possessing positive growth rate (third row). (C) The linear correlation between the steady-state frequencies of Detoxifier predicted by Eqn. [4] and those frequencies obtained by our Individual-based simulations. The dashed line shows the linear curve in which the predicting results is completely identical to simulated results. The best fitting value of *ks, Fd*_*max*_, *kt, TS*_*max*_, *kd*_*1*_, and *kd*_*2*_ in this case are 30.8, 0.446, 1.46, 1.05, 14000, and 44.8 respectively.

In this formula, 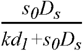 represents an estimate of the positive effect of increasing substrate diffusion level via thickening cell ‘active layer’, related to the initial substrate concentration (*s*_0_ ; Figure 6B; [48]); 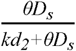 represents an estimate of the negative effect of increasing substrate diffusion level, influenced by toxic strength of the substrate (Figure 6A; the second row). Eqn. [3] accurately estimated the structure of MDOL community in our IB simulations (Figure 6C; R^2^=0.994). Overall, we concluded that the traits of substrate, including concentration, toxicity, and diffusivity, are fundamental to shaping the structure of MDOL community.

## Discussion

Here we show how substrate traits shape the structure of the microbial communities engaged in metabolic division of labor (MDOL) when degrading organic compounds. The population performing the first step is favored by both higher substrate concentration and its toxicity. This rule is applicable when the community grow both in a well-mixed and a spatially structured environment.

Recently, numerous studies have explored the strategy of dividing metabolic roles across different populations in a consortium toward removal of organic pollutants [8, 49-53]. Our proposed rule may be expanded to forecast the structure of these consortia. For instance, one recent study reported that a bacterial consortium composed of *Leucobacter* sp. GP and *Achromobacter denitrificans* PR1 efficiently degrades an antibiotic, sulfamethoxazole, in which the strain GP is responsible for the initial metabolism of the sulfamethoxazole (Detoxifier), and the strain PR1 carries out the subsequent conversion (Embezzler)[12]. This study measured the structures of the community across a gradient of initial substrate concentrations, and found that the proportion of the GP is positively correlated with the initial sulfamethoxazole concentration. This observation largely agrees with the idea derived from our model and experiments. The prediction on the structure of community may largely help to manage these communities for better performance [15, 28, 29].

Our study also indicated that limited mass diffusion in spatially structured environments is one key factor to determine the structure of a community. This finding is reminiscent of recent studies proposing that limited mass diffusion plays significant role on the structure of the communities engaged in other diffusion-based interaction modes, including syntrophic exchange [37, 40, 54], cross-protection [55], and ‘rock-paper-scissors’ interaction [56, 57]. One important hypothesis from these studies is that limited mass diffusion is one possible way to privatize public benefit [37, 40, 58]. We found this hypothesis is also applicable to explain the structuring of the community engaged in MDOL. On the one hand, limited mass diffusion helps the Embezzler population to privatize the final product for its own growth. On the other hand, it helps the Detoxifier population to privatize its benefit from detoxification. Therefore, limited mass diffusion may be a universally used avenue for microorganisms to maintain their private benefit in spatially structured environments. In our IB modelling, we also found that specific spatial patterns developed by the MDOL community. In agreement with previous studies [39, 59, 60], when two populations engaged in MDOL, cells from the two populations are spatially more proximal to each other than the scenario when the two populations did not exhibit defined interactions (Supplementary Figure 27). In addition, we also found that the level of spatial proximity was governed by substrate traits (Supplementary Figure 27). Interestingly, when the strength of substrate toxicity was higher, the Detoxifier cells occupied the periphery of the growing colony, forming a clearly ‘ring’ around the colony (Figure 5; Supplementary video 3; Supplementary Figure 28). The formation of this ring might be due to the fact that the substrate was present at higher concentrations at the colony edge, and hence more toxic, thus largely favoring Detoxifier cells at edge. These results suggest that substrate traits also govern the spatial distributions of different cells in the colony developed by MDOL community, which may in turn, affect the structure of such community. Although we did not observe this featured cell distribution in our experiments, one recent study found that a MDOL community that degrades toluene developed a similar ‘ring’-shape pattern as observed in our IB model [59]. Therefore, such cell distribution may represent a critical feature of the spatial patterns developed by a MDOL community that degrades toxic substrates.

While our study provides critical new insights into how the community engaged in MDOL assembles, a number of limitations need to be taken into consideration. First, our model analysis showed that substrate toxicity is vital to determine the structure of communities engaged in MDOL. However, due to the difficulties in manipulating the toxicity of the substrate (salicylate) *in vitro*, we were unable to experimentally compare the impact of the different toxic strengths on the structure of our community. Nevertheless, our model correctly predicts that simply increasing the initial substrate concentration is unlikely to shape a community dominated by the Detoxifier population, while the presence of substrate toxicity renders the ‘Detoxifier’ population in the community to become dominant. Therefore, the observation that Detoxifier population was able to dominate the synthetic consortium when supplying high concentration of salicylate, and the measured biotoxicity of salicylate strongly suggested that substrate toxicity should affect the structure of our synthetic microbial consortium. In agreement with this idea, our prediction functions involved in salicylate toxic strength fits the experiment results very well. To further examine this idea, it is necessary to design a better system in which the toxicity of the substrate can be modulated.

Second, our ODE model suggests that apart from substrate traits, five other key parameters exist that exhibit considerable effects on the structure of a MDOL community. Here, we primarily focused on the effects of substrate traits, without analyzing in detail how all the seven key factors collectively determine the structure of community. Nonetheless, our analysis presented here suggests that biotic factors such as speed of the first reaction (*α*_*1*_), mass transport rate (*γ*_*s*_, *γ*_*i*_, *γ*_*p*_), as well as consumption rate of P (*Cp*), affected the structure of the community, namely by determining the value of parameters in Eqn. [2] (i.e., *Fd*_*max*_, *ks, Ts*_*max*_, and *kt*). However, due to the difficulties in analytically solving non-linear ODEs, as well as the low efficiency of individual-based simulations [61], detailed quantitative understanding of how all these factors affect the structure of MDOL community remains limited. Further studies may use more simplified models that combine these elements to provide a more general description of the principles governing the structuring of a MDOL community.

To engineer stable and high-efficient microbial systems for bioproduction or biodegradation, it will be critical to predict how the communities assembled by a given set of strains exhibiting modularized functions. Our results demonstrate that, for a given community engaged in MDOL, its structure can be quantitatively estimated from the abiotic factors, such as the traits of its substrate, suggesting that it is feasible to manage microbial communities through manipulation of specific environmental factors, to address grand challenges facing human society in agriculture, degradation of the environment, and human health.

## Methods

### Formulation and analyses of the ODE model

#### Formulation of the ODE model

To simulate the dynamics of a MDOL community in well-mixed system, a mathematical model was formulated using ordinary differential equations (ODEs). Here, the dimensionless forms of the models were presented. The detailed derivations of the models, and choices of parameter values are described in Supplementary Information S1.

As described in the Results section, a two-step pathway was assumed to be implemented by MDOL between two populations (Figure 2A and Figure 3A). For simplicity, the basic model was built based on five simple assumptions: (1) The systems are well mixed in each compartment (inside a cell or in the extracellular space). (2) transport of substrate (S), intermediate (I) and final product (P) is mediated by passive diffusion; (3) P was assumed to be the sole and limited resource for the growth of the two populations and its consumption was calculated following Monod equations; (4) Basic biological properties (the coefficients in Monod equations) regarding the growth of the two populations are identical, since we only focused on the effects of abiotic factors; (5) when applicable, substrate toxicity was introduced by adding three different toxic terms to the growth equation (Supplementary Table 3), dependent on intracellular S concentration of the corresponding population. The dynamics of intracellular and extracellular I and P are given by

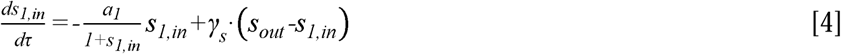

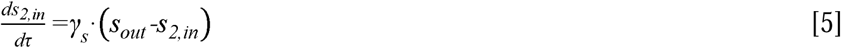

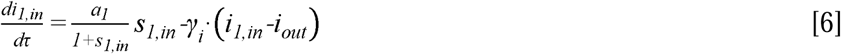

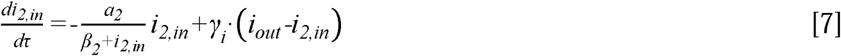

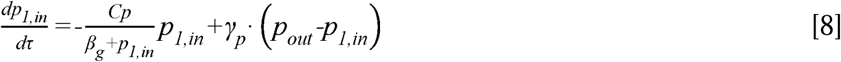

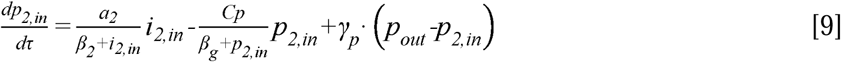

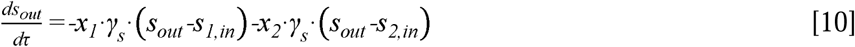

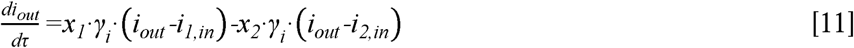

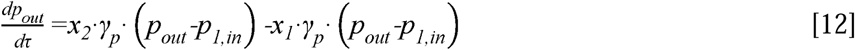

The growth of the two populations was modeled using a general logistic function with first-order cell death:

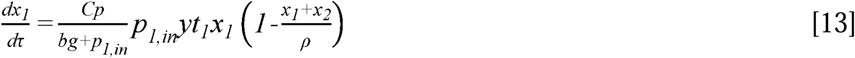

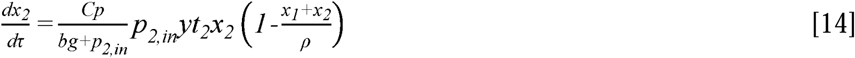

The definitions and dimensionless methods of all variables are listed in Supplementary Table 1. The definitions and dimensionless methods, as well as the value ranges of all the parameters involved in these equations are listed in Supplementary Table 2.

#### Simulation and analyzing protocol of the ODE model

Details of the simulation and analysis protocols of our ODE model and the downstream analyses are described in Supplementary Information S1.3. Briefly, to solve the community dynamics of the MDOL community with given parameter sets, numerical simulations of our ODE model were performed using *NDsolve* function of *Wolfram Mathematica*. The numerical solutions of all the variables, including the dynamics of mass (S, I, P) concentration and biomass, were recorded for further analyses. To perform simulations with numerous parameter sets, as well as the downstream analysis, custom *Mathematica* scripts were wrote mainly based on the *Do* loop function.

### Individual-based modeling

Our individual based (IB) model was constructed based on *gro* platform (https: https://github.com/liaupm/GRO-LIA), a simulator designed by Gutiérrez and colleagues aiming to describe multicellular bacterial behavior [62]. The model aims to simulate the growth of a microbial colony composed of two populations who execute substrate degradation via MDOL on a surface. The model was formulated mainly using the same equations as our dimensional ODE model (Supplementary Information S1.1, Eqns. [S1]-[S13]) to characterize the intra- and extracellular dynamics of mass (S, I, P) concentration, as well as to calculate the rate of cell growth. Four main differences exist between our IB model and the ODE model: (1) The IB model was formulated on a spatially structured surface, and the diffusion of S, I, and P was limited; (2) Mass dynamics was modelled at single-cell level; (3) The growth of both populations was modelled at single-cell level, and passive cell shoving during the cell growth was included; (4) cells were inoculated in the center of the surface, and the entire community underwent ‘colony range expansion’, a process whereby the community immigrate outwards as a whole, driven by the force generated from cell growth and division (Supplementary Figure 15). The mathematical framework formulating these four points is described in Supplementary Information S2.1. To implement our design of the IB model, custom codes were written in *gro* language. Variables and Parameters in the IB model are summarized in Supplementary Table 7. Details of the IB simulation workflow are described in Supplementary Information S2.

### Experimental verification of our model prediction

#### Genetic manipulation of the P. stutzeri strains

All *P. stutzeri* strains were engineered from a naphthalene-degrading bacterial strain *P. stutzeri* AN10 [63]. Genes that encode the key enzymes responsible for corresponding metabolic steps in salicylate degradation pathway were knocked out to generate the *P. stutzeri* strains. The details of the genetic manipulation of are described in Supplementary information S3.

#### Liquid cultivation of our synthetic microbial communities

Liquid cultivation of our synthetic microbial communities was performed in 96-well plates that contains 120 μL fresh minimum medium. Proportions of the two populations in the community were estimated by measuring the fluorescent intensity of the two strains involved using a microplate reader (Molecular Devices, Sunnyvale, America). Detailed protocols are described in Supplementary information S4.

#### Colony pattern formation assays

Colony pattern formation assays were performed on the agarose surface in a Petri dish (60 mm in diameter). Images of the colony patterns were taken under a 5× objective using a Leica DM6000B fluorescence microscope (Leica Corporation, Wetzlar, Germany) equipped with a LED fluorescence illuminator (Leica Corporation). The relative fraction of each population in the colonies was measured by image analysis, as well as similar fluorescence-measurement method as performed in liquid cultivation experiments. Detailed protocols are described in Supplementary information S5.

### Statistical analysis

Unless indicated otherwise, the number of replicates was three for each simulation, and six for each experiment. For comparative statistics, unpaired, two-tailed, Student’s t-test was performed in Wolfram Mathematica (version 12.4). To fit the data to the proposed function, Nonlinearmodelfit function of the Wolfram Mathematica (version 12.4) was applied.

## Code availability

All custom *Mathematica* codes used for ODE simulation and data analyses, as well as the source *gro* codes used for our IB simulations are available at Github: https://github.com/RoyWang1991/MDOLcode/tree/master/MDOL-spatial.

## Competing Interests

The authors declare that they have no conflict of interest.

## Acknowledgments

We wish to thank Professor Ping Xu (Shanghai Jiao Tong University, Shanghai, P.R. China) for supplying plasmid pMMPc-Gm, used for fluorescence labeling in this study; Dr. Min Lin (Chinese Academy of Agricultural Sciences, Beijing, P.R. China) for providing plasmid pK18mobsacB and pRK2013, used for genetic engineering in this work; Professor Martin Ackermann (ETH Zurich, Zurich, Switzerland), Dr. David Johnson (Eawag, Dübendorf, Switzerland) and Yinyin Ma (Eawag, Dübendorf, Switzerland) for constructive inputs on the design of this study; Professor Martín Gutiérrez (Universidad Politécnica de Madrid, Madrid, Spain) for his kindly guidance for the set-up of the *gro* platform for the individual-based simulations; Dr. T. Juelich (UCAS, Beijing) for linguistic assistance during the preparation of this manuscript. This work was supported by National Key R&D Program of China (2018YFA0902100 and 2018YFA0902103), and National Natural Science Foundation of China (91951204, 31761133006, 31770120, and 31770118).

## Notes

### Competing Interest Statement

The authors have declared no competing interest.

