## Supplementary Information for "Substrate traits shape the structure of microbial community engaged in metabolic division of labor"

### **S1 Formulation, simulation and analyses of our ODE model**

#### **S1.1 Formulation of our ODE model**

As described in the main text, the basic model was constructed to describe the dynamics of a microbial community of two populations that degraded a substrate (S) through metabolic division of labor (MDOL). The first population was named as ‘Detoxifier’, who was only capable of transforming potentially toxic S to an intermediate (I). The second population was named as ‘Embezzler’, who was able to convert transforming I to final product (P). The model was formulated based on five biologically relevant assumptions, as follow

(1) All the substances (S, I and P) in the system are well-mixed in each compartment (inside a cell or in the extracellular space).

(2) transport of S, I and P is mediated by passive diffusion;

(3) P was assumed to be the sole and limited resource for the growth of the two populations and its consumption was calculated following Monod equations;

(4) Basic biological properties (the coefficients in Monod equations) regarding the growth of the two populations are same, since we only focus the effects of abiotic factors;

(5) when applicable, substrate toxicity was introduced by adding three different toxic terms to the growth equation (Supplementary Table 3), depending on intracellular S concentration of the corresponding population.

Based on these assumptions, an ordinary differential equation (ODE) system regarding the mass dynamics was first built:

$\frac{\text{d}\text{S}_{\text{1,in}}}{\text{dt}}\text{=-}\frac{\text{k}_{\text{1}}\text{E}_{\text{1}}}{\text{K}_{\text{1}}\text{+}\text{S}_{\text{1,in}}}\text{S}_{\text{1,in}}\text{+}\text{r}_{\text{S}}\text{∙}\left( \text{S}_{\text{out}}\text{-}\text{S}_{\text{1,in}} \right)$	[S1]

$\frac{\text{d}\text{S}_{\text{2,in}}}{\text{dt}}\text{=}\text{r}_{\text{S}}\text{∙}\left( \text{S}_{\text{out}}\text{-}\text{S}_{\text{2,in}} \right)$	[S2]

$\frac{\text{d}\text{I}_{\text{1,in}}}{\text{dt}}\text{=}\frac{\text{k}_{\text{1}}\text{E}_{\text{1}}}{\text{K}_{\text{1}}\text{+}\text{S}_{\text{1,in}}}\text{S}_{\text{1,in}}\text{-}\text{r}_{\text{I}}\text{∙}\left( \text{I}_{\text{1,in}}\text{-}\text{I}_{\text{out}} \right)$	[S3]

$\frac{\text{d}\text{I}_{\text{2,in}}}{\text{dt}}\text{=-}\frac{\text{k}_{\text{2}}\text{E}_{\text{2}}}{\text{K}_{\text{2}}\text{+}\text{I}_{\text{2,in}}}\text{I}_{\text{2,in}}\text{+}\text{r}_{\text{I}}\text{∙}\left( \text{I}_{\text{out}}\text{-}\text{I}_{\text{2,in}} \right)$	[S4]

$\frac{\text{d}\text{P}_{\text{1,in}}}{\text{dt}}\text{=-}\frac{\text{kg}}{\text{K}_{\text{g}}\text{+}\text{P}_{\text{1,in}}}\text{ }\text{P}_{\text{1,in}}\text{+}\text{r}_{\text{P}}\text{∙}\left( \text{P}_{\text{out}}\text{-}\text{P}_{\text{1,in}} \right)$	[S5]

$\frac{\text{d}\text{P}_{\text{2,in}}}{\text{dt}}\text{=}\frac{\text{k}_{\text{2}}\text{E}_{\text{2}}}{\text{K}_{\text{2}}\text{+}\text{I}_{\text{2,in}}}\text{I}_{\text{2,in}}\text{-}\frac{\text{kg}}{\text{K}_{\text{g}}\text{+}\text{P}_{\text{2,in}}}\text{ }\text{P}_{\text{2,in}}\text{+}\text{r}_{\text{P}}\text{∙}\left( \text{P}_{\text{out}}\text{-}\text{P}_{\text{2,in}} \right)$	[S6]

$\frac{\text{d}\text{S}_{\text{out}}}{\text{dt}}\text{=-}\frac{\text{X}_{\text{1}}}{\text{V}_{\text{E}}}\text{∙}\text{r}_{\text{S}}\text{∙}\left( \text{S}_{\text{out}}\text{-}\text{S}_{\text{1,in}} \right)\text{-}\frac{\text{X}_{\text{2}}}{\text{V}_{\text{E}}}\text{∙}\text{V}_{\text{c}}\text{∙}\text{r}_{\text{S}}\text{∙}\left( \text{S}_{\text{out}}\text{-}\text{S}_{\text{2,in}} \right)$	[S7]

$\frac{\text{d}\text{I}_{\text{out}}}{\text{dt}}\text{=}\frac{\text{X}_{\text{1}}}{\text{V}_{\text{E}}}\text{∙}\text{r}_{\text{I}}\text{∙}\left( \text{I}_{\text{out}}\text{-}\text{I}_{\text{1,in}} \right)\text{-}\frac{\text{X}_{\text{2}}}{\text{V}_{\text{E}}}\text{∙}\text{r}_{\text{I}}\text{∙}\left( \text{I}_{\text{out}}\text{-}\text{I}_{\text{2,in}} \right)$	[S8]

$\frac{\text{d}\text{P}_{\text{out}}}{\text{dt}}\text{=}\frac{\text{X}_{\text{1}}}{\text{V}_{\text{E}}}\text{∙}\text{r}_{\text{P}}\text{∙}\left( \text{P}_{\text{out}}\text{-}\text{P}_{\text{1,in}} \right)\text{-}\frac{\text{X}_{\text{2}}}{\text{V}_{\text{E}}}\text{∙}\text{r}_{\text{P}}\text{∙}\left( \text{P}_{\text{out}}\text{-}\text{P}_{\text{2,in}} \right)$	[S9]

To facilitate modeling analysis, the equations were non-dimensionalized following the method described in Supplementary Table 1 and Supplementary Table 2, which formulated the model (Eqns. [4]-[12]) in the Methods section of the main text.

The growth dynamics of the two populations was modelled using a general logistic function modified with toxic terms ($\text{T}_{\text{1}}$, $\text{T}_{2}$)

$\frac{\text{d}\text{X}_{\text{1}}}{\text{dt}}\text{=}\text{g}_{\text{D}}\text{X}_{\text{1}}\text{T}_{\text{1}}\left( \text{1-}\frac{\text{X}_{\text{1}}\text{+}\text{X}_{\text{2}}}{\text{N}_{\text{m}}} \right)$	[S10]

$\frac{\text{d}\text{X}_{\text{2}}}{\text{dt}}\text{=}\text{g}_{\text{E}}\text{X}_{\text{2}}\text{T}_{\text{2}}\left( \text{1-}\frac{\text{X}_{\text{1}}\text{+}\text{X}_{\text{2}}}{\text{N}_{\text{m}}} \right)$	[S11]

Note that here $\text{X}_{\text{1}}$ and $\text{X}_{\text{2}}$, represents the total cell volume of the two populations. For a population with sufficiently large cell number, the total biomass is proportional to the total cell volume. Thus, the population size above can be interpreted as the total cell volume, which is then use as the basis to derive the dimensionless models. Three forms of $\text{T}_{\text{1}}$, $\text{T}_{2}$ were considered, which is listed in Supplementary Table 3.

As described in assumption (3), both of the two populations were assumed to use the final product as the sole carbon source, thus the growth rate of the two populations, $\text{g}_{D}$ and $\text{g}_{E}$, are calculated via a Monod equation limited by the intracellular concentration of P, so that,

$\text{g}_{\text{D}}\text{=}\frac{\text{kg}_{\text{1}}\text{P}_{\text{1,in}}}{\text{Kg}_{\text{1}}\text{+}\text{P}_{\text{1,in}}}\text{Y}_{\text{1}}\text{c}_{\text{1}}$	[S12]

$\text{g}_{\text{E}}\text{=}\frac{\text{kg}_{\text{2}}\text{P}_{\text{2,in}}}{\text{Kg}_{\text{2}}\text{+}\text{P}_{\text{2,in}}}\text{Y}_{\text{2}}\text{c}_{\text{2}}$	[S13]

Then Eqn [S12] and [S13] were substituted into Eqn [S10] and [S11], which were then non-dimensionalized following the method described in Supplementary Table 1 and Supplementary Table 2, so the dimensionless cell growth equations were obtained (Eqns. 13-14 in the Methods section of the main text.).

#### **S1.2 Definitions and values of dimensionless variables and parameters**

The definition of all the variables, their corresponding dimensionless counterparts, and their value range during the numerical simulations are listed in Supplementary Table 1. In addition, the definition of all the parameters and their corresponding dimensionless methods (when applicable), are listed in Supplementary Table 2. Parameter values are chosen/calculated according to estimated values from literature, and are listed in Supplementary Table 2. Dimensionless (ND) values are calculated based on methods in Supplementary Table 2. Parameter ranges are chosen to incorporate at least ~10-fold higher or lower than the estimated value.

#### **S1.3 Analyses to identify key parameters that affect the assembly of MDOL community**

To test whether and how the parameters included in the ODE model affect community assembly, two rounds of simulations were performed. In the first round, numerical simulations were performed using 885,735 (not considering substrate toxicity) and 7,971,615 (considering substrate toxicity) parameter sets that contained three gradient values for each parameter, as well as three different forms of terms to characterize how substrate toxicity affects population growth [1-3]. The steady-state fractions of Detoxifier in these simulations were then collected. Values of the parameter and the steady-state fraction were then normalized and subjected to multivariate regression analyses using *LinearModelFit* function of *Wolfram* *Mathematica*. The parameter that significantly (p < 0.01) and largely (fitting coefficient over 0.01) affect the community assembly were identified as the key parameters (Figure 3; Supplementary Figure 3; Supplementary Table 4).

The second round of simulations were designed to further test how substrate concentration and toxicity affect the assembly of MDOL communities. Numerical simulations were performed using 163,296 (not considering substrate toxicity) and 489,888 (considering substrate toxicity) parameter sets that contained 18 gradient values for initial substrate concentration (*s_0_*), 21 gradient values for toxic strength of substrate (*θ*), six gradient values for each of other five key parameters, as well as the three forms of toxic terms when applicable. The values of other four parameters (*y*, *bg*, *a2*, *ρ*) were set as the default values shown in Supplementary Table 2. The simulation results were classified into 7776 groups. Each group possessed same values of the other five key parameters, but 18 varied values of *s_0_* and 21 varied values of *θ* (when substrate toxicity is included). Then the results were fitted to Eqn. [1] (not considering substrate toxicity) or Eqn. [2] (considering substrate toxicity) using *NonLinearModelFit* function of *Wolfram* *Mathematica*. Adjusted R^2^ and significance of each fitting parameters were analyzed. Furthermore, the best fitting values of $\text{Fd}_{\text{max}}$, $\text{ks}$, $\text{Ts}_{\text{max}}$, and $\text{kt}$ were grouped with the corresponding parameter sets, and subjected to multivariate regression analyses using the same protocol as above (Supplementary Figure 4-5; Supplementary Figure 7-10; Supplementary Table 5). In addition, the relationships between the parameters in Eqn. [1] or Eqn. [2] ($\text{Fd}_{\text{max}}$, $\text{ks}$, $\text{Ts}_{\text{max}}$, and $\text{kt}$) and the seven five key parameters in our ODE system ($\text{Cp}$, $a_{1}$, $\text{γ}_{\text{s}}$,$\text{γ}_{\text{i}}$,$\text{γ}_{\text{p}}$) were also built. As shown in Supplementary Figure 4 and Supplementary Figure 5, $\text{Fd}_{\text{max}}$ was strongly negatively correlated with the consuming rate of the final product ($\text{Cp}$), while the value of $\text{ks}$ was largely affected by the consuming rate of the final product ($\text{Cp}$), transport rate of S ($\text{γ}_{\text{s}}$), I ($\text{γ}_{\text{i}}$) and P ($\text{γ}_{\text{p}}$). When substrate toxicity is included, the parameters affecting the value of $\text{Fd}_{\text{max}}$ and $\text{ks}$ were similar as the non-toxic scenario (Supplementary Figure 7-8); the value of $\text{kt}$ were affected by $\text{Cp}$, $\text{γ}_{\text{p}}$, and $\text{a}_{\text{1}}$ (Supplementary Figure 9). the values of $\text{Ts}_{\text{max}}$ was largely determined by $\text{Cp}$, $\text{γ}_{\text{s}}$, as well as the rate of the first reaction ($\text{a}_{\text{1}}$) (Supplementary Figure 10);

Furthermore, to perform mathematical prediction regarding the assembly of our synthetic communities, exact parameter values were taken from experimental measurements or previous reported (as listed in Supplementary Table 6). Mathematical simulations were performed and the data were fitted to Eqn. [2], thus predicting function listed in Figure 4 was obtained. All the *Mathematica* scripts used for simulations, analyses, and data visualization are available in Github: <https://github.com/RoyWang1991/MDOLcode/tree/master/MDOL-spatial>.

### **S2 Formulation, simulation and analyses of our Individual-based model**

#### **S2.1 Formulation of the Individual-based model**

As we mentioned in Methods section, the formulation of our Individual-based (IB) was similar as our ODE model, but possess four main differences. Here, we described detailedly how these differences were formulated in the IB model.

##### *S2.1.1 Assumptions of a spatially structured environment*

The IB model was formulated on a spatially structured 2D plane with a size of 160 μm × 160 μm, in which the mass diffusion was limited. To facilitate the simulation of mass diffusion, the space was divided into 1600 grid units with a size of 4 μm × 4 μm. Then the diffusion of S, I and P was modelled by finite element method built in *gro*, as well as other similar studies [4-7], which assumes that the mass diffusion basically follows Fick Law. Therefore, the extracellular concentration of substrate or a metabolite, in the grid coordinate ($\text{i}$, *j*), $\text{C}_{\text{i,j}}$, is updated at each step via

Δ$\text{C}_{\text{i,j}}$= -6⸱$D$⸱$\text{C}_{\text{i,j}}$+$D$⸱(0.5⸱$\text{C}_{\text{i+1,j-1}}$+$\text{C}_{\text{i+1,j}}$+0.5⸱$\text{C}_{\text{i+1,j+1}}$+$\text{C}_{\text{i+1,j}\text{-}\text{1}}$+$\text{C}_{\text{i,j+1}}$+ 0.5⸱$\text{C}_{\text{i}\text{-}\text{1,j-1}}$ +$\text{C}_{\text{i-1,j}}$+ 0.5⸱$\text{C}_{\text{i}\text{-}\text{1,j+1}}$)	[S14]

Here, $D$ is the diffusion rate of the substance. Therefore, the diffusion rate of S, I and P was mediated by their diffusion coefficients, $\text{D}_{\text{s}}$, $\text{D}_{\text{i}}$, and $\text{D}_{\text{p}}$.

##### *S2.1.2 Assumptions on cell growth and passive cell shoving*

To model cell growth at individual level, each cell in IB model were characterized as a rod-shaped capsule, as most of the previous studies [8-11]. Biological traits of the *ith* cell were represented by a vector containing the related variables, including cell length ($\text{L}_{\text{D,i}}$ and $\text{L}_{\text{E,i}}$; ‘D’ means ‘Detoxifier’ and ‘E’ means ‘Embezzler’, indicating which population the cells belong to), its orientation ($\varphi_{i}$) and its position coordinates. The values of these variables were updated using time-resolved discrete simulation. Growth of cells was modelled by calculating their elongation (that is, the increase of $\text{L}_{\text{D,i}}$ or $\text{L}_{\text{E,i}}$), as follow:

$$\frac{\text{d}\text{L}_{\text{D,i}}}{\text{dt}}\text{=}\text{g}_{\text{D,i}}\text{L}_{\text{D,i}}$$

$$\frac{\text{d}\text{L}_{\text{E,i}}}{\text{dt}}\text{=}\text{g}_{\text{E,i}}\text{L}_{\text{E,i}}$$

Since the model assumes that the radius of cell remains unchanged, the volume of a cell is proportional to its length. Thus, the growth of cell volume is directly related to the intracellular P concentration, which is consistent with our ODE model. The growth rate ($\text{g}_{\text{D,i}}$ and $\text{g}_{\text{E,i}}$) of each cell was calculated using the same quantitative framework as defined in Eqn. [S10]-[S13], except that the intracellular P concentration of the *ith* cell was used as the argument to introduce the differences in individuality. Cells were initiated with a length of $\text{L}_{\text{0}}\text{+}\text{ε}_{\text{1}}$, in which $\text{ε}_{\text{1}}$is the random noise of initial cell length of a seeding population, following a uniform distribution. When the volume of a cell reached the division length $\text{L}_{\text{d}}\text{+}\text{ε}_{\text{2}}$, it divided via binary fission. $\text{ε}_{\text{2}}$ is the random noise in the cell cycle, and also follows a uniform distribution. In addition, although our IB model did not include active cell movement in the model, passive cell shoving derived from cell-cell contact was included, and modelled by the ‘collision detection and collision response’ strategy built-in *gro* [12, 13].

##### *S2.1.3 Assumptions on mass dynamics*

To the intra- and extracellular mass dynamics at single-cell level, the intracellular mass concentration of the *ith* cell, $\text{S}_{\text{D,in}\text{,}\text{i}}$, $\text{S}_{\text{E,in}\text{,}\text{i}}$, $\text{I}_{\text{D,in}\text{,}\text{i}}$, $\text{I}_{\text{E,in}\text{,}\text{i}}$ $\text{P}_{\text{D,in}\text{,}\text{i}}$, and $\text{P}_{\text{E,in}\text{,}\text{i}}$, were also represented by a vector containing these variables. Similar as above, the values of these variables were updated using time-resolved discrete simulation. In each step, the dynamics of intracellular mass concentration of a cell was simulated using same quantitative framework defined in as Eqn. [S1]-[S6], except that the calculations were performed at individual level. In addition, the heterogenous distribution of extracellular mass concentration was modelled using the finite element method mentioned in S2.1.1. Notably, the extracellular substance concentration of a cell was actually represented by the concentration of the substance in the 4 μm × 4 μm grid unit containing the corresponding cell. Transport (absorbing or emitting) of substance (S, I and P) of all the cells in one grid unit was modelled using Eqn. [S7]-[S9], but calculated at individual level. Initially, S were evenly distributed across the plane with a concentration of *S_0_*, while the initial concentrations of I and P, were set to zero.

##### *S2.1.4 Simulating colony range expansion*

Inocula consisting of a 1:1 mix (2000-cell) of Detoxifier and Embezzler cells, were randomly scattered and oriented within a 300-μm circle to start colony growth. Simulations were set to terminate once the cell population exceeded 6,000 individuals, during which colony range expansion of a MDOL community was simulated (Supplementary Figure 15).

#### **S2.2 The effects of substance diffusivity on the structure of the MDOL community**

To test whether the diffusivity of S, I and P affects the structure of the MDOL community, we simultaneously modulated the diffusion level of S, I, and P (changing $\text{D}_{\text{s}}$, $\text{D}_{\text{i}}$, and $\text{D}_{\text{p}}$). We also constructed a ‘well-mixed’ IB model by changing assumptions defined in S2.1.1, in which S, I and P were assumed to be evenly mixed during the simulations. In this model, S, I and P were rapidly and evenly distributed across the entire space at each time step. To fulfill this setting, at the beginning of each time step, the amount of S, I and P were summed up and equally spread among 2D plane (the 1600 grid units). Therefore, there is no difference in extracellular concentration of S, I and P among all the cells at each time step. This modification was realized by changing the original settings of mass diffusion in our *gro* code.

As shown in Fig. 6A, the prediction functions derived from the scenarios with higher level of diffusivity, as well as the ‘well-mixed’ IB model, are more similar to the function derived from ‘well-mixed’ ODE model. This result confirmed that limited mass diffusion plays a significant role in shaping the structure of the MDOL community.

How the diffusion level affects the structure of MDOL community? Firstly, limited diffusion of P promoted the product privatization by the Embezzler cells, since product was distributed closer to its producers when its diffusion is limited (Supplementary Figure 22). From this perspective, limited diffusion of P favors the Embezzler. We tested this hypothesis by solely modulating the diffusion level of P (solely changing $\text{D}_{\text{p}}$) in our IB model. As shown in Supplementary Figure 23, we found that higher level of P diffusion indeed favors the Detoxifier. Secondly, Embezzler cells require intermediate metabolite I to produce final product P for growth, and I was largely distributed around the Detoxifier cells (Supplementary Figure 24). Thus, when I diffused at a lower speed, more Detoxifier cells are required to fulfill the supply of I. Therefore, we hypothesized that limited diffusion of I favored the growth of Detoxifier. As shown in Supplementary Figure 25, with an increase of $\text{D}_{\text{i}}$, I was distributed more around the Embezzler cells, and the relative proportion of Detoxifier cells decreases.

Finally, we investigated how substrate diffusivity affects the structure of MDOL community by solely changing diffusion levels of the substrate ($\text{D}_{\text{s}}$) in our IB simulations. As shown Figure 6A, when substrate toxicity is omitted, the proportion of Detoxifier cells increased with an increase of diffusion levels of the substrate. This observation is due to the fact that higher diffusion levels of S thicken the cell’s ‘active layer’, thus increasing production and secretion of the final product by Embezzler cells (Figure 6B; [14]). However, limited diffusion of S also helps to maintain a lower local extracellular S concentration around the Detoxifier cells (Supplementary Figure 26). When substrate toxicity was considered, this local gradient caused the substrate toxicity to be less harmful to the Detoxifier cells. Thus, in addition to thickening cell ‘active layer’ that benefits Detoxifier cells, higher diffusion level of S also decreases the fitness of the Detoxifier cells by changing relative toxic level of S. Combining these two effects, we hypothesized that with the increase of substrate toxicity and its concentration, Detoxifier should gradually turn to be favored in an intermediate diffusion level of substrate. This hypothesis was successfully verified by our IB simulations (Figure 6A; second row). Overall, substance diffusivity also affects the structure of MDOL community in spatially structured environment.

#### **S2.3 Model parameterization**

Basically, the measured or previous reported values of parameter (Supplementary Table 6) that match with our experimental system were used to perform IB simulations. In addition, the values of the additional parameters regarding the mass diffusion, as well as characteristics of bacterial cells were obtained from literature, as listed in Supplementary Table 7. In particular, to mainly focus on the effects of substrate traits, the value of initial substrate concentration *S_0_*, was varied from 2.5 Cmmol/L to 20 Cmmo/L, which is consistent with our experimental design; the value of toxic strength of substrate, *θ*, was varied from 10^-4^ to 10^-0.5^, around the scale of our measured toxic strength of salicylate. In addition, sensitive analyses of three key parameters, *kg,* $\text{E}_{\text{1}}$, and $\text{r}_{\text{s}}$ were performed to test their effects on the best fitting of parameters in Eqn. [2] (i.e., $\text{Fd}_{\text{max}}$, $\text{ks}$, $\text{Ts}_{\text{max}}$, and $\text{kt}$; Supplementary Figure 18; Supplementary Figure 20).

#### **S2.4 Simulation protocols and data analyses**

IB model simulations were run on a Cloud server (https://www.yisu.com) running Windows Server 2019 system using *gro* platform that is available online (https: https://github.com/liaupm/GRO-LIA). To perform simulations with numerous parameter sets and replicates, a custom *Mathematica* script were designed to control automatic running of *gro*, mainly based on previously developed *RobotTools* package of *Mathematica*. Unless indicated otherwise, three replicates were performed for each condition and the unconcerning parameters were assigned with the default values listed in Supplementary Table 7. During simulations, images of simulated colony pattern at each time point were simply generated from *gro* using the built-in function ‘*Snapshot*’. Custom functions were included in the *gro* codes to record the position coordinates of every cell, intra- and extra- concentrations of the masses, as well as the cell number of each population at each time point. These simulation data were analyzed and visualized using custom *Mathematica* scripts (version 12.0). Specifically, the maps characterizing extracellular distribution of the masses (Supplementary video 1-4) were first generated using *ArrayPlot* function of *Mathematica*. These maps were then merged with the time-series image sequences of colony exported from *gro.* Finally, the videos were created by exporting these images as the video format using *Mathematica*. To separately analyze the community structures in different colony region, cells were divided into two groups according to their distance to the colony center, in which the cells with distance over 300 μm were grouped as the expanding-region cells and the rest were grouped as the inoculating-region cells. This analysis was also performed using a custom *Mathematica* script. Then the community structures were analyzed and fitted to Eqn. [2] or Eqn. [3] using the same methods as described in ODE model. The source code used for these analyses are available on Github:

<https://github.com/RoyWang1991/MDOLcode/tree/master/MDOL-spatial>.

### **S3 Genetic manipulation of the *Pseudomonas stutzeri* strains involved in our synthetic microbial community**

#### **S3.1 Construction of the *P. stutzeri* strains**

All *P. stutzeri* strains were engineered from a naphthalene-degrading bacterial strain *P. stutzeri* AN10[15]. *P. stutzeri* AN0011 is a derived strain that can degrade salicylate autonomously, and all the genes encoding the corresponding enzymes were located in an operon induced by IPTG (unpublish data). To generate *P. stutzeri* AN0010 and *P. stutzeri* AN0001, *nahG* or *nahH* gene of *P. stutzeri* AN0011 was knocked out, respectively. To generate the pilus mutants, *pilA* and *pilB* genes were simultaneously removed from the host strains, respectively. The genetic manipulation was implemented by allele exchange using the suicide plasmid pK18mobsacB [16, 17]. The constructed strains were validated by PCR and DNA sequencing.

#### **S3.2 Identification of the *P. stutzeri* strains**

To test whether the phenotypes of the strains were consistent with our initial design, two experiments were performed. Firstly, enzymic activity assays were performed to identify whether the strains can execute the defined metabolic step, following the methods reported before ([18] for salicylate 1-hydroxylase and [19] for catechol 2,3-dioxygenase). As shown in Supplementary Figure 29A, cell extract of strain *P. stutzeri* AN0010 only exhibited high enzymic activity of salicylate 1-hydroxylase, while that of strain *P. stutzeri* AN0001 only showed activity of catechol 2,3-dioxygenase. Secondly, the strains were monocultured in minimum medium using salicylate (S) or catechol (I) as sole carbon source. As shown in Supplementary Figure 29B, neither of the Detoxifier and Embezzler strains grew autonomously using salicylate as the sole carbon source, but Embezzler was capable of growing using catechol as the sole carbon source. These results indicate that the phenotypes of the strains follow our initial design.

#### **S3.3 Fluorescence labelling of the *P. stutzeri* strains**

Finally, to measure the community structure of our synthetic community, the strains were labelled with fluorescence. To this end, *mCherry* or *eGFP* was cloned into a constitutive vector, pMMPc-Gm[20], and delivered to the host cells via triparental filter mating [17].

#### **S3.4 Measurement of salicylate toxicity to the *P. stutzeri* strains**

The exact toxic strength of salicylate was measured by culturing the corresponding strains in minimum medium by supplying sufficient amount of final product pyruvate (34 mM, which means adding more pyruvate will no longer increase the bacterial growth rate) and varied amount of toxic salicylate. The experiments were performed in 96-well plates, and each well contained 120 μL liquid culture media. Growth curves were then estimated for every salicylate concentration using a microplate reader (Molecular Devices, Sunnyvale, America), and the data were fitted to the Logistic equation to calculate the specific growth rate $\text{g}_{\text{s}}$.

$$\frac{\text{dN}}{\text{dt}}\text{=}\text{g}_{\text{s}}\text{N(1-}\frac{\text{N}}{\text{K}}\text{)}$$

Here, *N* represents the biomass of the population; $\text{K}$ represents the biomass of the population. Both *N* and *K* are quantified by OD_600_. Subsequently, the obtained growth rate, as well as salicylate concentration data, was fitted to the three equations listed in Supplementary Table 3 to build the relationship between growth rate and salicylate concentration. As shown in Supplementary Figure 13, the reciprocal-form equation fits better with data than the other two forms, and the measured dimensionless *θ* value was approximately 0.0032, which was applied to perform mathematical prediction of our experimental results. All of these fitting analyses were performed using the Nonlinearmodelfit function of the Wolfram Mathematica (version 12.4).

### **S4 Liquid cultivation of our synthetic microbial communities**

To prepare the inoculum, *P. stutzeri* strains were first grown at 30˚C RB liquid medium [21] by shaking at 220 rpm, supplemented with 50 μg/mL gentamicin. The cells were then washed by the minimum medium [22] for twice to make an inoculum. For co-culture experiments, inocula of two strains the were concentrated to an Optical density (OD, measured at 600 nm) of 5.0, and mixed at a 1:1 ratio, and then inoculated to 96-well plates that contains 120 μL fresh minimum medium (starting OD: 0.05), supplemented with 2 mM IPTG, 50 μg/mL gentamicin, as well as salicylate as the sole carbon source. During the cultivation, OD was measured to estimate the total biomass, and fluorescence intensity was measured to estimate the growth of each population. The relative fraction was calculated by a method previously described[23]. Briefly, cultures of each populations were grown to mid-log phase at 30˚C (OD: ~0.3), diluted two-fold for eleven times, and the dilutions measured for their OD and fluorescence. Correlations between OD and fluorescence were then determined using the basic method defined in the LinerModelFit function of Mathematica software. Eventually, fluorescence values were transformed to the OD-estimated biomass to assess the growth of each population, and relative fraction was then calculated. These measurements, as well as the related measurements described below, were performed using a microplate reader (Molecular Devices, Sunnyvale, America).

### **S5 Colony pattern formation assays**

#### **S5.1 Colony pattern formation assays**

Minimum medium (1.5% agarose, Takara, Dalian, China), supplemented with 2 mM IPTG, 50 μg/mL gentamicin, and supplying salicylate as sole carbon source, was used in these studies. To prepare the culture plate, five milliliters of this medium was poured in a Petri dish (60 mm in diameter) and left on the bench overnight before inoculation. Preparation of the inocula is performed by the same way as liquid cultivation section. Inocula of two strains were then concentrated to an OD of 1.0, and mixed at a 1:1 ratio. For each colony, 1 μl of the inoculum was spotted on the prepared plate, after which it was allowed to dry for 10 min. After the inoculum dried on the plate, the plates were incubated at 30 ˚C for 120 h.

#### **S5.2 Microscopy imaging**

Colony patterns were imaged under 5× objective using a Leica DM6000B fluorescence microscope (Leica Corporation, Wetzlar, Germany) equipped with a LED fluorescence illuminator (Leica Corporation). Images were recorded with a DFC360 FX camera (Leica Corporation) using a GFP filter cube for eGFP (exciter: 475/40; emitter: 525/50: beamsplitter: 495) and a TX2 filter cube for mCherry (exciter: 560/40; emitter: 645/75: beamsplitter: 595). Tile scan function of Leica LAS X acquisition software (Leica Microsystems) were applied to assemble the full view of a colony from multiple fields, and composite images were also created by this software.

#### **S5.3 Measurements of relative fraction of each population in the colonies**

Two methods were applied to measure the relative fraction of each population in the colonies. Image analyses of the colonies were applied using the related functions in *Wolfram Mathematica* (version 12.4). After microscopy imaging, the assembled eGFP and mCherry images were separately exported as grayscale tiff files. Then *ColorQuantize* function was used to give an approximation to the image by quantizing it to distinct colors. Subsequently the images were transformed into binarized data using *ImageData* function. To separately analyze the community structures in different colony region, pixels were were divided into two groups according to their distance to the colony center. Pixels located in the inoculating-region (inside a circle with a radius of approximately 1mm) were grouped as inoculating-region pixels, while the rest were grouped as the expansion-region pixels. Relative fraction of each population in the two regions, as well as in the entire colony, was measured by counting the frequency of pixels belonging to each color group. These analyses were performed in custom Wolfram Mathematica scripts. The source codes used are available on Github: <https://github.com/RoyWang1991/MDOLcode/tree/master/MDOL-spatial>.

### **S6 Supplementary Tables**

**Supplementary Table 1 Definitions of variables and dimensionless methods for the ODE model in this study**

| **Variable** | **Description** | **Units** | **ND variable** | **ND Value Range** |
| --- | --- | --- | --- | --- |
| $\text{S}_{\text{D,in}}$ | Concentration of Intracellular *substrate* in the *Detoxifier*. | M | $\text{s}_{\text{D,in}}\text{=}\text{S}_{\text{D,in}}\text{/}\text{K}_{\text{1}}$ | 0 ~ 10^4^ |
| $\text{S}_{\text{E,in}}$ | Concentration of Intracellular *substrate* in the *Embezzler*. | M | $\text{s}_{\text{E,in}}\text{=}\text{S}_{\text{E,in}}\text{/}\text{K}_{\text{1}}$ | 0 ~ 10^4^ |
| $\text{S}_{\text{out}}$ | Concentration of extracellular *substrate*. | M | $\text{s}_{\text{out}}\text{=}\text{S}_{\text{out}}\text{/}\text{K}_{\text{1}}$ | 0 ~ 10^4^ |
| $\text{I}_{\text{D,in}}$ | Concentration of Intracellular *intermediate* in the *Detoxifier*. | M | $\text{i}_{\text{D,in}}\text{=}\text{I}_{\text{D,in}}\text{/}\text{K}_{\text{1}}$ | 0 ~ 10^4^ |
| $\text{I}_{\text{E,in}}$ | Concentration of Intracellular *intermediate* in the *Embezzler*. | M | $\text{i}_{\text{E,in}}\text{=}\text{I}_{\text{E,in}}\text{/}\text{K}_{\text{1}}$ | 0 ~ 10^4^ |
| $\text{I}_{\text{out}}$ | Concentration of extracellular *intermediate*. | M | $\text{i}_{\text{out}}\text{=}\text{I}_{\text{out}}\text{/}\text{K}_{\text{1}}$ | 0 ~ 10^4^ |
| $\text{P}_{\text{D,in}}$ | Concentration of Intracellular *product* in the *Detoxifier*. | M | $\text{p}_{\text{D,in}}\text{=}\text{P}_{\text{D,in}}\text{/}\text{K}_{\text{1}}$ | 0 ~ 10^4^ |
| $\text{P}_{\text{E,in}}$ | Concentration of Intracellular *product* in the *Embezzler*. | M | $\text{p}_{\text{E,in}}\text{=}\text{P}_{\text{E,in}}\text{/}\text{K}_{\text{1}}$ | 0 ~ 10^4^ |
| $\text{P}_{\text{out}}$ | Concentration of extracellular *product*. | M | $\text{p}_{\text{out}}\text{=}\text{P}_{\text{out}}\text{/}\text{K}_{\text{1}}$ | 0 ~ 10^4^ |
| $\text{X}_{\text{D}}$ | Total volume of *Detoxifier* population. | L^-1^ | $\text{x}_{\text{D}}\text{=}\text{X}_{\text{D}}\text{/}\text{V}_{\text{E}}$ | 0 ~ $\rho$ |
| $\text{X}_{\text{E}}$ | Total volume of *Embezzler* population. | L^-1^ | $\text{x}_{\text{E}}\text{=}\text{X}_{\text{E}}\text{/}\text{V}_{\text{E}}$ | 0 ~ $\rho$ |
| $\text{t}$ | Time | s | $\text{τ=t∙}\text{k}_{\text{1}}$ | 0 ~ 10^7^ |

**Supplementary Table 2 Definitions, nondimensionalization and values of the parameters for the ODE model in this study**

| **Parameter** | **Description** | **Value** | **ND parameter** | **Default value Search range** | **Source** |
| --- | --- | --- | --- | --- | --- |
| $\text{K}_{\text{j}}$ | Michaelis-Menten constant for the *jth* reaction (*j* =1, 2). | 10^-5^ M | $\text{β}_{\text{j}}\text{=}\text{K}_{\text{j}}\text{/}\text{K}_{\text{1}}$ | $\text{β}_{\text{1}}\text{=1}$, $\text{β}_{2}\text{=1}$  10^-2^~10^2^ | [24] |
| $\text{E}_{\text{j}}$ | Concentration of the *jth* enzyme. | 10^-3^ M | $\text{e}_{\text{j}}\text{=}\text{E}_{\text{j}}\text{/}\text{K}_{\text{1}}$  $\text{α}_{\text{j}}\text{=}\text{k}_{\text{j}}\text{/}\text{k}_{\text{1}}$, $\text{a}_{\text{1}}\text{=}\text{e}_{\text{1}}$, $\text{a}_{\text{2}}\text{=}\text{α}_{\text{2}}\text{e}_{\text{2}}$ | $\text{a}_{\text{1}}\text{=}$10000, $\text{a}_{\text{2}}\text{=}$1000  1~10^4^ | [24-26] |
| $\text{k}_{\text{j}}$ | Specific rate of the *jth* reaction (*j* =1, 2). | 10 s^-1^ |  |  |  |
| $\text{r}_{\text{s}}$, $\text{r}_{\text{i}}$, $\text{r}_{\text{p}}$ | Transport rate of S, I, and P. | 10 s^-1^ | $\text{γ}_{\text{s}}\text{=}\text{r}_{\text{s}}\text{/}\text{k}_{\text{1}}$, $\text{γ}_{\text{i}}\text{=}\text{r}_{\text{i}}\text{/}\text{k}_{\text{1}}$, $\text{γ}_{\text{p}}\text{=}\text{r}_{\text{p}}\text{/}\text{k}_{\text{1}}$ | 1  10^-2^~10^2^ | [27] |
| $\text{Kg}_{\text{1}}$, $\text{Kg}_{\text{2}}$ | Half-saturation constant of Monod growth. | 10^-5^ M | $\text{bg}\text{=}\text{K}_{\text{g}}\text{/}\text{K}_{\text{1}}$ | 1  10^-2^~10^2^ | [28] |
| $\text{kg}$ | Maximum consumption rate of *product*. | 10^-4^ M⸱s^-1^ | $\text{Cp=}\text{kg/}\text{(}\text{K}_{\text{1}}\text{k}_{\text{1}}\text{)}$ | 10  10^-2^~10^2^ | [28] |
| $\text{g}_{\text{D}}$, $\text{g}_{\text{E}}$ | Specific growth rate of the two populations. | s^-1^ | $\text{μ}_{\text{1}}\text{=}\frac{\text{Cp}}{\text{bg}\text{+}\text{p}_{\text{1,in}}}\text{p}_{\text{1,in}}\text{y}$*,* $\text{μ}_{\text{2}}\text{=}\frac{\text{Cp}}{\text{β}_{\text{g}}\text{+}\text{p}_{\text{2,in}}}\text{p}_{\text{2,in}}\text{y}$ |  |  |
| $\text{Y}$ | Yield coefficient for biomass production | 10^2^ M^-1^ | $\text{y=Y∙}\text{K}_{\text{1}}$ | 1  10^-5^~10^-3^ | [7] |
| $\text{N}_{\text{m}}$ | Carrying capacity of the whole community | 10^-6^ L^-1^ | $\text{ρ=}\text{N}_{\text{m}}\text{/}\text{V}_{\text{E}}$ | 1  10^-3^ ~10^-1^ | [28] |
| $\text{V}_{\text{E}}$ | The volume of extracellular space. | 10^-4^ L |  |  |  |
| $\text{S}_{\text{0}}$ | Initial concentration of *substrate* | 10^-2^ M | $\text{s}_{\text{0}}\text{=}\text{S}_{\text{0}}\text{/}\text{K}_{\text{1}}$ | 0~10^4^ |  |

**Supplementary Table 3 Definitions, dimensionless methods and values/ranges of different toxic coefficients**

| **Variables**  **(Description)** | **Formula*** | **ND variables** | **ND formula** | **Value Range (**$\boldsymbol{\theta}$**)** |
| --- | --- | --- | --- | --- |
| $\text{ToS}_{\text{1}}$, $\text{ToS}_{\text{2}}$  (Toxic coefficient of *substrate* in the two populations.) | $\frac{\text{1}}{\text{1}\text{+}\frac{\text{S}_{\text{1,in}}}{\text{K}_{\text{s,tox}}}}$, $\frac{\text{1}}{\text{1}\text{+}\frac{\text{S}_{\text{1,in}}}{\text{K}_{\text{s,tox}}}}$ | $\text{θ=}\frac{\text{K}_{\text{1}}}{\text{K}_{\text{s,tox}}}$ | $\text{to}_{\text{1}}\text{=}\frac{\text{1}}{\text{1}\text{+θ}\text{s}_{\text{1,in}}}$*,* $\text{to}_{\text{2}}\text{=}\frac{\text{1}}{\text{1}\text{+θ}\text{s}_{\text{2,in}}}$ | 0 ~ 1 |
|  | $\text{1}\text{-}\frac{\text{S}_{\text{1,in}}}{\text{K}_{\text{s,tox}}}$, $\text{1}\text{-}\frac{\text{S}_{\text{2,in}}}{\text{K}_{\text{s,tox}}}$ |  | $\text{to}_{\text{1}}\text{=}\text{(1}\text{-θ}\text{s}_{\text{1,in}}\text{)}$*,* $\text{to}_{\text{2}}\text{=}\text{(1}\text{-θ}\text{s}_{\text{2,in}}\text{)}$ | 0 ~ 1 |
|  | $\text{e}^{\text{-}\frac{\text{S}_{\text{1,in}}}{\text{K}_{\text{s,tox}}}}$, $\text{e}^{\text{-}\frac{\text{S}_{\text{2,in}}}{\text{K}_{\text{s,tox}}}}$ |  | $\text{to}_{\text{1}}\text{=}\text{e}^{\text{-θ}\text{s}_{\text{1,in}}}$*,* $\text{to}_{\text{2}}\text{=}\text{e}^{\text{-θ}\text{s}_{\text{2,in}}}$ | 0 ~ 1 |

*Note: The three expressions in the second and fourth column show the reciprocal, linear, and logarithmic forms of toxic coefficients, respectively

**Supplementary Table 4** Multiple linear regression analyses of the first-round simulation results of our ODE model

| Parameter | Without Substrate toxicity | | | With Substrate toxicity | | | | | | | | |
| --- | --- | --- | --- | --- | --- | --- | --- | --- | --- | --- | --- | --- |
|  | $\boldsymbol{to=0}$ | | | $\boldsymbol{to}$ follows reciprocal form | | | $\boldsymbol{to}$ follows exponential form | | | $\boldsymbol{to}$ follows linear form | | |
|  | n= 177,147 | | | n= 708, 588 | | | n= 708, 588 | | | n= 708, 588 | | |
|  | Coffs* | P-value | R^2**^ | Coffs* | P-value | R^2**^ | Coffs* | P-value | R^2**^ | Coffs* | P-value | R^2**^ |
| $\text{y}$ | -0.0083 | <0.0001 | 0.844 | -0.0052 | <0.0001 | 0.845 | -0.0050 | <0.0001 | 0.770 | 0.0016 | <0.0001 | 0.833 |
| $\text{Cp}$ | -0.3043 | <0.0001 |  | -0.2262 | <0.0001 |  | -0.2311 | <0.0001 |  | -0.2615 | <0.0001 |  |
| $\text{bg}$ | 0.0040 | <0.0001 |  | 0.0044 | <0.0001 |  | 0.0041 | <0.0001 |  | 0.0003 | <0.0001 |  |
| $\text{a}_{\text{1}}$ | 0.0305 | <0.0001 |  | 0.0733 | <0.0001 |  | 0.0509 | <0.0001 |  | 0.0814 | <0.0001 |  |
| $\text{a}_{\text{2}}$ | 0.0085 | <0.0001 |  | 0.0021 | <0.0001 |  | 0.0028 | <0.0001 |  | 0.0005 | <0.0001 |  |
| $\text{β}_{\text{2}}$ | -0.0012 | 0.260 |  | -0.0012 | 0.0848 |  | -0.0015 | 0.0415 |  | 0.0000 | 0.0118 |  |
| $\text{γ}_{\text{s}}$ | 0.1175 | <0.0001 |  | -0.0993 | <0.0001 |  | -0.0919 | <0.0001 |  | -0.1256 | <0.0001 |  |
| $\text{γ}_{\text{i}}$ | 0.0457 | <0.0001 |  | 0.0521 | <0.0001 |  | 0.0570 | <0.0001 |  | 0.0573 | <0.0001 |  |
| $\text{γ}_{\text{p}}$ | 0.1077 | <0.0001 |  | 0.0978 | <0.0001 |  | 0.0875 | <0.0001 |  | 0.0913 | <0.0001 |  |
| $\text{ρ}$ | -0.0016 | 0.149 |  | -0.0011 | <0.0001 |  | -0.0030 | <0.0001 |  | -0.0001 | <0.0001 |  |
| $\text{s}_{\text{0}}$ | 0.2319 | <0.0001 |  | 0.2612 | <0.0001 |  | 0.2504 | <0.0001 |  | 0.2779 | <0.0001 |  |
| $\text{θ}$ | N.A. | |  | 0.2464 | <0.0001 |  | 0.2623 | <0.0001 |  | 0.2976 | <0.0001 |  |

* The coefficient of each parameter in the result of multiple linear regression analyses.

** Shown is the value of the adjusted R-square.

**Supplementary Table 5** Multiple linear regression analyses of the second-round simulation results indicate the relative effects of the seven key parameters involved in the ODE model on the final fraction of Detoxifier, as well as the four fitting parameters in Eqn[1] or Eqn [2] (i.e., $\text{ks}$, $\text{Fd}_{\text{max}}$, $\text{kt}$, $\text{Ts}_{\text{max}}$).

| Parameters | Without Substrate toxicity | | | With Substrate toxicity | | | | | | | | |
| --- | --- | --- | --- | --- | --- | --- | --- | --- | --- | --- | --- | --- |
|  | $\text{to}\text{ }\text{=}\text{ }\text{0}$ | | | $\text{to}$ follows reciprocal form | | | $\text{to}$ follows exponential form | | | $\text{to}$ follows linear form | | |
|  | n= 163, 296 | | | n= 2, 776, 032 | | | n= 2, 776, 032 | | | n= 2, 776, 032 | | |
|  | Coffs* | P-value | R^2**^ | Coffs* | P-value | R^2**^ | Coffs* | P-value | R^2**^ | Coffs* | P-value | R^2**^ |
| $\text{DF}$ | | | | | | | | | | | | |
| $\text{Cp}$ | -0.5088 | <0.0001 | 0.894 | -0.4117 | <0.0001 | 0.909 | 0.1963 | <0.0001 | 0.810 | -0.3731 | <0.0001 | 0.735 |
| $\text{a}_{\text{1}}$ | 0.0655 | <0.0001 |  | 0.1309 | <0.0001 |  | -0.3848 | <0.0001 |  | 0.1407 | <0.0001 |  |
| $\text{γ}_{\text{s}}$ | 0.0704 | <0.0001 |  | -0.1702 | <0.0001 |  | 0.1383 | <0.0001 |  | -0.1113 | <0.0001 |  |
| $\text{γ}_{\text{i}}$ | 0.1194 | <0.0001 |  | 0.1219 | <0.0001 |  | -0.1750 | <0.0001 |  | 0.1059 | <0.0001 |  |
| $\text{γ}_{\text{p}}$ | 0.1696 | <0.0001 |  | 0.1232 | <0.0001 |  | 0.1266 | <0.0001 |  | 0.1279 | <0.0001 |  |
| $\text{s}_{\text{0}}$ | 0.3257 | <0.0001 |  | 0.3226 | <0.0001 |  | 0.1220 | <0.0001 |  | 0.2894 | <0.0001 |  |
| $\theta$ | N.A. | |  | 0.4350 | <0.0001 |  | 0.3411 | <0.0001 |  | 0.3674 | <0.0001 |  |
| $kd$ | | | | | | | | | | | | |
| $\text{Cp}$ | 0.0923 | <0.0001 | 0.899 | 0.0924 | <0.0001 | 0.747 | 0.1309 | <0.0001 | 0.766 | 0.1289 | <0.0001 | 0.751 |
| $\text{a}_{\text{1}}$ | -0.0055 | 0.0108 |  | -0.0024 | 0.2568 |  | -0.0076 | 0.0042 |  | -0.0035 | 0.0141 |  |
| $\text{γ}_{\text{s}}$ | 0.0558 | <0.0001 |  | 0.0304 | <0.0001 |  | 0.0102 | 0.0001 |  | 0.0209 | 0.0209 |  |
| $\text{γ}_{\text{i}}$ | -0.0431 | <0.0001 |  | -0.0392 | <0.0001 |  | -0.0557 | <0.0001 |  | -0.0536 | <0.0001 |  |
| $\text{γ}_{\text{p}}$ | -0.0197 | 0.0064 |  | -0.0250 | <0.0001 |  | -0.0248 | <0.0001 |  | -0.0153 | <0.0001 |  |
| $\text{Fd}_{\text{max}}$ | | | | | | | | | | | | |
| $\text{Cp}$ | -0.7384 | <0.0001 | 0.870 | -0.7163 | <0.0001 | 0.812 | -0.6952 | <0.0001 | 0.897 | -0.7279 | <0.0001 | 0.772 |
| $\text{a}_{\text{1}}$ | 0.2385 | <0.0001 |  | 0.2388 | <0.0001 |  | 0.2621 | <0.0001 |  | 0.2730 | <0.0001 |  |
| $\text{γ}_{\text{s}}$ | -0.0152 | 0.0400 |  | 0.0505 | 0.0296 |  | 0.0732 | <0.0001 |  | 0.0920 | <0.0001 |  |
| $\text{γ}_{\text{i}}$ | 0.1917 | <0.0001 |  | 0.0948 | <0.0001 |  | 0.1106 | <0.0001 |  | 0.0292 | <0.0001 |  |
| $\text{γ}_{\text{p}}$ | 0.2182 | <0.0001 |  | 0.2386 | <0.0001 |  | 0.2385 | <0.0001 |  | 0.2101 | <0.0001 |  |
| $\text{kt}$ | | | | | | | | | | | | |
| $\text{Cp}$ | N.A. | | | 0.1425 | <0.0001 | 0.798 | 0.1632 | <0.0001 | 0.840 | 0.1762 | <0.0001 | 0.768 |
| $\text{a}_{\text{1}}$ |  |  |  | 0.1003 | <0.0001 |  | 0.1052 | <0.0001 |  | 0.1038 | <0.0001 |  |
| $\text{γ}_{\text{s}}$ |  |  |  | -0.0627 | <0.0001 |  | -0.0597 | <0.0001 |  | -0.0206 | <0.0001 |  |
| $\text{γ}_{\text{i}}$ |  |  |  | -0.0321 | <0.0001 |  | -0.0127 | <0.0001 |  | -0.0357 | <0.0001 |  |
| $\text{γ}_{\text{p}}$ |  |  |  | -0.0273 | <0.0001 |  | -0.0110 | <0.0001 |  | -0.0646 | <0.0001 |  |
| $\text{Ts}_{\text{max}}$ | | | | | | | | | | | | |
| $\text{Cp}$ | N.A. | | | 0.1439 | <0.0001 | 0.774 | 0.1417 | <0.0001 | 0.810 | 0.1277 | <0.0001 | 0.803 |
| $\text{a}_{\text{1}}$ |  |  |  | 0.2500 | <0.0001 |  | 0.2781 | <0.0001 |  | 0.2521 | <0.0001 |  |
| $\text{γ}_{\text{s}}$ |  |  |  | -0.2432 | <0.0001 |  | -0.2623 | <0.0001 |  | -0.2592 | <0.0001 |  |
| $\text{γ}_{\text{i}}$ |  |  |  | -0.0054 | 0.4390 |  | -0.0047 | 0.1254 |  | -0.0594 | <0.0001 |  |
| $\text{γ}_{\text{p}}$ |  |  |  | -0.0152 | <0.0001 |  | -0.0110 | <0.0001 |  | -0.0530 | <0.0001 |  |

* The coefficient of each parameter in the result of multiple linear regression analyses.

** Shown is the value of the adjusted R-square.

**Supplementary Table 6** Values of the parameters used in the model for predicting the experimental results

| **Parameter** | **Input Value** | **Source** | **ND parameter** | **ND Value** |
| --- | --- | --- | --- | --- |
| $\text{K}_{\text{1}}$ | 9.6×10^-6^ M | [29] | $\text{β}_{\text{1}}\text{=}\text{K}_{\text{1}}\text{/}\text{K}_{\text{1}}$ | 1 |
| $\text{k}_{\text{1}}$ | 4.6 s^-1^ | [29] | $\text{e}_{\text{1}}\text{=}\text{E}_{\text{1}}\text{/}\text{K}_{\text{1}}$  $\text{α}_{\text{1}}\text{=}\text{k}_{\text{1}}\text{/}\text{k}_{\text{1}}$*,* $\text{a}_{\text{1}}\text{=}\text{e}_{\text{1}}$ | 2080 |
| $\text{E}_{\text{1}}$ | 2×10^-2^ M | [26] |  |  |
| $\text{K}_{\text{2}}$ | 1.5×10^-6^ M | [30] | $\text{β}_{\text{2}}\text{=}\text{K}_{\text{2}}\text{/}\text{K}_{\text{1}}$ | 0.156 |
| $\text{k}_{\text{2}}$ | 200 s^-1^ | [29] | $\text{e}_{\text{2}}\text{=}\text{E}_{\text{2}}\text{/}\text{K}_{\text{1}}$  $\text{α}_{\text{2}}\text{=}\text{k}_{\text{2}}\text{/}\text{k}_{\text{1}}$*,* $\text{a}_{\text{2}}\text{=}\text{α}_{\text{2}}\text{e}_{\text{1}}$ | 90600 |
| $\text{E}_{\text{2}}$ | 2×10^-2^ M | [26] |  |  |
| $\text{r}_{\text{s}}$ | 26.3 s^-1^ | [27] | $\text{γ}_{\text{s}}\text{=}\text{r}_{\text{s}}\text{/}\text{k}_{\text{1}}$ | 5.71 |
| $\text{r}_{\text{i}}$ | 4.27 s^-1^ | [27] | $\text{γ}_{\text{i}}\text{=}\text{r}_{\text{i}}\text{/}\text{k}_{\text{1}}$ | 0.928 |
| $\text{r}_{\text{p}}$ | 25.1 s^-1^ | [27] | $\text{γ}_{\text{p}}\text{=}\text{r}_{\text{p}}\text{/}\text{k}_{\text{1}}$ | 5.47 |
| $\text{V}_{\text{E}}$ | 1.5×10^-4^ L | Experimental settings |  |  |
| $\text{K}_{\text{g}}$ | 2×10^-5^ M | Experimental measure | $\text{bg}\text{=}\text{K}_{\text{g}}\text{/}\text{K}_{\text{1}}$ | 2.08 |
| $\text{kg}$ | 6.37×10^-4^ M⸱s^-1^ | Experimental measure | $\text{Ig=}\text{kg/}\text{(}\text{K}_{\text{1}}\text{k}_{\text{1}}\text{)}$ | 14.4 |
| $\text{Y}$ | 22.92 M^-1^ | Experimental measure | $y=Y\cdot K_{1}$ | 2.2×10^-4^ |
| $\text{N}_{\text{m}}$ | 1.6 ×10^-5^ L^-1^ | Experimental measure | $\rho=N_{m}/V_{E}$ | 0.11 |
| $\text{S}_{\text{0}}$ | 10^-3^~10^-1^ C-M | Experimental settings | $\text{s}_{\text{0}}\text{=}\text{S}_{\text{0}}\text{/}\text{K}_{\text{1}}$ | 10 ~ 1000 |
| $\text{K}_{\text{s,tox}}$ | 3.00×10^-3^ M | Experimental measure | $\text{θ}_{\text{s}}\text{=}\text{K}_{\text{1}}\text{/}\text{K}_{\text{s,tox}}$ | 0.0032 |

**Supplementary Table 7** Variables and parameters used in our individual-based model

| **Variables/parameters** | **Description** | **Value and Units** | **Source** |
| --- | --- | --- | --- |
| **Variables** | | | |
| $\text{S}_{\text{D,in,i}}$ | Concentration of Intracellular substrate in the *ith* cell of Detoxifier population. | C-mmol/L |  |
| $\text{S}_{\text{E,in,i}}$ | Concentration of Intracellular substrate in the *ith* cell of Embezzler population. | C-mmol/L |  |
| $\text{S}_{\text{out,i}}$ | Concentration of extracellular substrate around the *ith* cell. | C-mmol/L |  |
| $\text{I}_{\text{D,in,i}}$ | Concentration of Intracellular intermediate in the *ith* cell of Detoxifier population. | C-mmol/L |  |
| $\text{I}_{\text{E,in,i}}$ | Concentration of Intracellular intermediate in the *ith* cell of Embezzler population. | C-mmol/L |  |
| $\text{I}_{\text{out,i}}$ | Concentration of extracellular intermediate around the *ith* cell. | C-mmol/L |  |
| $\text{P}_{\text{D,in,i}}$ | Concentration of Intracellular final product in the *ith* cell of Detoxifier population. | C-mmol/L |  |
| $\text{P}_{\text{E,in,i}}$ | Concentration of Intracellular final product in the *ith* cell of Embezzler population. | C-mmol/L |  |
| $\text{P}_{\text{out,i}}$ | Concentration of extracellular final product around the *ith* cell. | C-mmol/L |  |
| $\text{V}_{\text{D,i}}$ | Volume of the *ith* Detoxifier cells | fL |  |
| $\text{V}_{\text{E,i}}$ | Volume of the *ith* Embezzler cells | fL |  |
| $\text{L}_{\text{D,i}}$ | Length of the *ith* Detoxifier cells | μm |  |
| $\text{L}_{\text{E,i}}$ | Length of the *ith* Embezzler cells | μm |  |
| $\text{g}_{\text{D,i}}$ | Growth rate of the *ith* Detoxifier cells | s^-1^ |  |
| $\text{g}_{\text{E,i}}$ | Growth rate of the *ith* Embezzler cells | s^-1^ |  |
| **Parameters** | | | |
| $\text{K}_{\text{1}}$ | Michaelis-Menten constant for the first reaction | 9.6×10^-6^ M | [29] |
| $\text{k}_{\text{1}}$ | Specific rate of the first reaction. | 4.6 s^-1^ | [29] |
| $\text{E}_{\text{1}}$ | Concentration of the first enzyme. | 2×10^-2^ M | [26] |
| $\text{K}_{\text{2}}$ | Michaelis-Menten constant for the second reaction | 1.5×10^-6^ M | [30] |
| $\text{k}_{\text{2}}$ | Specific rate of the first reaction. | 200 s^-1^ | [29] |
| $\text{E}_{\text{2}}$ | Concentration of the first enzyme. | 2×10^-2^ M | [26] |
| $\text{r}_{\text{s}}$ | Transport rate of S | 26.3 s^-1^ | [27] |
| $\text{r}_{\text{i}}$ | Transport rate of I | 4.27 s^-1^ | [27] |
| $\text{r}_{\text{p}}$ | Transport rate of P | 25.1 s^-1^ | [27] |
| $\text{K}_{\text{g}}$ | Half-saturation constant of Monod growth. | 2×10^-5^ M | Experimental measure |
| $\text{kg}$ | Maximum consumption rate of *product*. | 6.37×10^-4^ M⸱s^-1^ | Experimental measure |
| $\text{Y}$ | Yield coefficient for biomass production. | 22.92 M^-1^ | Experimental measure |
| $\text{S}_{\text{0}}$ | Initial concentration of *substrate* | 0 ~ 20 C-M* | Experimental settings |
| $\text{K}_{\text{s,tox}}$ | toxic coefficients | 0 ~ 3×10^-2^ M* | Experimental settings |
| $\text{D}_{\text{s}}$ | Diffusivity of the S | 0.886 μm^2^/s  0.5-10.0 μm^2^/s | [27, 31, 32] |
| $\text{D}_{\text{i}}$ | Diffusivity of the I | 0.993 μm^2^/s  0.5-10.0 μm^2^/s | [27, 31, 32] |
| $\text{D}_{\text{p}}$ | Diffusivity of the P | 1.12 μm^2^/s  0.5-10.0 μm^2^/s | [27, 31, 32] |
| $\text{L}_{\text{0}}$ | Mean initial cell length | 2.0 μm | [33] |
| $\text{L}_{\text{d}}$ | Mean division cell length | 3.75 μm | [33] |
| $\text{ε}_{\text{1}}$ | Variance of initial cell length | 0.15 μm | [33] |
| $\text{ε}_{2}$ | Variance of division cell length | 0.15 μm | [33] |
| $\text{d}_{\text{c}\text{ell}}$ | Diameter of the cells | 1.0 μm | [33] |

* Dimensionless values were used to fit with simulation data and generate predicting functions, methods according to Supplementary Table 2.

### **S7 Supplementary Figures and Videos**

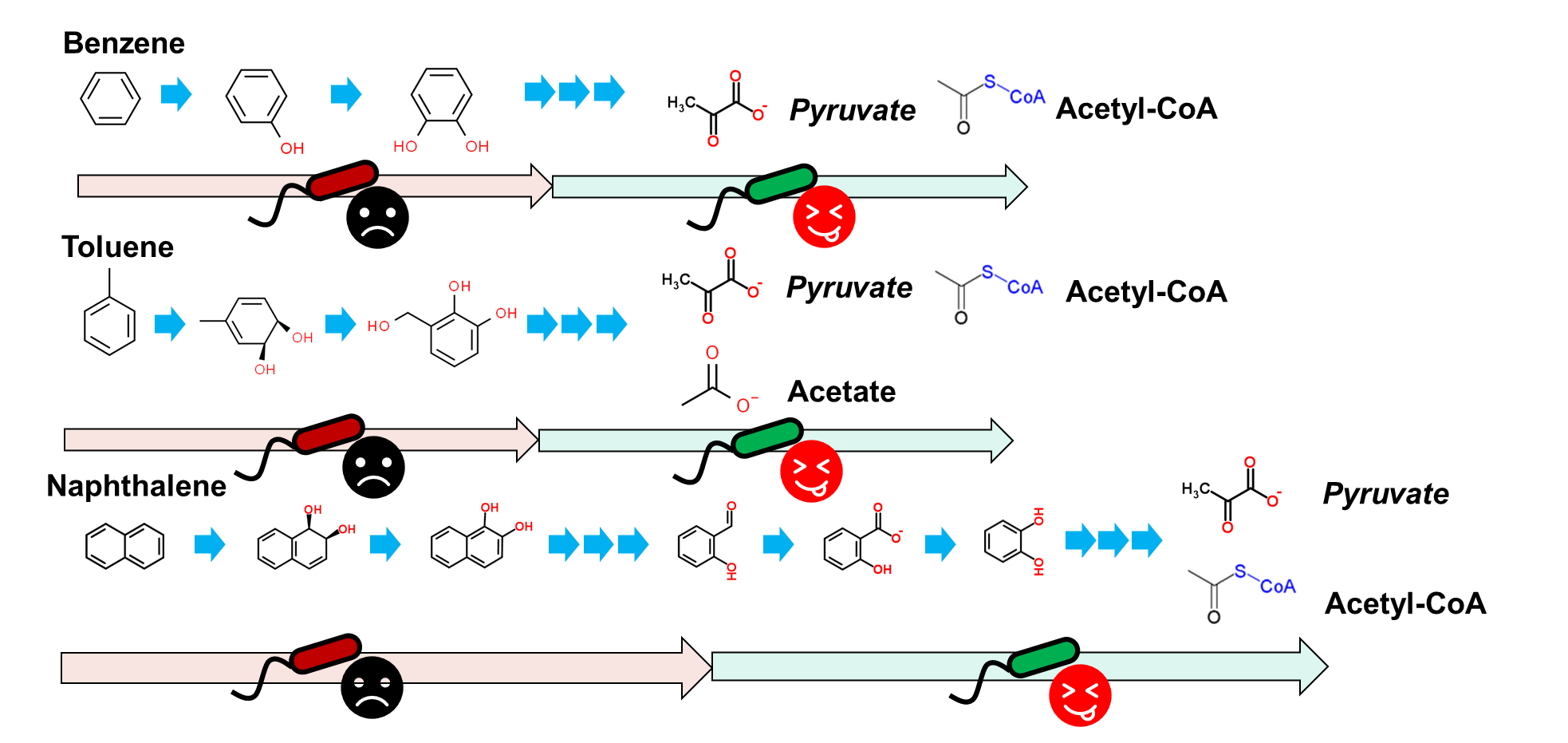

**Supplementary Figure 1** Three typical cases of organic compound degradation suggest that carbon source allocation is asymmetric between different populations when implementing these pathways by MDOL. Since direct carbon sources, such as small organic acids or coenzyme A, are normally present as the final product of a pathway, the population performing the last steps can get more benefit.

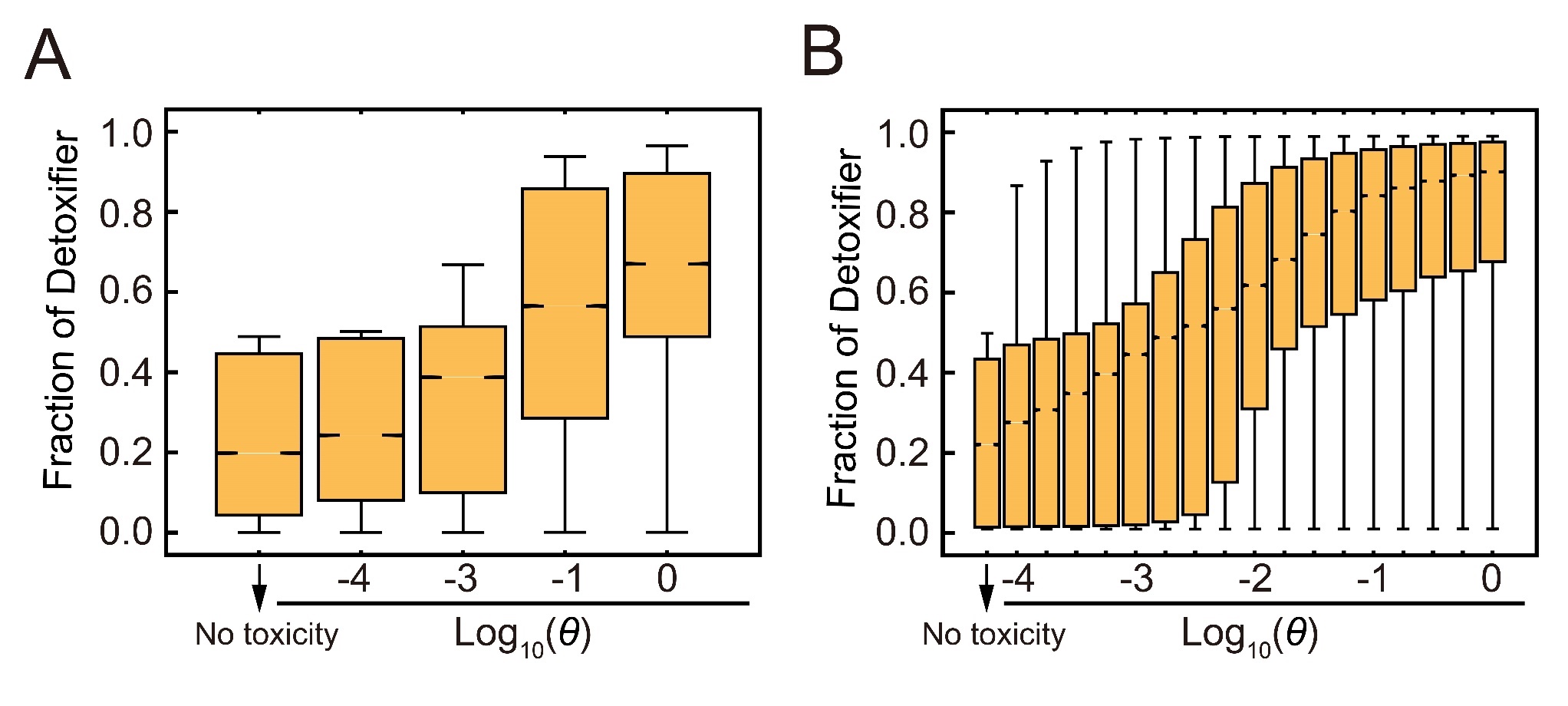

**Supplementary Figure 2** Distributions of the steady-state fraction of Detoxifier population in the first round (A) and second round (B) of simulations using our ordinary differential equation model. The simulation results that did not include substrate toxicity is denoted as ‘No toxicity’, while the simulation results initialized with different toxic strength of substrate were separately shown.

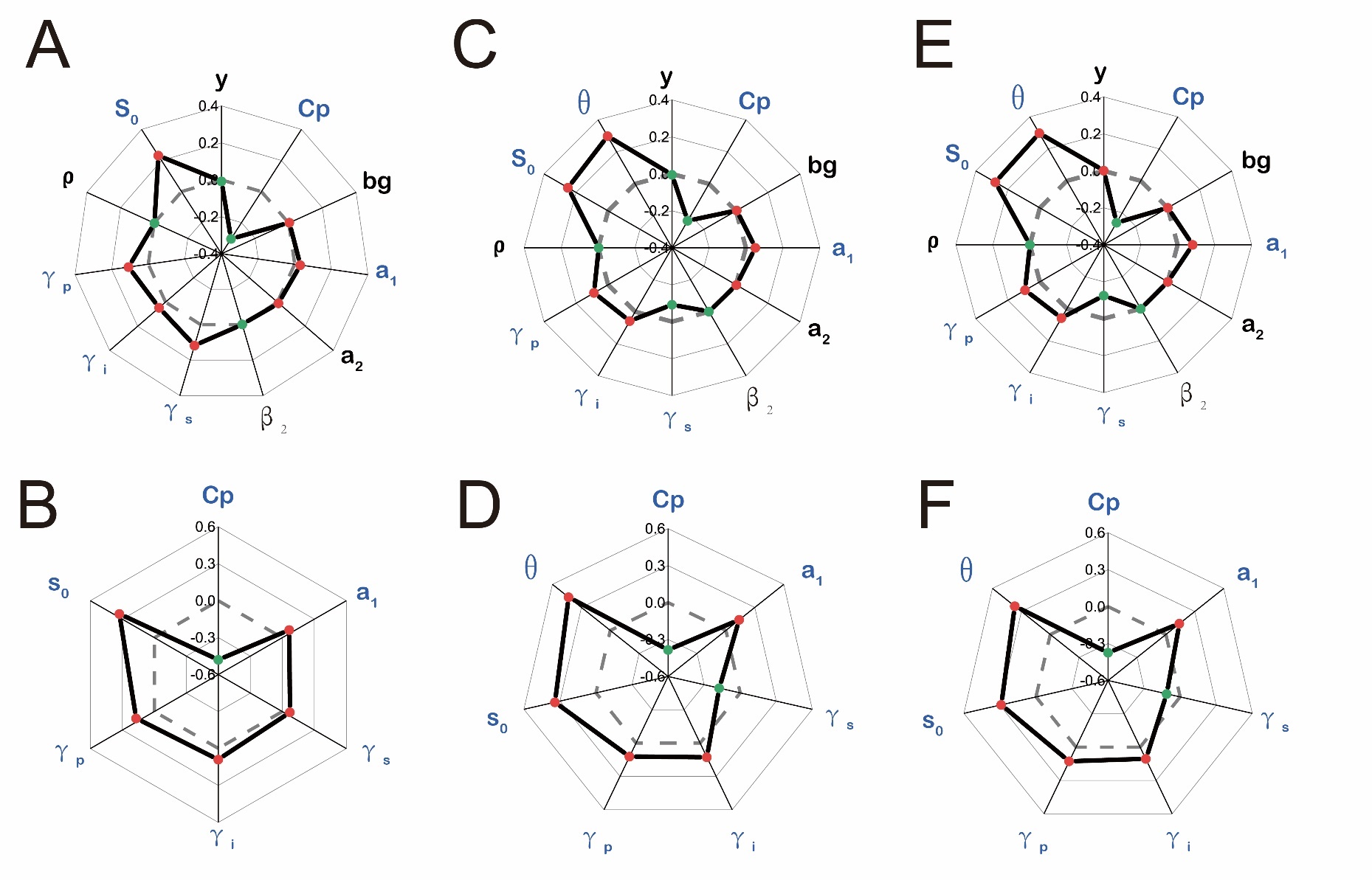

**Supplementary Figure 3** Multiple linear regression analyses of the simulation results of our ODE model. (A-B) Results from the simulations that did not include substrate toxicity. (C-D) Results from the simulations in which toxic effects of substrate was assumed to follow exponential form. (C-D) Results from the simulations in which toxic effects of substrate was assumed to follow linear form. Results from the first-round simulations are shown by (A), (C) and (E) that considered all the eleven or twelve parameters involved in our ODE model. Blue font denotes the identified key parameters. Results from the second-round simulations are shown by (B), (D) and (F), which only considered the six or seven key parameters. The axis of the radar plot denotes the values of fitting coefficients of the parameters from multiple linear regression analyses. Red dots denote the steady-state fraction of Detoxifier is positively correlated with corresponding parameter, while the green dots show the negative correlation. The origin axis (0) is highlighted by dash line to emphasize that, the value closer to zero means the corresponding parameter have smaller effect on community assembly. The data used here are listed in Supplementary Table 4 and Supplementary Table 5.

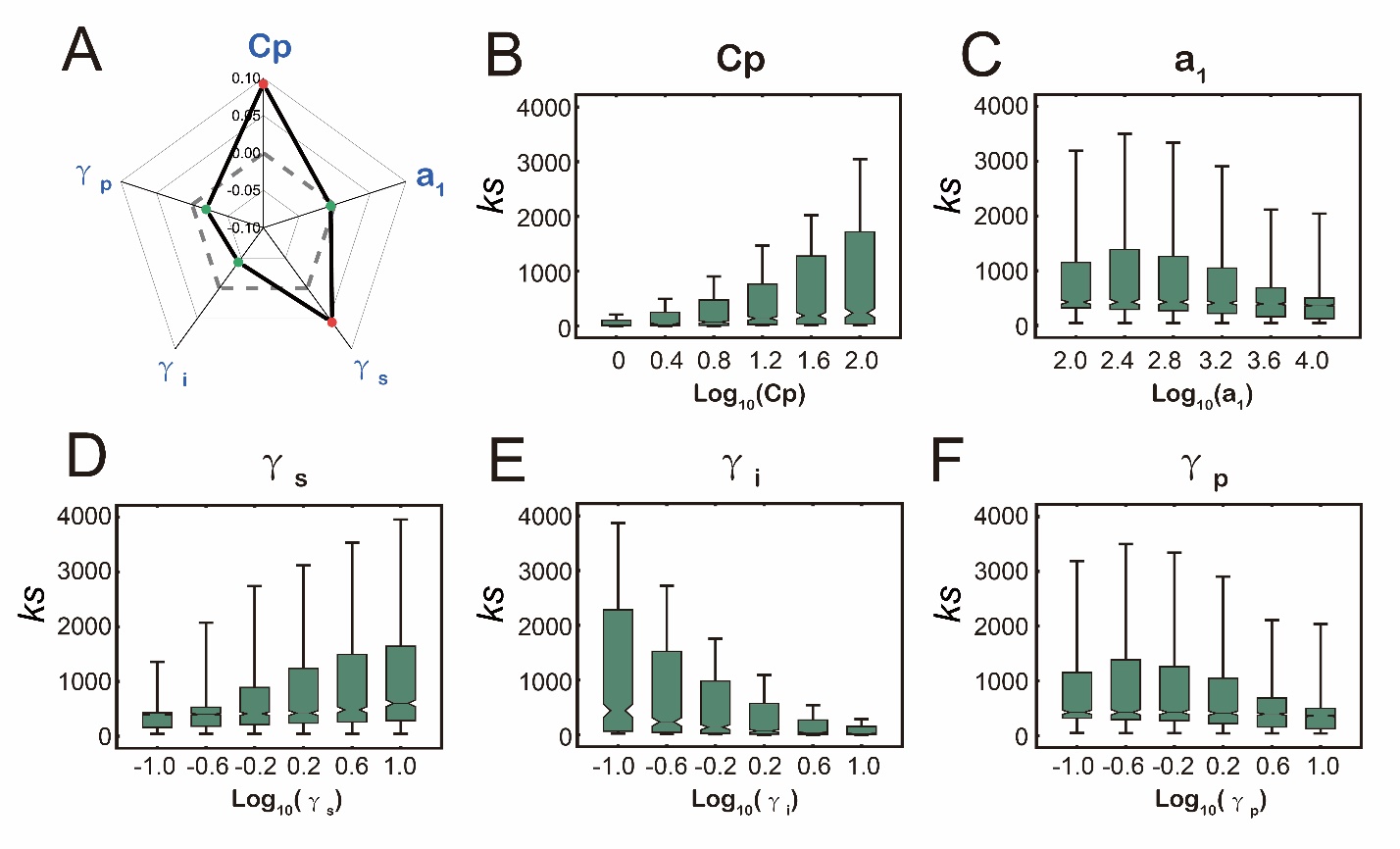

**Supplementary Figure 4** The effects of key parameters on the best fitting value of *ks* in absence of substrate toxicity. The data used here are listed in Supplementary Table 5. (A) Radar plot shows the values of fitting coefficients of the parameters from multiple linear regression analyses, indicating the degree of how the five key parameters affect the best fitting value of *ks*. (B-F) The distributions of best fitting value of *ks* along the different values of five key parameters. Data are from the second round of simulations that does not include substrate toxicity.

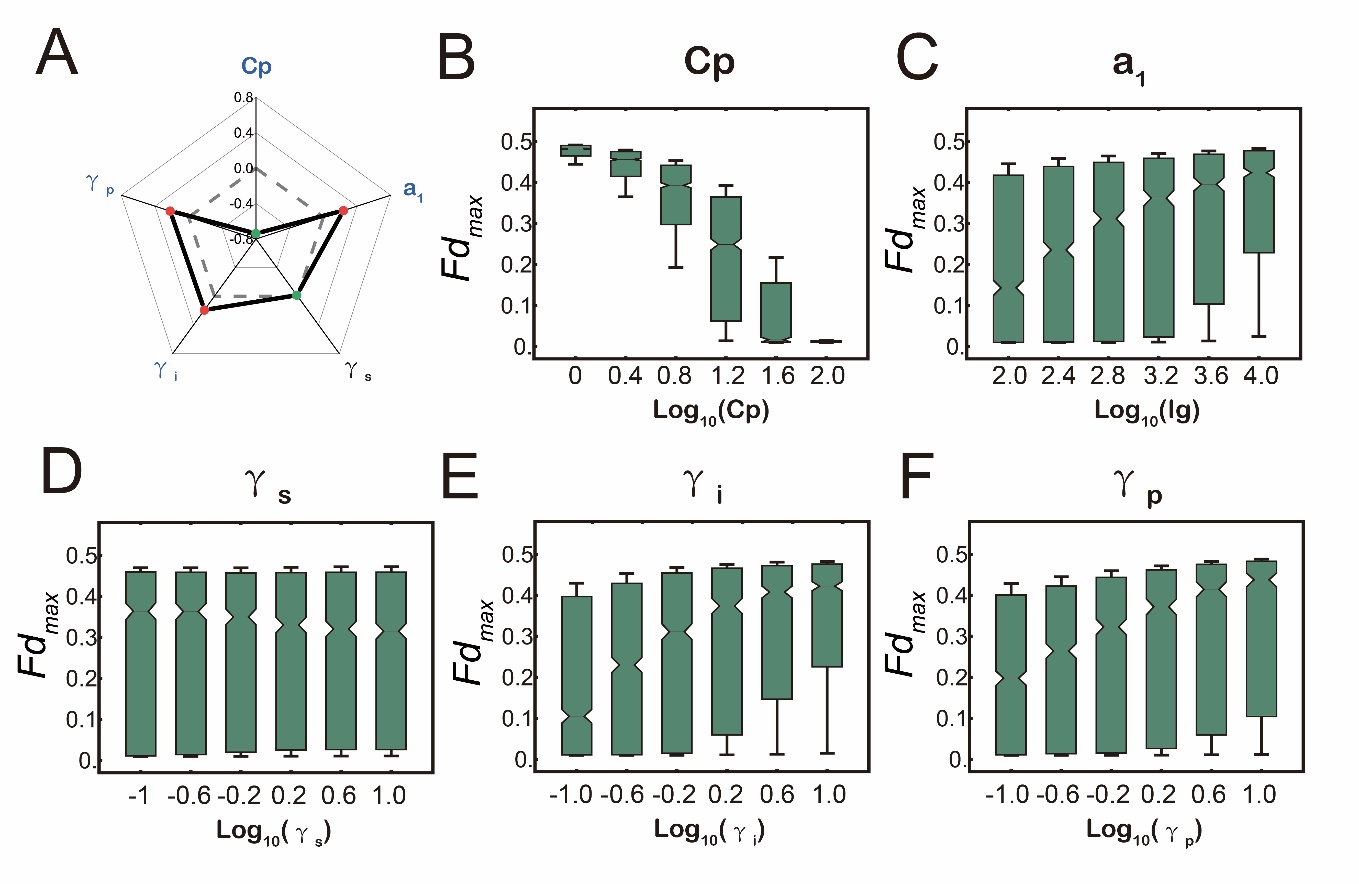

**Supplementary Figure 5** The effects of key parameters on the best fitting value of *Fd_max_* in absence of substrate toxicity. (A) Radar plot shows the values of fitting coefficients of the parameters from multiple linear regression analyses, indicating the degree of how the five key parameters affect the best fitting value of *Fd_max_*. The data used here are listed in Supplementary Table 5. (B-F) The distributions of best fitting value of *Fd_max_* along the different values of five key parameters. Data are from the second round of simulations that does not include substrate toxicity.

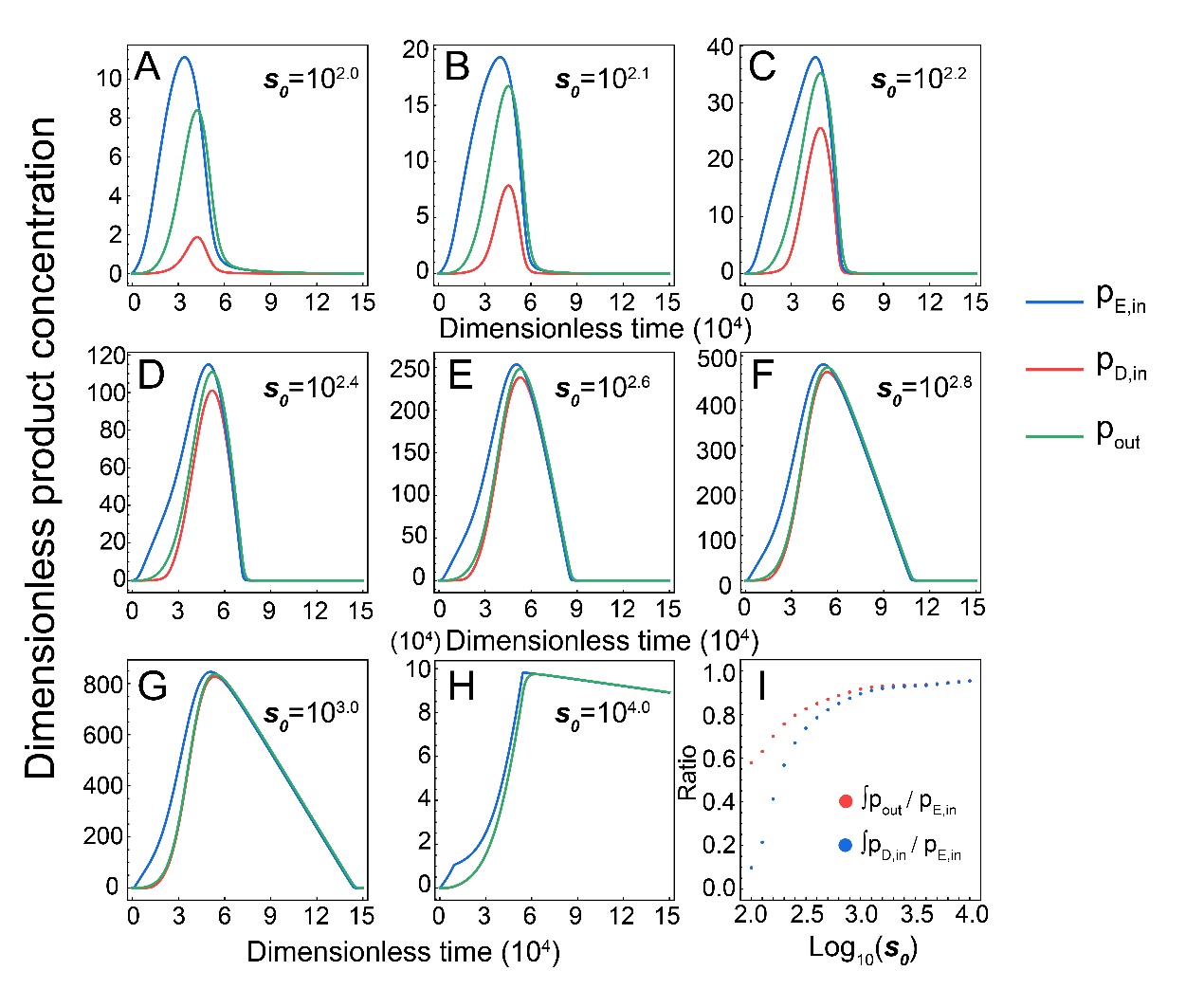

**Supplementary Figure 6** Dynamics of extra- and intracellular final product concentration of both populations in absence of substrate toxicity. (A-H) Shifts of $\text{p}_{\text{D,in}}$, $\text{p}_{\text{E,in}}$, and $\text{p}_{\text{out}}$ in the simulations with different substrate concentration. (I) Dynamics of integrated ratio of the extracellular to intracellular product concentration ($\int\text{p}_{\text{out}}/\text{p}_{\text{E,in}}$), as well as the integrated ratio of the intracellular product concentration of Detoxifier population to that of the Embezzler population ($\int\text{p}_{\text{D,in}}/\text{p}_{\text{E,in}}$), across different initial substrate concentration. Ratio $\int\text{p}_{\text{out}}/\text{p}_{\text{E,in}}$lower than 1 means majority of final product was privatized by the Embezzler cells, while higher ratio indicates more final product was released outside of cells. The plot indicates that higher substrate concentration leads to higher value of $\int\text{p}_{\text{out}}/\text{p}_{\text{E,in}}$ ratio, i.e., more release of final product. Ratio$\int\text{p}_{\text{D,in}}/\text{p}_{\text{E,in}}$ lower than 1 means Embezzler accumulate more product inside the cell than Detoxifier. The increase of $\int\text{p}_{\text{D,in}}/\text{p}_{\text{E,in}}$ indicates this benefit was weakened. The plot indicates that substrate concentration leads to higher value of $\int\text{p}_{\text{D,in}}/\text{p}_{\text{E,in}}$, suggesting benefit from product privatization decreased. Parameter values used in this simulation: *y* =10^-4^, *Cp* = 10, *bg* = 1, *a_1_* = 10000, *a_2_* = 1000, *β_2_* = 1, $\text{γ}_{\text{s}}$ = 1, $\text{γ}_{\text{i}}$ =1, $\text{γ}_{\text{p}}$ = 1, *ρ =*1.

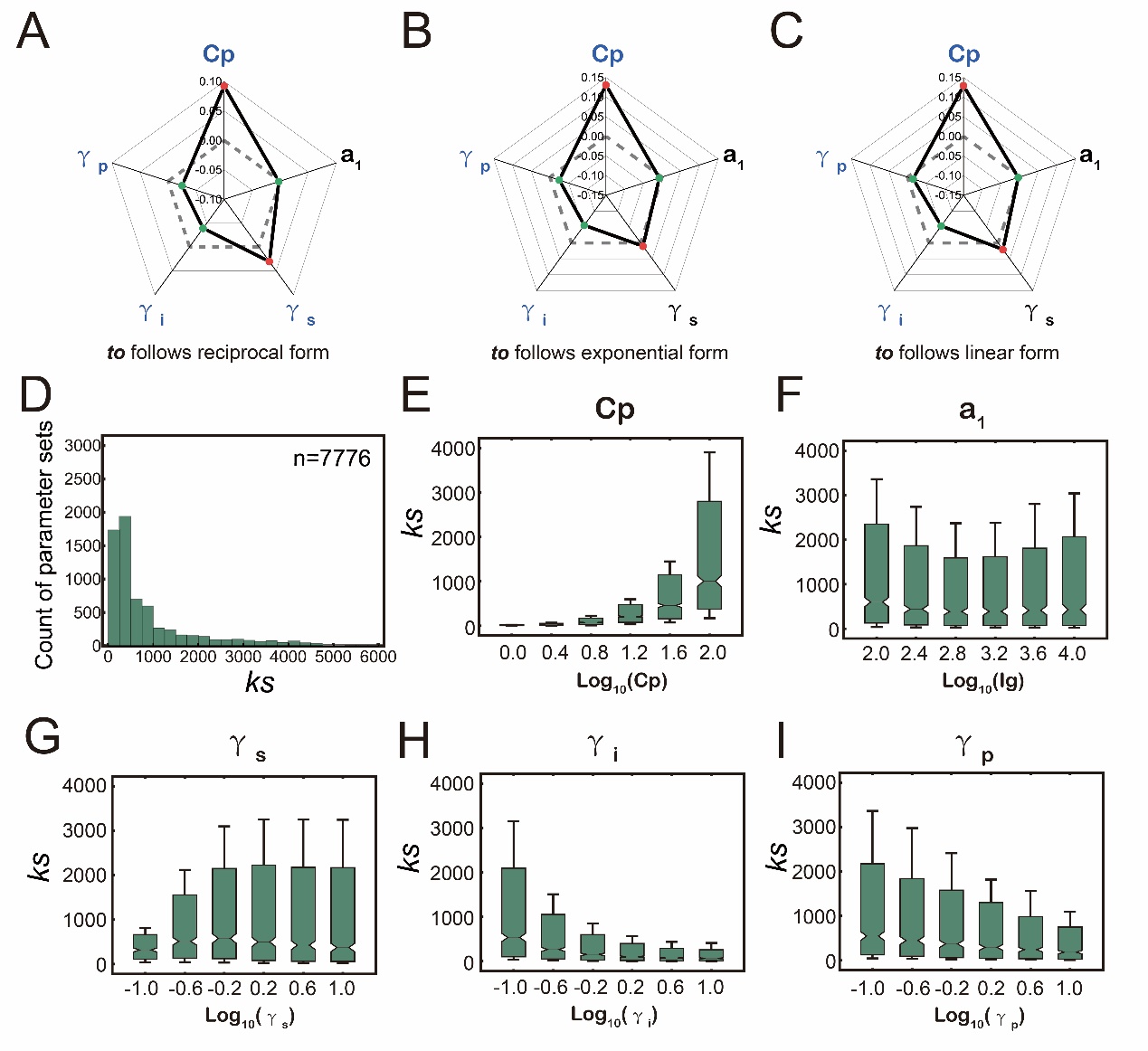

**Supplementary Figure 7** The effects of key parameters on the best fitting value of *ks* in presence of substrate toxicity. (A-C) Radar plot shows the values of fitting coefficients of the parameters from multiple linear regression analyses, in which toxic effect of substrate was assumed to follow three different forms. The data used here are listed in Supplementary Table 5. (D-I) Results when toxic effects following reciprocal form are shown in detail. (D) The distributions of best fitting value of *ks.* (E-I) The changes of best fitting value of *ks* with the different values of five key parameters.

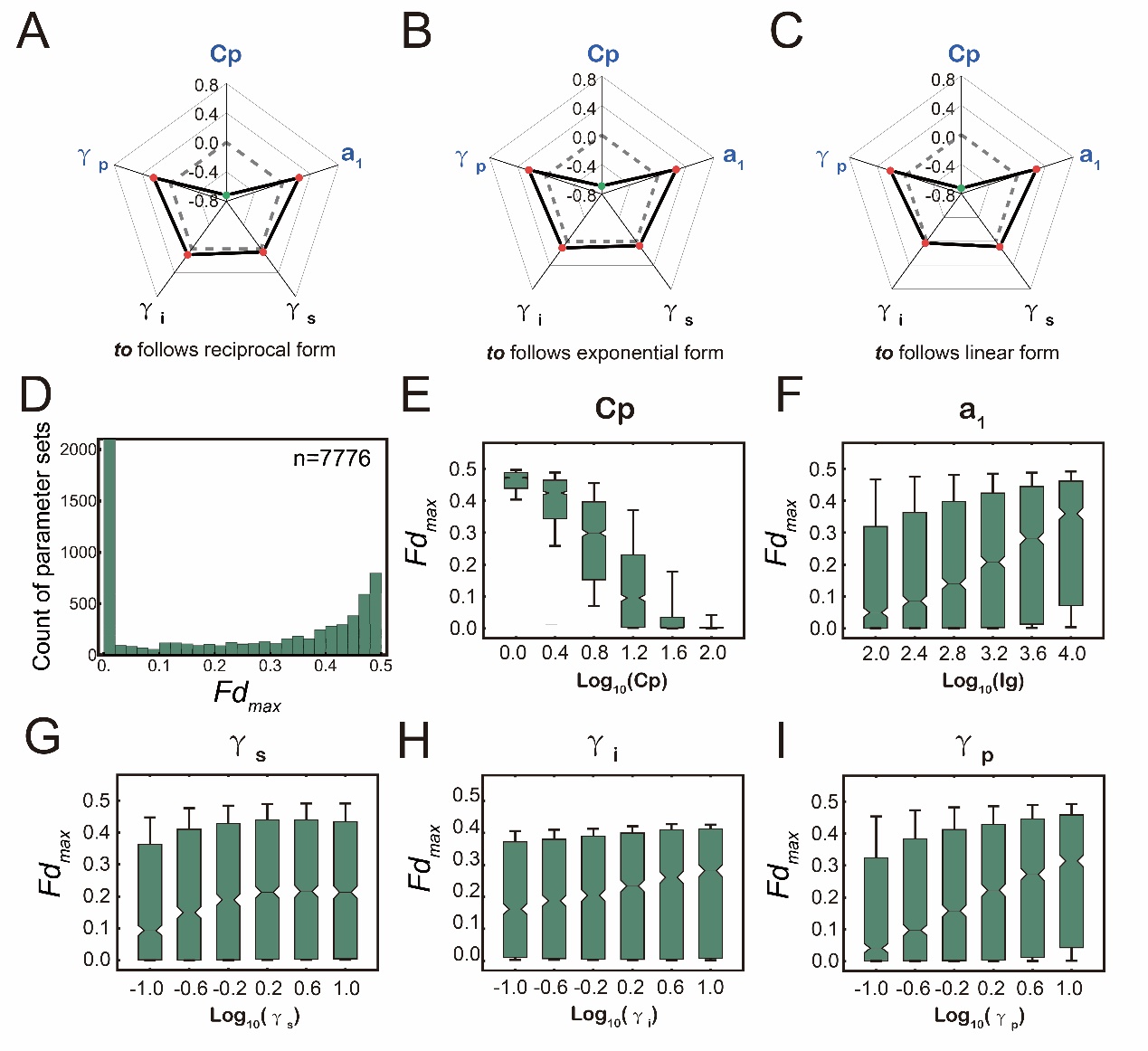

**Supplementary Figure 8** The effects of key parameters on the best fitting value of *Fd_max_* in presence of substrate toxicity. (A-C) Radar plot shows the values of fitting coefficients of the parameters from multiple linear regression analyses, in which toxic effect of substrate was assumed to follow three different forms. The data used here are listed in Supplementary Table 5. (D-I) Results when toxic effects following reciprocal form are shown in detail. (D) The distributions of best fitting value of *Fd_max_.* (E-I) The changes of best fitting value of *Fd_max_* with the different values of five key parameters.

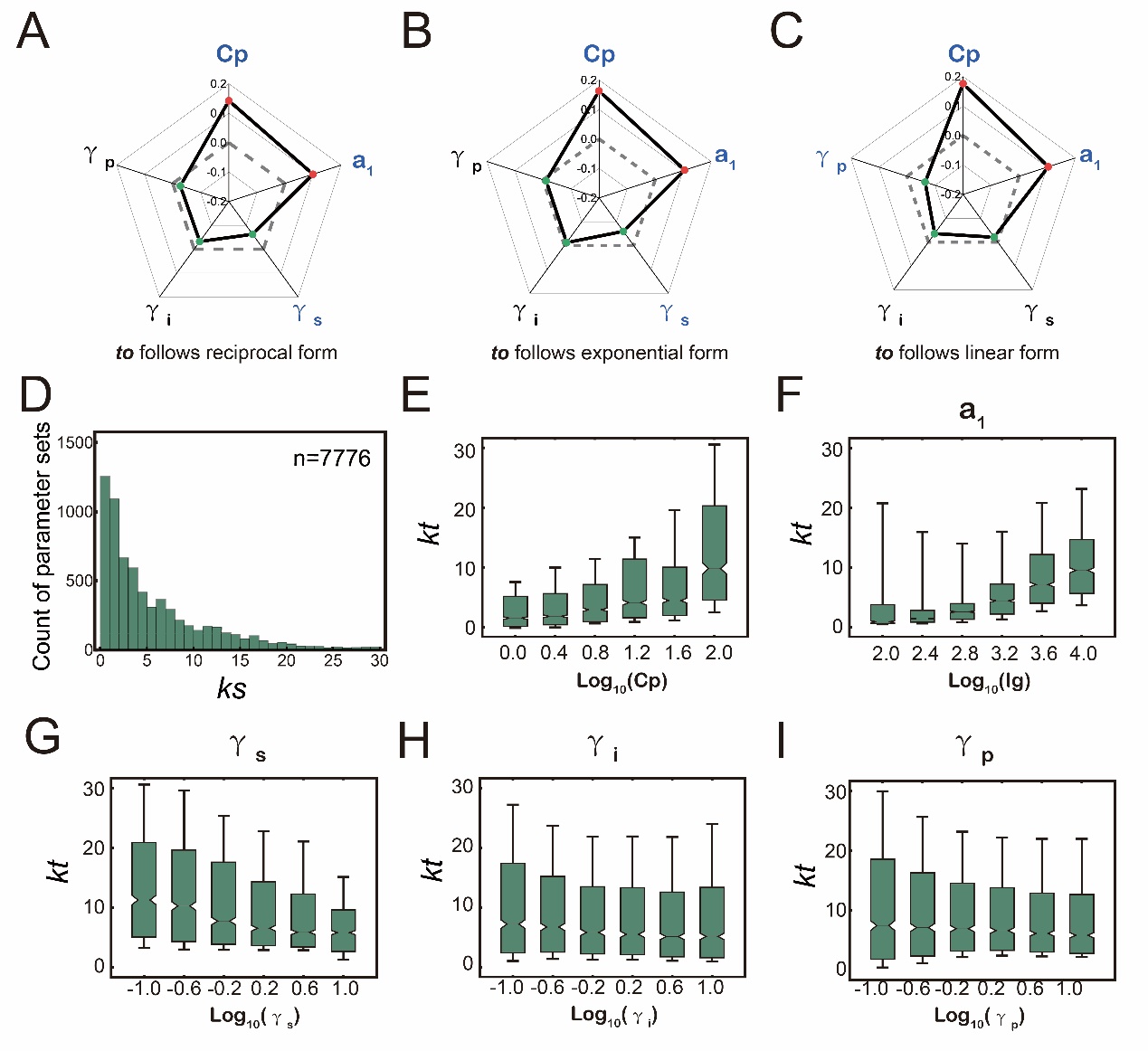

**Supplementary Figure 9** The effects of key parameters on the best fitting value of *kt*. (A-C) Radar plot shows the values of fitting coefficients of the parameters from multiple linear regression analyses, in which toxic effect of substrate was assumed to follow three different forms. The data used here are listed in Supplementary Table 5. (D-I) Results when toxic effects following reciprocal form are shown in detail. (D) The distributions of best fitting value of *kt.* (E-I) The changes of best fitting value of *kt* with the different values of five key parameters.

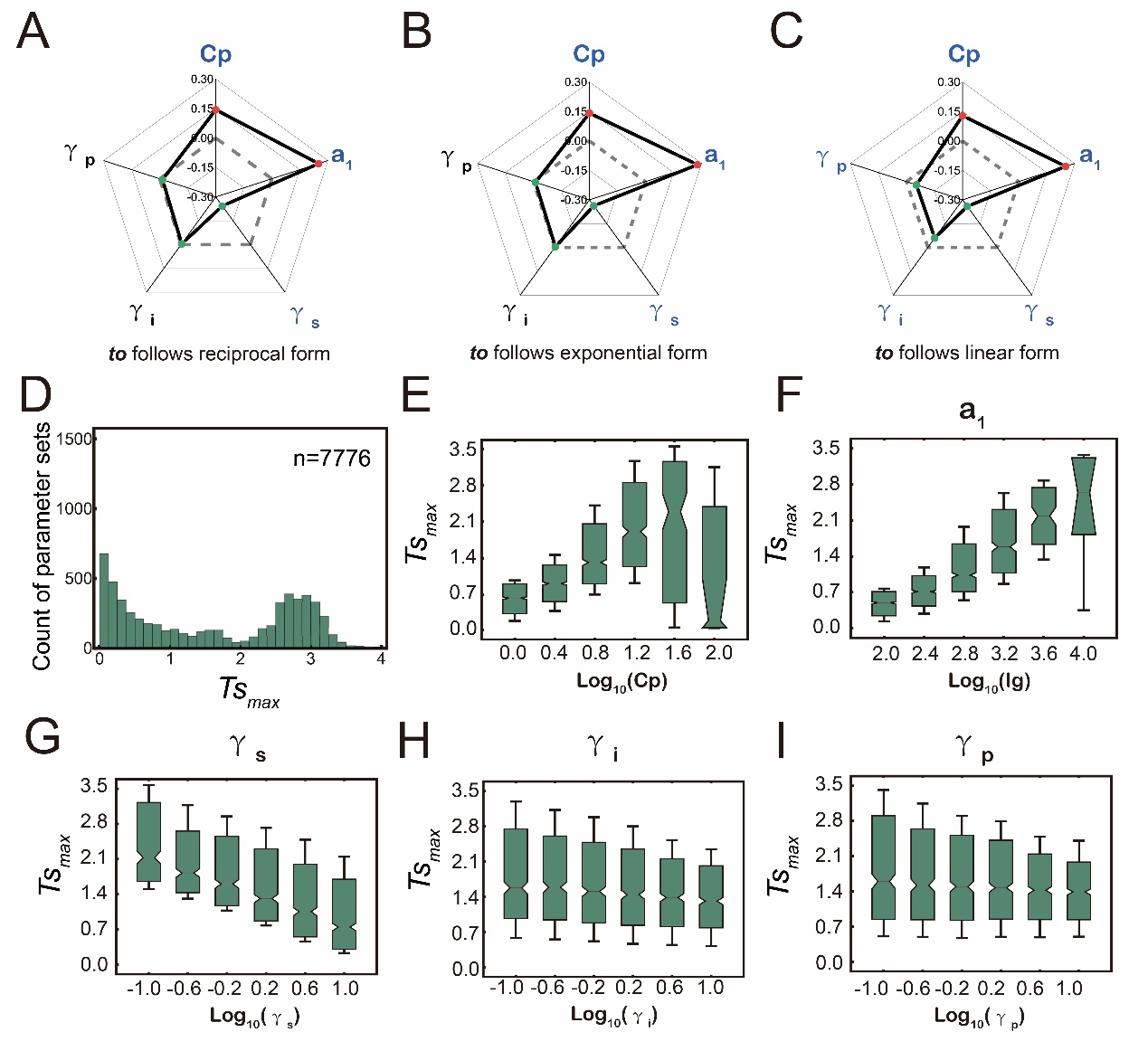

**Supplementary Figure 10** The effects of key parameters on the best fitting value of *Ts_max_*. (A-C) Radar plot shows the values of fitting coefficients of the parameters from multiple linear regression analyses, in which toxic effect of substrate was assumed to follow three different forms. The data used here are listed in Supplementary Table 5. (D-I) Results when toxic effects following reciprocal form are shown in detail. (D) The distributions of best fitting value of *Ts_max_.* (E-I) The changes of best fitting value of *Ts_max_* with the different values of five key parameters.

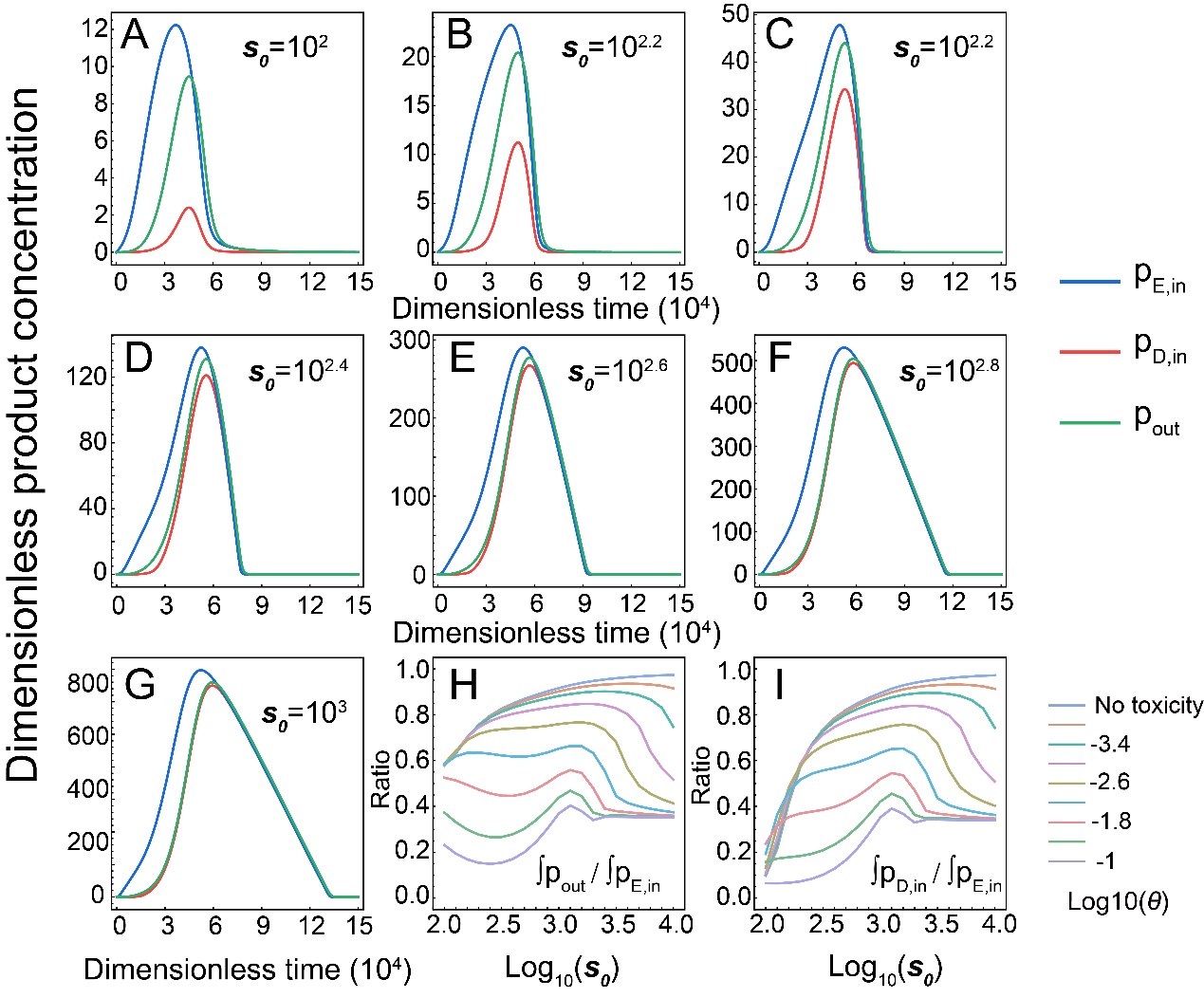

**Supplementary Figure 11** Dynamics of extra- and intracellular final product concentration of both populations in presence of substrate toxicity. (A-G) Dynamics of $\text{p}_{\text{D,in}}$, $\text{p}_{\text{E,in}}$, and $\text{p}_{\text{out}}$ in the simulations with different substrate concentration. In these simulations, toxic effect of substrate was assumed to follow reciprocal form and *θ* equals 0.001. (H) Shifts of integrated ratio of the extracellular to intracellular product concentration ($\int\text{p}_{\text{out}}/\text{p}_{\text{E,in}}$), across different initial substrate concentration and substrate toxicity. Ratio $\int\text{p}_{\text{out}}/\text{p}_{\text{E,in}}$ lower than 1 means majority of final product was privatized by the Embezzler cells, while higher ratio indicates more final product was released outside of cells. The plot indicates that higher substrate concentration leads to higher value of $\int\text{p}_{\text{out}}/\text{p}_{\text{E,in}}$ ratio, i.e., more release of final product. (I) Shifts of integrated ratio of the intracellular product concentration of Detoxifier population to that of the Embezzler population ($\int\text{p}_{\text{D,in}}/\text{p}_{\text{E,in}}$). Ratio$\int\text{p}_{\text{D,in}}/\text{p}_{\text{E,in}}$ lower than 1 means Embezzler accumulate more product inside the cell than Detoxifier. The increase of $\int\text{p}_{\text{D,in}}/\text{p}_{\text{E,in}}$ indicates this benefit was weakened. The plot indicates that higher substrate concentration leads to higher value of $\int\text{p}_{\text{D,in}}/\text{p}_{\text{E,in}}$, suggesting benefit from product privatization decreased. Parameter values used in this simulation: *y* =10^-4^, *Cp* = 10, *bg* = 1, *a_1_* = 10000, *a_2_* = 1000, *β_2_* = 1, $\text{γ}_{\text{s}}$ = 1, $\text{γ}_{\text{i}}$ =1, $\text{γ}_{\text{p}}$ = 1, *ρ =*1.

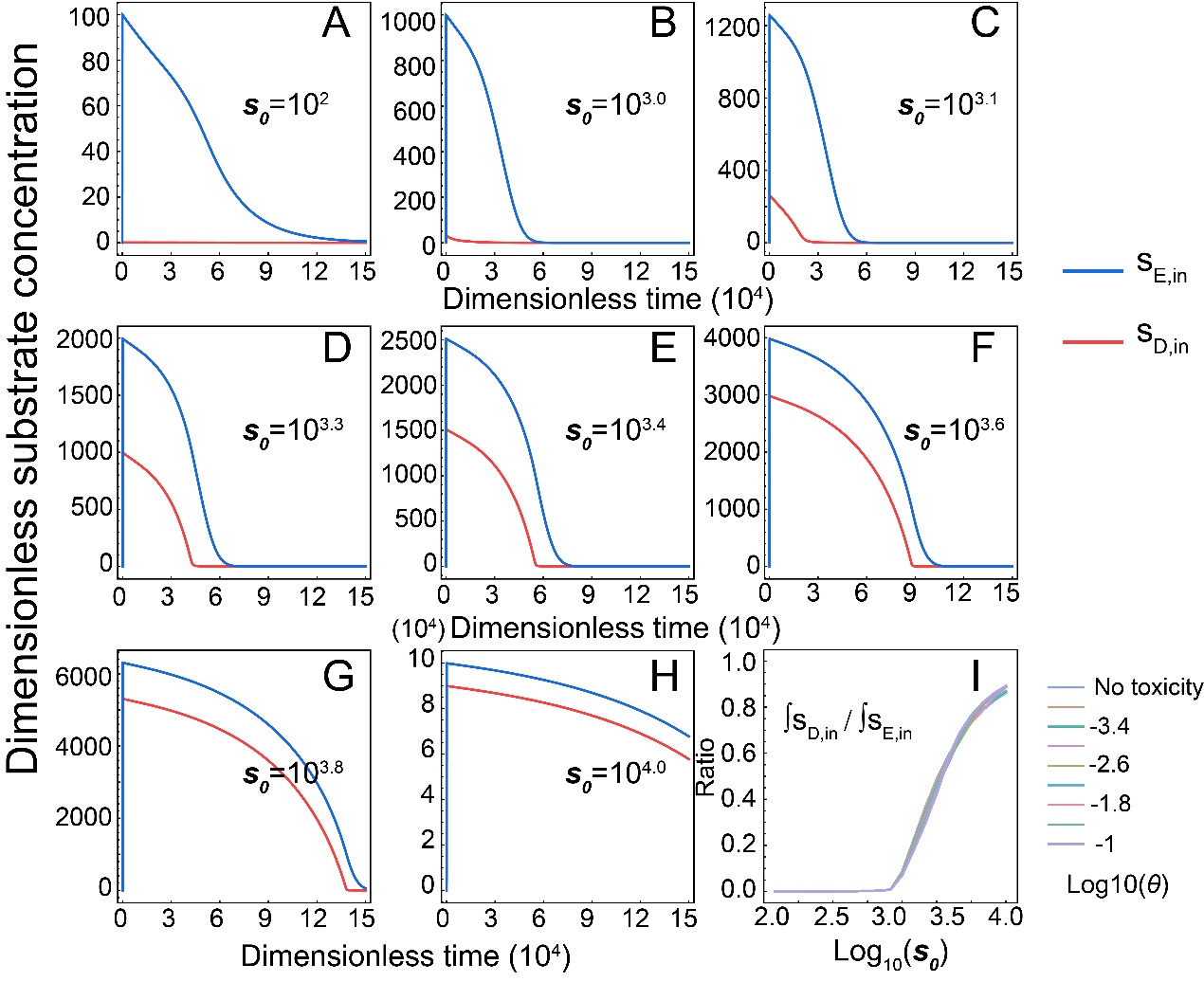

**Supplementary Figure 12** Dynamics of intracellular substrate concentration of both populations in presence of substrate toxicity. (A-H) Dynamics of $\text{s}_{\text{D,in}}$ and $\text{s}_{\text{E,in}}$ in the simulations with different substrate concentration. In these simulations, toxic effect of substrate was assumed to follow reciprocal form and *θ* equals 0.001. (I) Shifts of integrated ratio of the intracellular substrate concentration of Detoxifier population to that of the Embezzler population ($\int\text{s}_{\text{D,in}}/\text{s}_{\text{E,in}}$). Ratio$\int\text{s}_{\text{D,in}}/\text{s}_{\text{E,in}}$ lower than 1 means Detoxifier accumulate lower amount of substrate inside the cell than Embezzler. The plot indicates that the Detoxifier population possess lower intracellular substrate concentration than the Embezzler population, showing potential benefit when the substrate is toxic. Parameter values used in this simulation: *y* =10^-4^, *Cp* = 10, *bg* = 1, *a_1_* = 10000, *a_2_* = 1000, *β_2_* = 1, $\text{γ}_{\text{s}}$ = 1, $\text{γ}_{\text{i}}$ =1, $\text{γ}_{\text{p}}$ = 1, *ρ =*1.

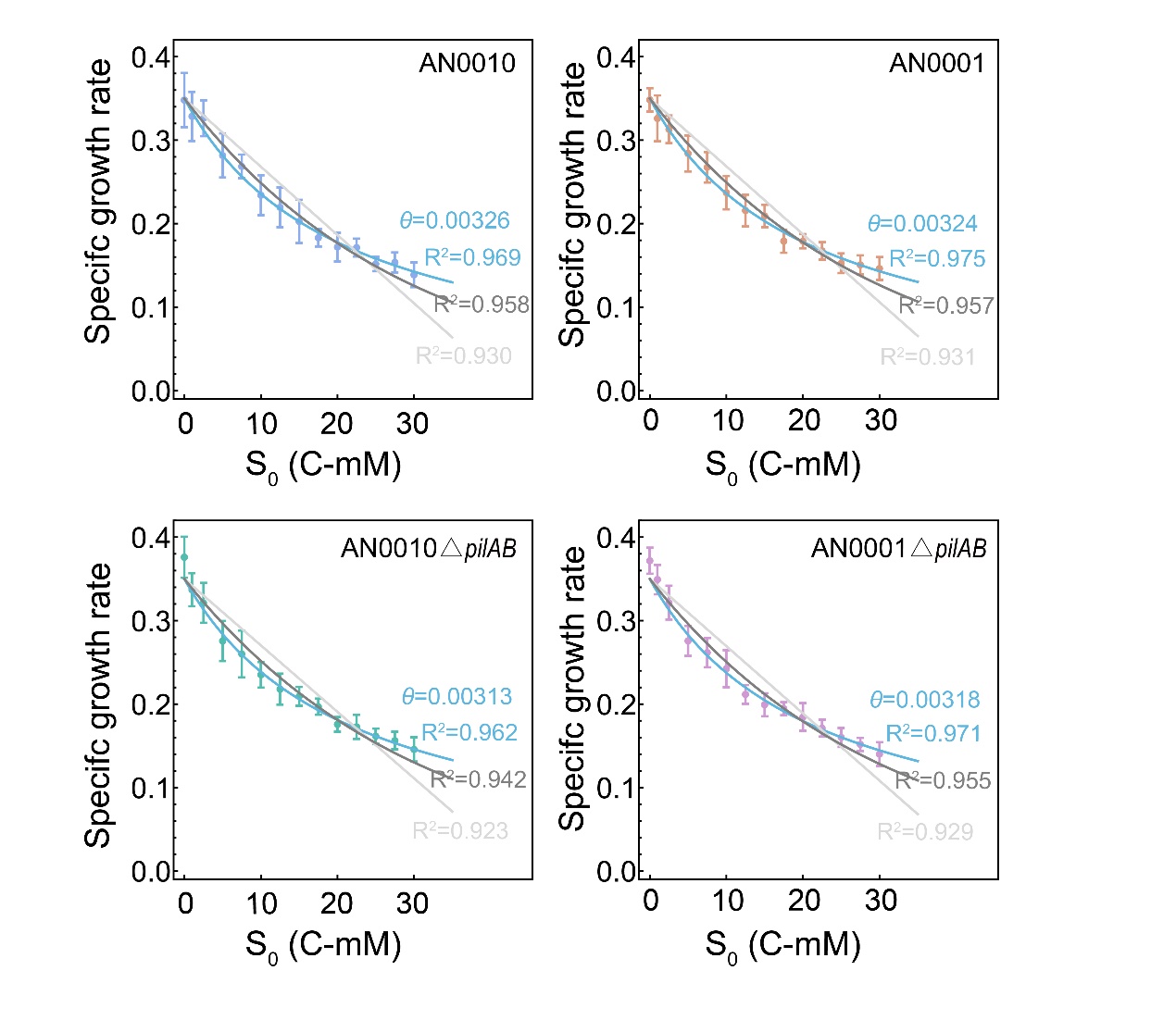

**Supplementary Figure 13** Determination of the toxic strength of the substrate salicylate. We measured the growth rate of four strains used in this study supplemented with pyruvate, one of the end products, as the carbon source, and adding different amount of salicylate. The growth rates were fitted to salicylate concentration, using a formula modified from logistic function and considering three different forms of toxic terms (See Supplementary Information S3.4 for detailed description). According to the value of Adjusted R^2^, the reciprocal-form toxic term fitted best with the data. Thus, the value of toxic strength (*θ*) was then obtained, and used for subsequent mathematical prediction.

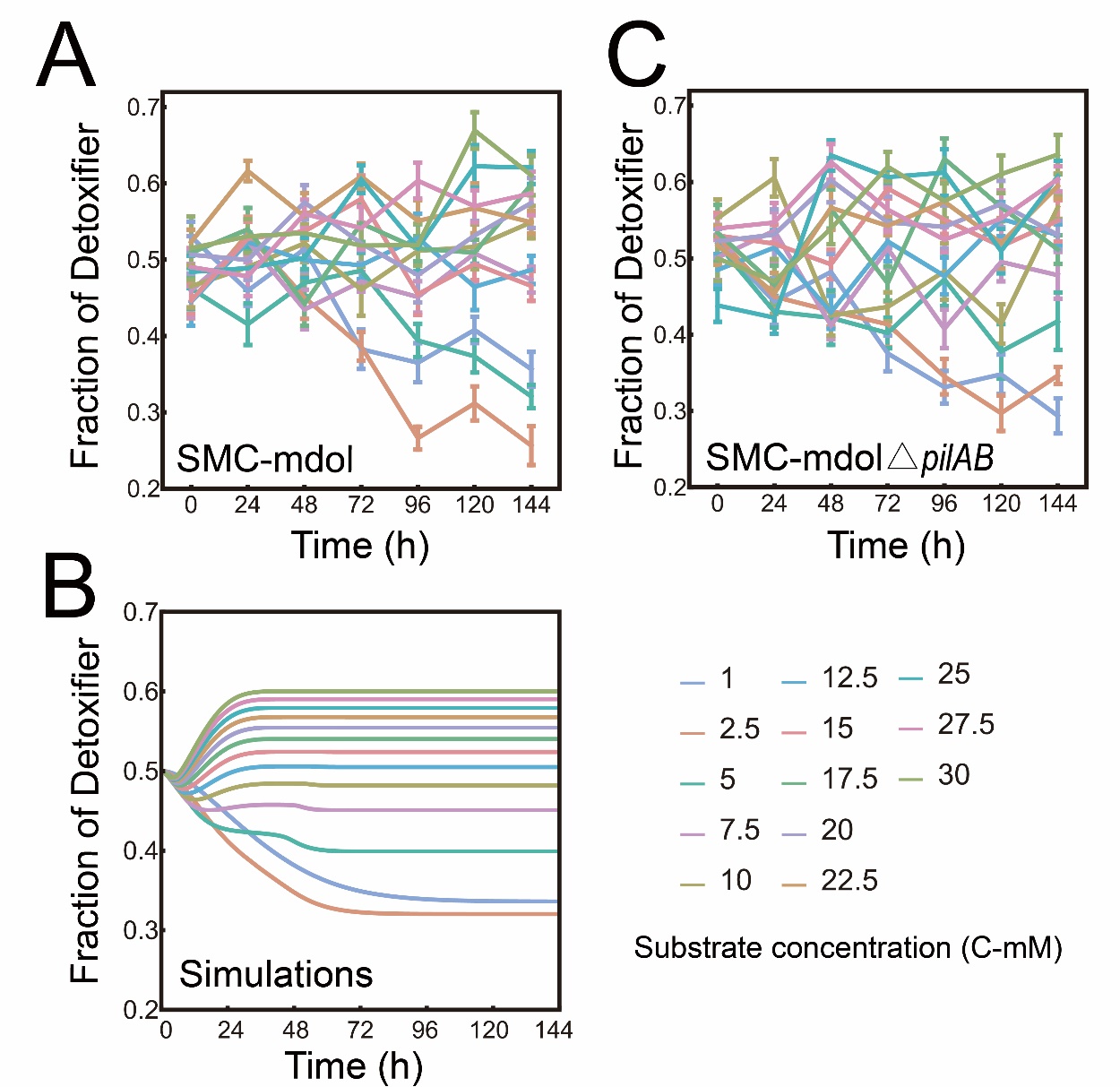

**Supplementary Figure 14** Dynamics of the steady-state fraction of Detoxifier population in the liquid culturing experiment of SMC-mdol (A) and SMC-mdol△*pilAB* (B), as well as the associated mathematical simulations (C).

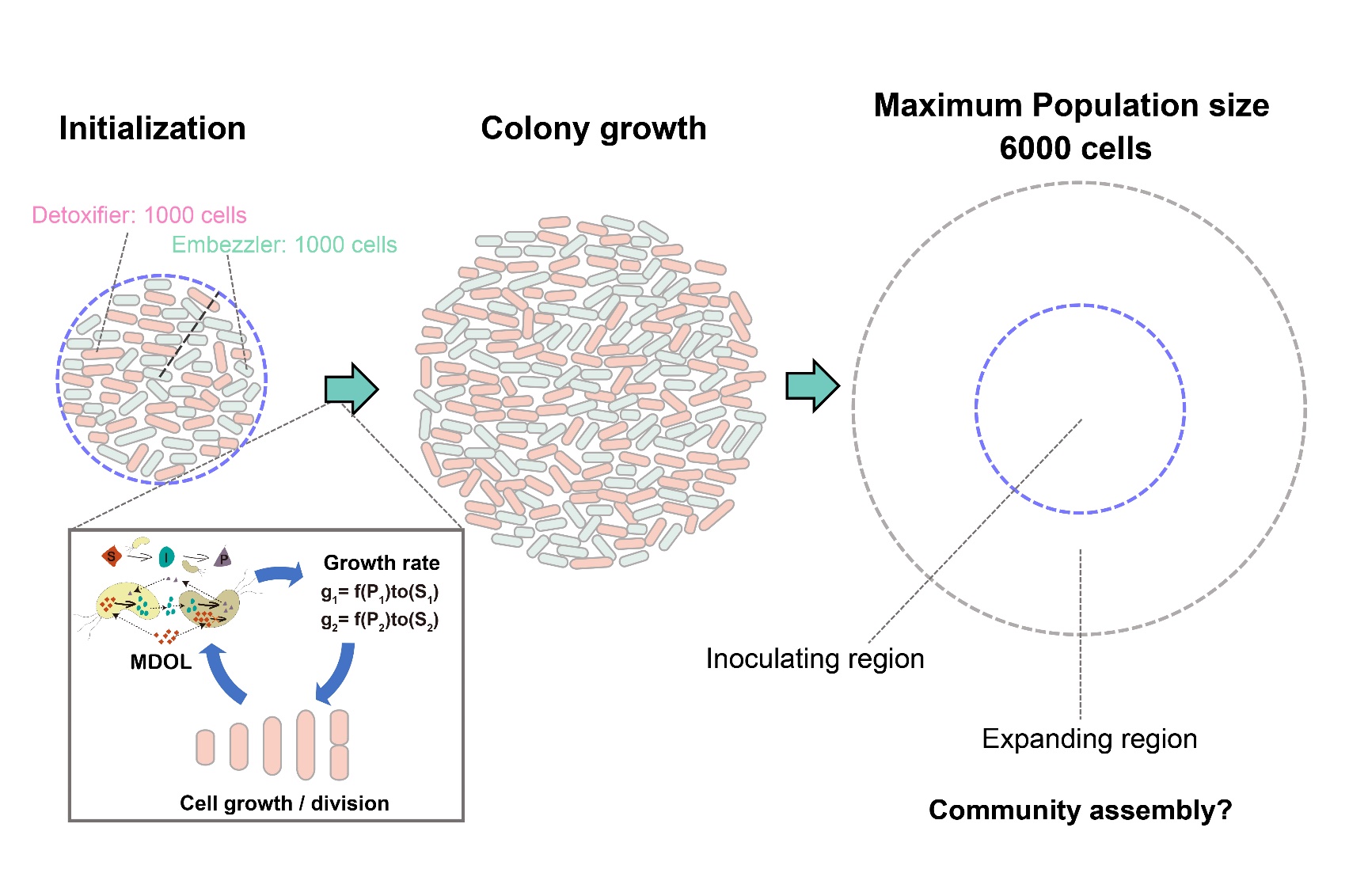

**Supplementary Figure 15** The logic of the individual-based model. Individual-based simulations were performed on a 2D plane. Bacterial cells were characterized as rigid capsules of variable length and fixed radius. Cells of the two populations were initialized in an inoculating ring with a radius of 300 μm, and 1000 cells for each population were randomly distributed. They grew and divided in this habitat by division of labor, driving the range expansion of the colony. By sequentially updating the cell configuration according to the rules defined by Eqn. [S1-S13], we simulated the development of colony structure until the total number of cells reaches 6000. We focused on the assembly of this community in the entire colony, the inoculating region (where cells are initiated, denoted by blue dashed circle) and expanding region of the colonies (the region between blue circle and gray circle).

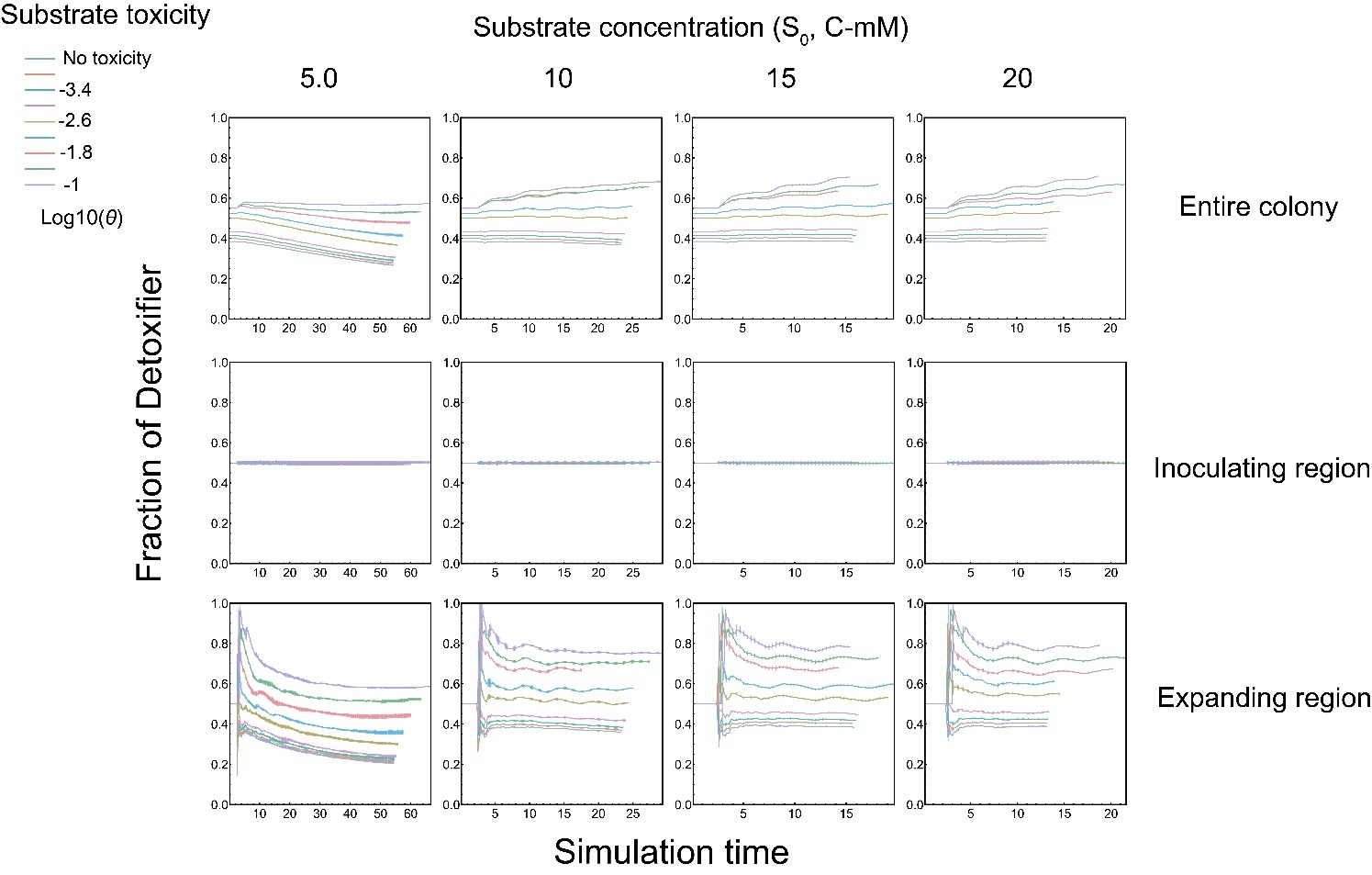

**Supplementary Figure 16** Temporal dynamics of community composition in the entire colony, inoculating region and expanding region of the colonies derived from our individual-based (IB) simulations. Plot suggests that the community composition in the inoculating region rarely change with time, but the community composition in the expanding region significantly shifted with time and gradually reaches a steady-state.

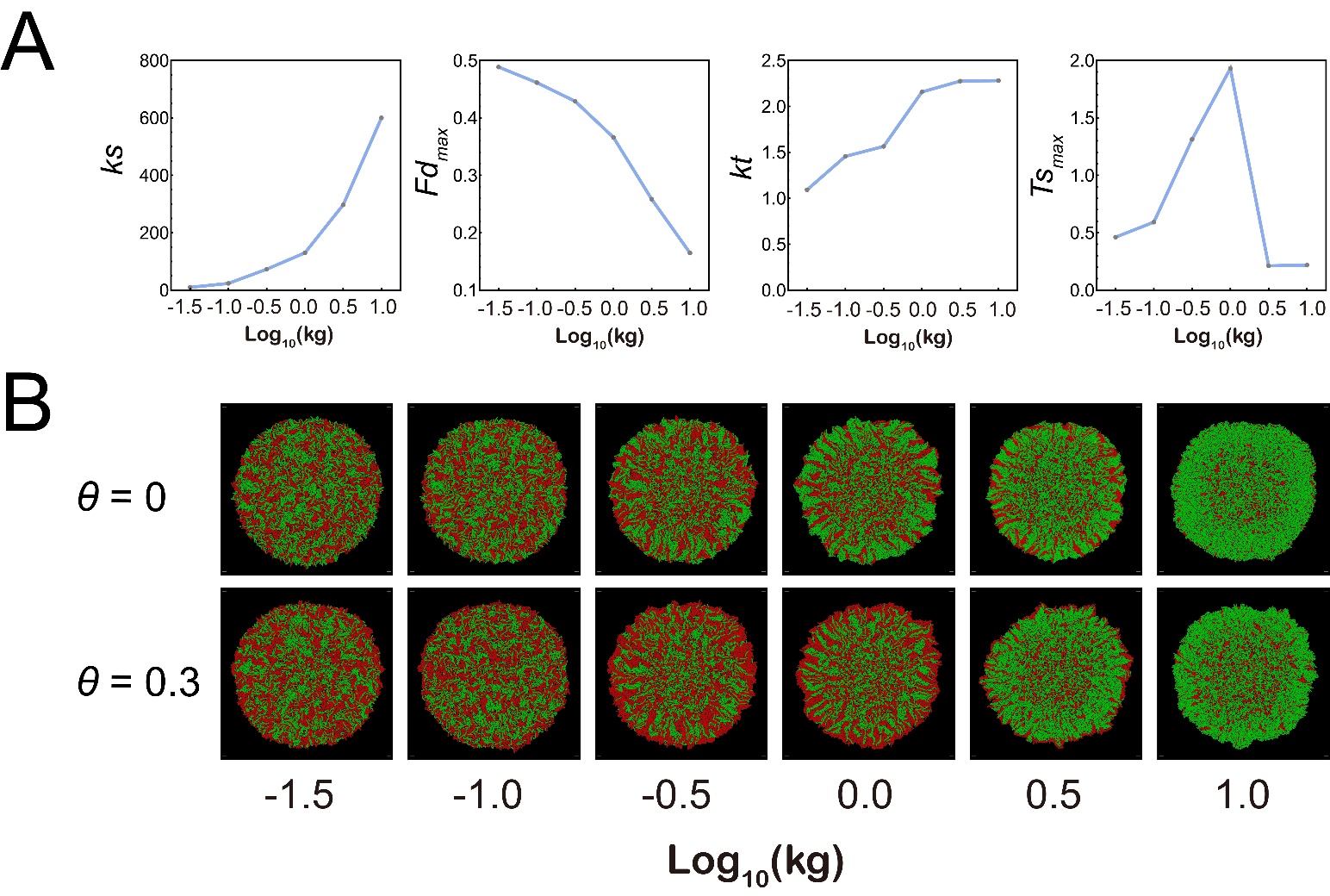

**Supplementary Figure 17** The effect of the consuming rate of P on the best fitting value of *ks*, *Fd_max_*, *kt*, and *Ts_max_* in our individual-based simulations. (A) The relationship between the best fitting value of *ks*, *Fd_max_*, *kt*, and *Ts_max_* with the consuming rate of P. In these simulations, the consuming rate of P was modulated by adjusting the value of *kg* (Supplementary Table 7). In the simulations of each given value of *kg*, eight different substrate concentration (*s_0_*) and nine different toxic strength of substrate (*θ*) were applied. Other parameters were initialized with the default values shown in Supplementary Table 7. Then the simulations data were fitted to Eqn. [2] to obtain the best fitting value of *ks*, *Fd_max_*, *kt*, and *Ts_max_*. The Adjust R^2^ values for these fitting analyses range from 0.984 to 0.999. (B) Representative colony images at steady-state obtained from individual-based simulations initialized with different level of the consuming rate of P. Shown are the results in which S0 was set to 10 C-mol/L. Detoxifier cells are shown in red, while Embezzler cells are shown in green.

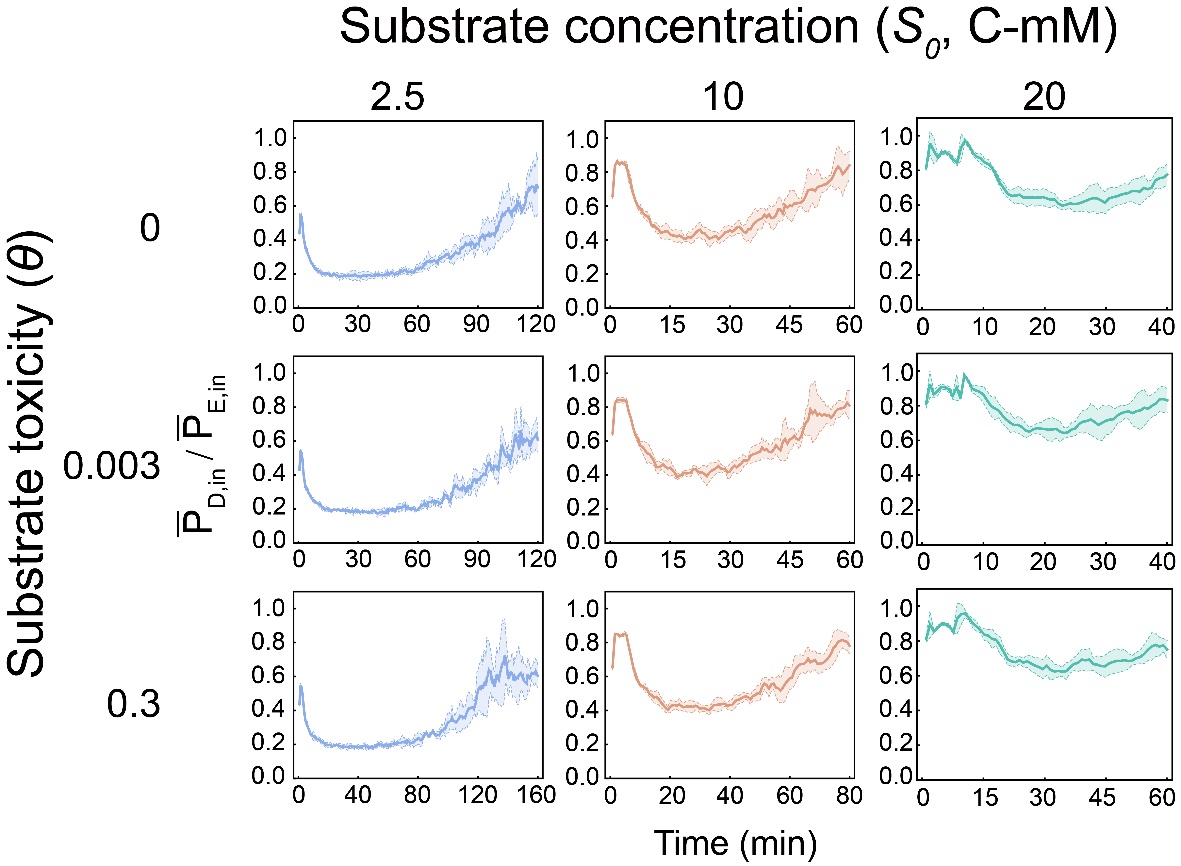

**Supplementary Figure 18** Dynamics of intracellular final product concentration of the cells from both populations in our individual-based simulation considering different conditions of substrate concentration and toxicity. $\bar{\text{P}_{\text{D,in}}}\text{/}\bar{\text{P}_{\text{E,in}}}$ was defined as the ratio of average the intracellular product concentration of all Detoxifier cells to that of the Embezzler cells. The ratio lower than 1 means Embezzler cells accumulate more product inside the cell than Detoxifier. The increase of the ratio indicates this benefit was weakened. The plot indicates that higher substrate concentration leads to higher value of $\bar{\text{P}_{\text{D,in}}}\text{/}\bar{\text{P}_{\text{E,in}}}$, suggesting benefit from product privatization decreased. Shown are the results using the same parameters as in Figure 5A.

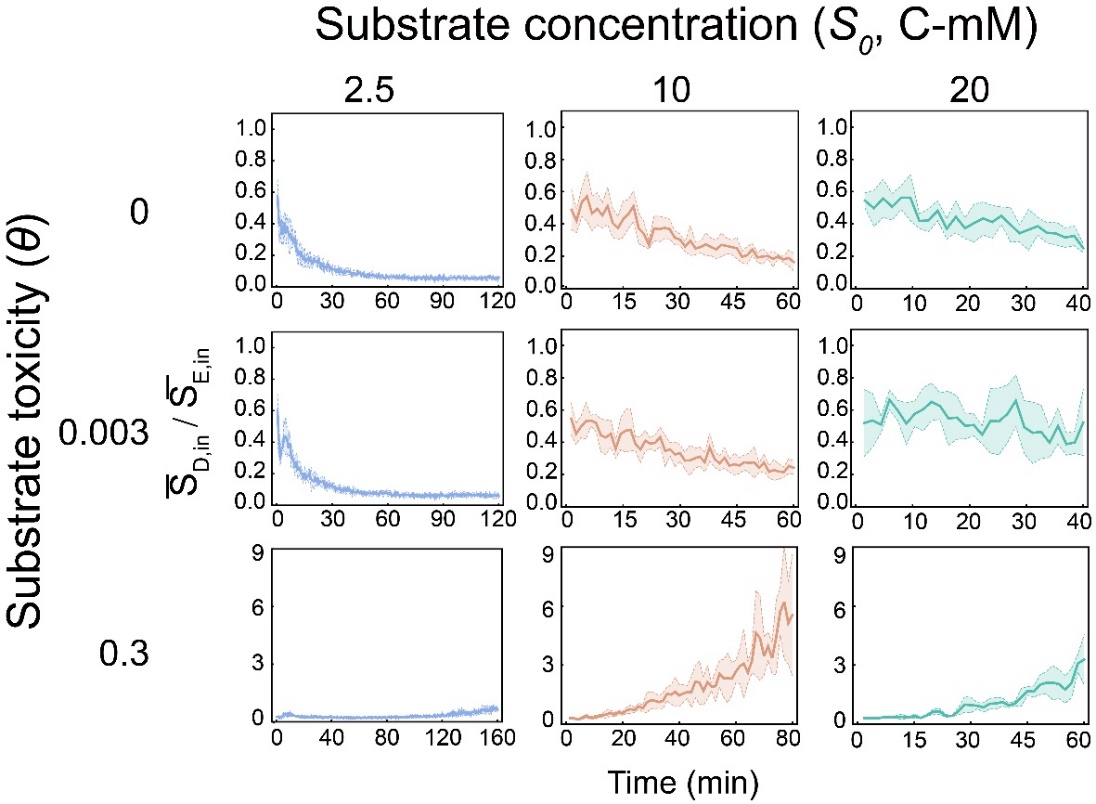

**Supplementary Figure 19** Dynamics of intracellular substrate product concentration the cells from of both populations in our individual-based simulation considering different conditions of substrate concentration and toxicity. $\bar{\text{S}_{\text{D,in}}}\text{/}\bar{\text{S}_{\text{E,in}}}$ was defined as the ratio of average the intracellular substrate concentration of all Detoxifier cells to that of the Embezzler cells. The ratio lower than 1 means Detoxifier cells accumulate lower amount of substrate inside the cell than Embezzler. The plot indicates that Detoxifier cells possess lower intracellular substrate concentration than Embezzler cells when substrate toxicity is relative lower, thus get potential benefit when the substrate is toxic. However, when substrate toxicity is higher, Detoxifier cells possess higher intracellular substrate concentration than Embezzler cells. This was because of the asymmetric distribution of the two populations in the developed pattern (Supplementary Figure 28). In this pattern, the Detoxifier cells occupied the place at the frontiers of the colony, where the substrate concentration is much higher. Shown are the results using the same parameters as in Figure 5A.

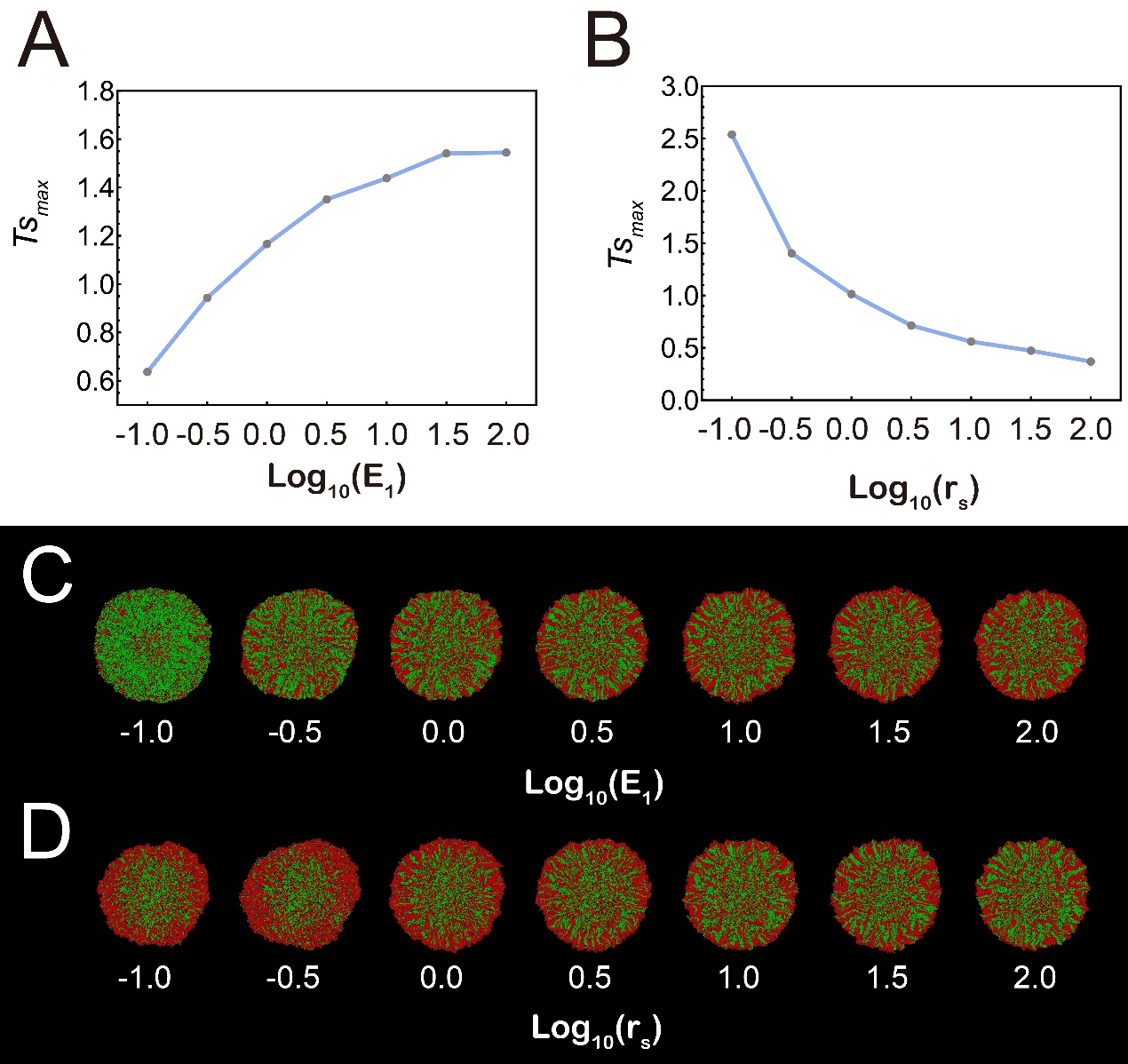

**Supplementary Figure 20** The effects of the speed of the first reaction and transport rate of S on the best fitting value of *Ts_max_* in our individual-based simulations. (A-B) The relationship between the best fitting value of *Ts_max_* with the speed of the first reaction (A) and transport rate of S (B). In these simulations, the speed of the first reaction was modulated by adjusting the value of the concentration of the first enzyme (E1) and transport rate of S was modulated by adjusting *r_s_* (Supplementary Table 7). In the simulations of each given value of E1 or *r_s_*, eight different substrate concentration (*s_0_*) and nine different toxic strength of substrate (*θ*) were applied. Other parameters were initialized with the default values shown in Supplementary Table 7. Then the simulations data were fitted to Eqn. [2] to obtain the best fitting value of *Ts_max_*. The Adjust R^2^ values for these fitting analyses range from 0.986 to 0.998. (C-D) Representative colony images at steady-state obtained from individual-based simulations initialized with different level of the speed of the first reaction (C) and transport rate of S (D). Shown are the results in which S0 was set to 10 C-mol/L and *θ* was 0.03. Detoxifier cells are shown in red, while Embezzler cells are shown in green.

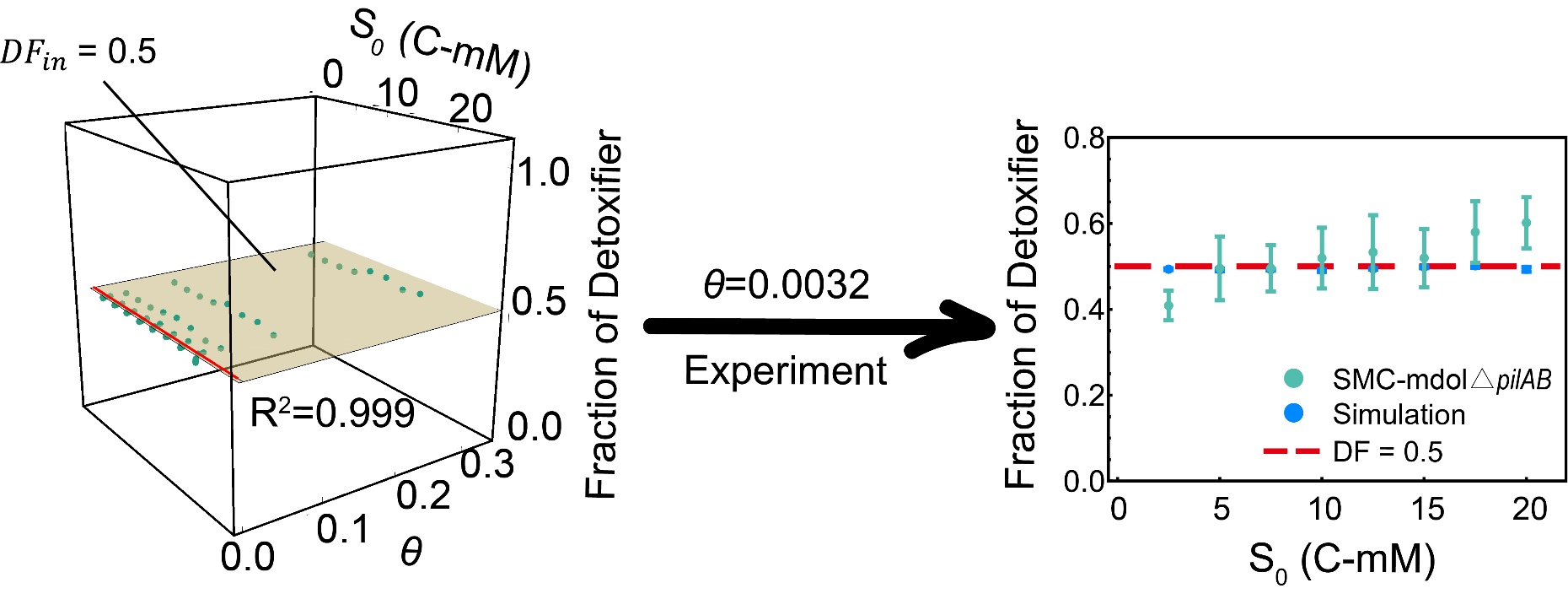

**Supplementary Figure 21** Analyses of community composition in the inoculating region of the colonies derived from our individual-based (IB) simulations (left) and experiments using SMC-mdol△*pilAB* (Right). Plot suggests that substrate concentration and its toxicity rarely affect the stead-state fraction of Detoxifier in the inoculating region. Left: The green dots denote the stead-state fraction of Detoxifier in the inoculating region from IB simulations. The surface shows the plot of the best fitting function using Eqn. [2]. The Red line in the surface denotes the scenarios *θ*=0.0032, which is the toxic strength of salicylate obtained from experiemental measurements. Right: Comparation between the results obtained from experimental measures and those from IB simulations.

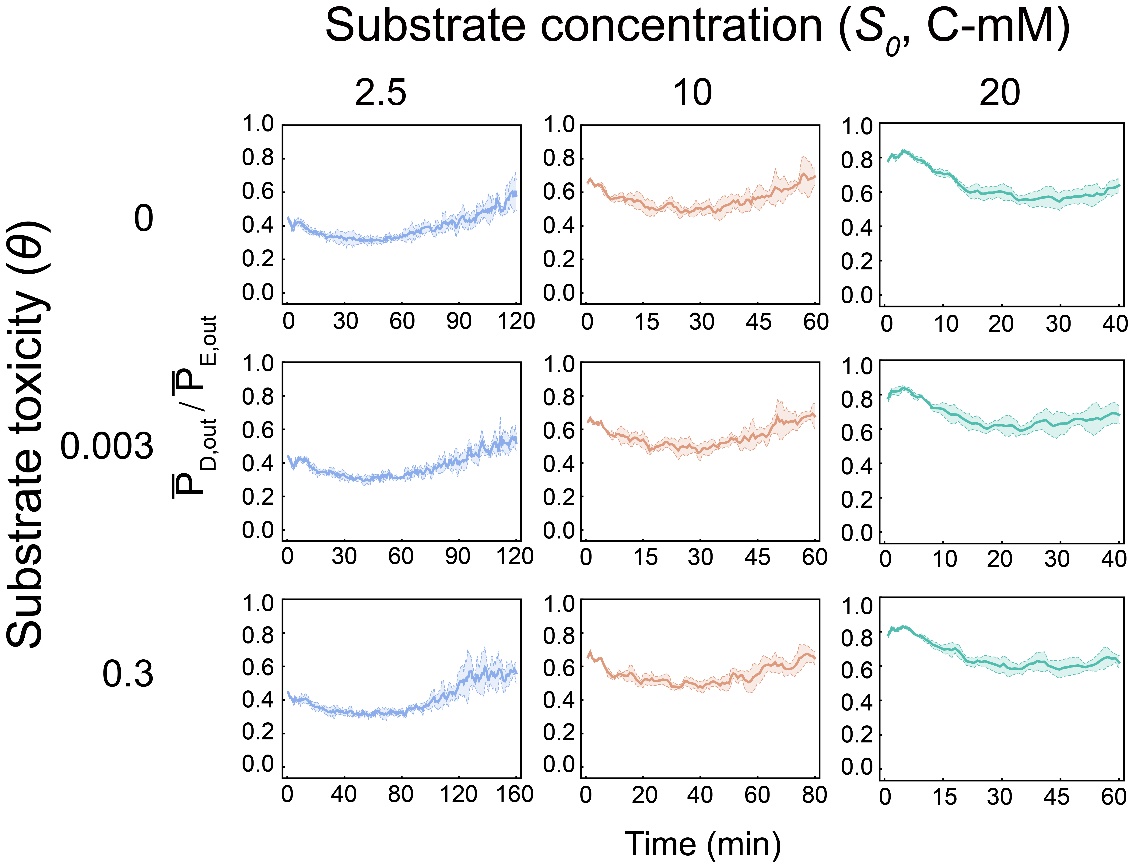

**Supplementary Figure 22** Dynamics of local extracellular final product concentration the cells from of both populations in our individual-based simulation considering different conditions of substrate concentration and toxicity. $\bar{\text{P}_{\text{D,out}}}\text{/}\bar{\text{P}_{\text{E,out}}}$ was defined as the ratio of average the local extracellular product concentration (that is, the concentration in the grid containing the cell) of all Detoxifier cells to that of the Embezzler cells. The ratio lower than 1 means product tends to be distributed more around Embezzler cells than Detoxifier. The plot indicates that Embezzler cells possess higher local extracellular product concentration than Detoxifier. Shown are the results using the same parameters as in Figure 5A.

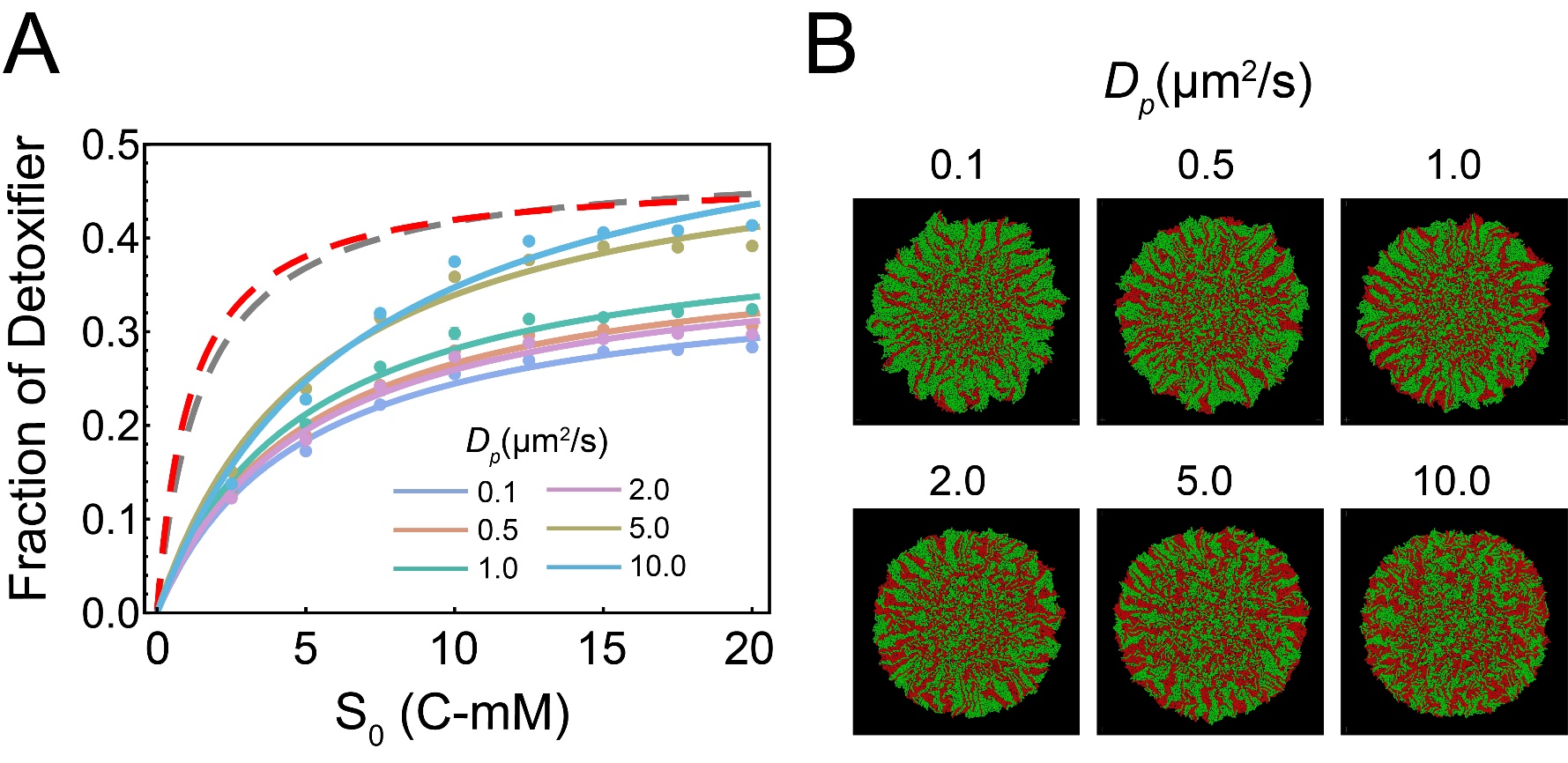

**Supplementary Figure 23** The effects of diffusivity of final product on the assembly of microbial community engaged in metabolic division of labor. (A) The relationship between initial substrate concentration (*S_0_*) with the steady-state frequency of Detoxifier cells in the expanding region of the colonies under different diffusion level of final product (*D_p_*). In these simulations, substrate toxicity (*θ*) was set to be zero. Other parameters were initialized with the default values shown in Supplementary Table 7. The simulations results are shown by the dots. The data were then fitted to Eqn. [1] to obtain the curves shown in the plot. The Adjust R^2^ values for these fitting analyses range from 0.996 to 0.999. (B) Representative colony images at steady-state obtained from individual-based simulations initialized with different diffusion level of final product. Shown are the results in which S0 was set to 10 C-mol/L. Detoxifier cells are shown in red, while Embezzler cells are shown in green.

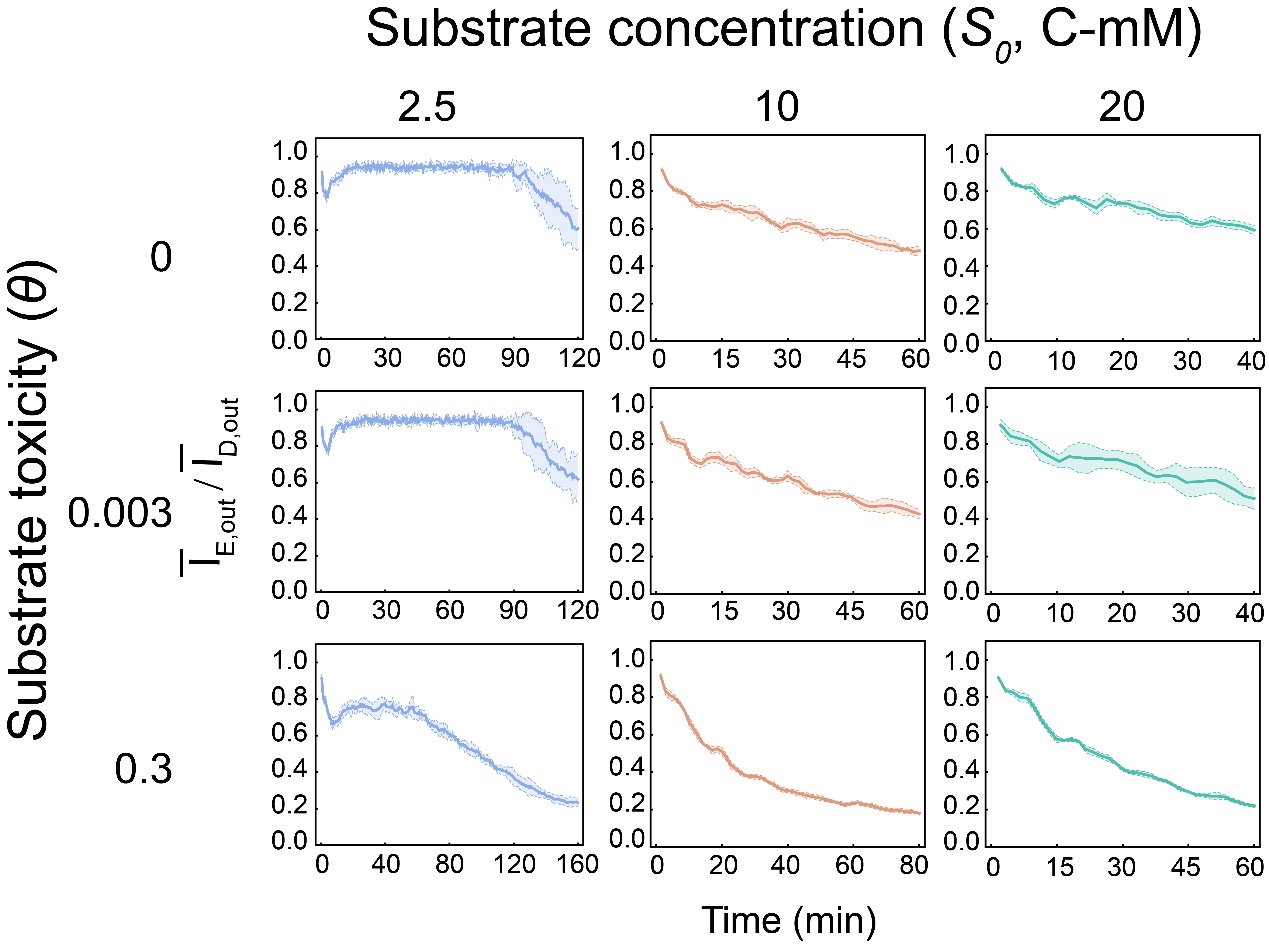

**Supplementary Figure 24** Dynamics of local extracellular intermediate concentration the cells from of both populations in our individual-based simulation considering different conditions of substrate concentration and toxicity. $\bar{\text{I}_{\text{E,out}}}\text{/}\bar{\text{I}_{\text{D,out}}}$ was defined as the ratio of average the local extracellular intermediate concentration (that is, the concentration in the grid containing the cell) of all Embezzler cells to that of the Detoxifier cells. The ratio lower than 1 means intermediate tends to be distributed more around Detoxifier cells than Embezzler. The plot indicates that Detoxifier cells possess higher local extracellular intermediate concentration than Detoxifier. Shown are the results using the same parameters as in Figure 5A.

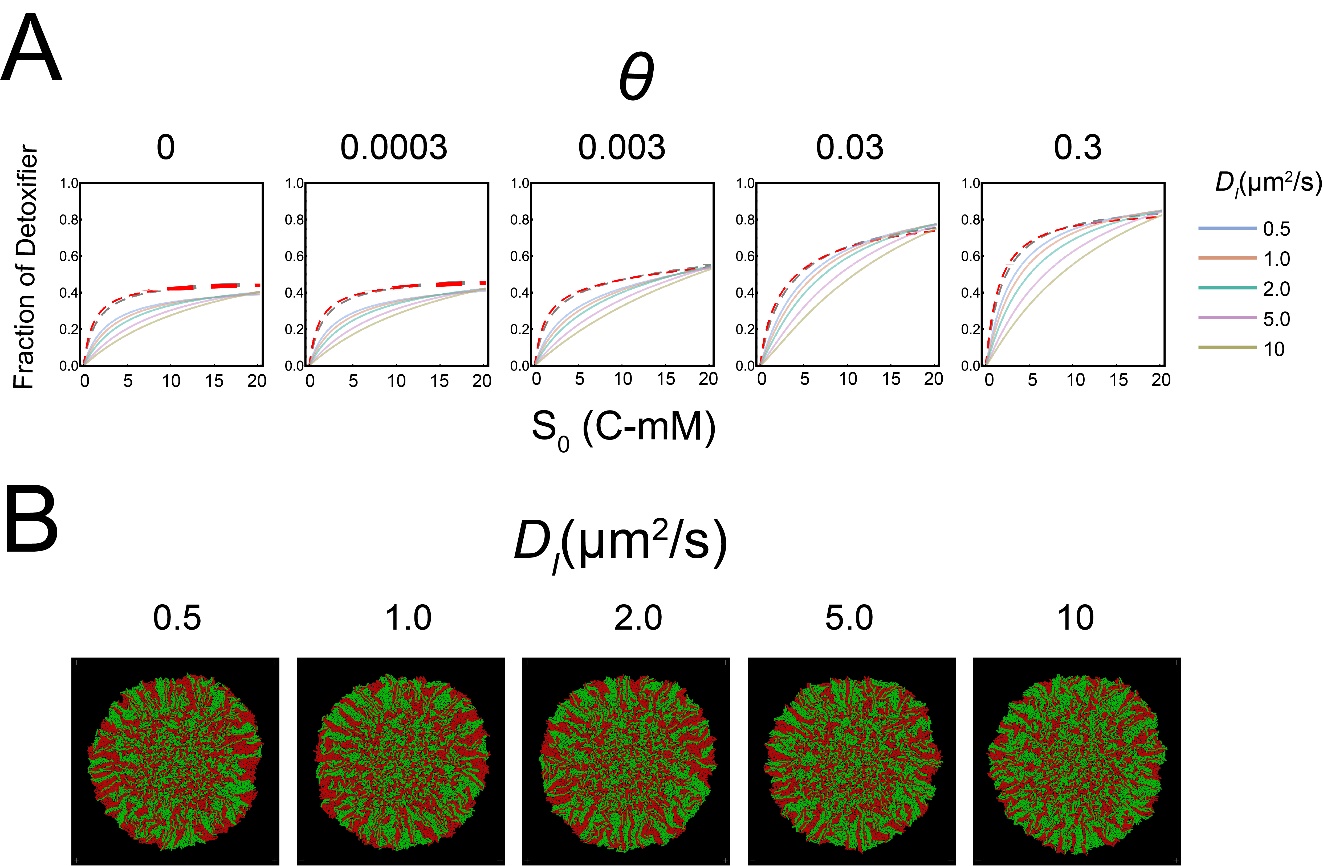

**Supplementary Figure 25** Diffusion level of intermediate affects the assembly of microbial community engaged in metabolic division labor. (A) The relationship between initial substrate concentration (*S_0_*) with the steady-state frequency of Detoxifier cells in the expanding region of the colonies, across different diffusion level of intermediates (*D_i,_* denoted by different curve colors) and different strength of substrate toxicity (*θ*, denoted by five subgraphs). Other parameters in these simulations were initialized with the default values shown in Supplementary Table 7. The simulation data were then fitted to Eqn. [2] to obtain the curves shown in the plot. The Adjust R^2^ values for these fitting analyses range from 0.993 to 0.997. (B) Representative colony images at steady-state obtained from individual-based simulations initialized with different diffusion level of intermediate. Shown are the results in which *S_0_* was set to 10 C-mol/L and *θ* was 0.03. In the colony images, Detoxifier cells are shown in red, while Embezzler cells are shown in green.

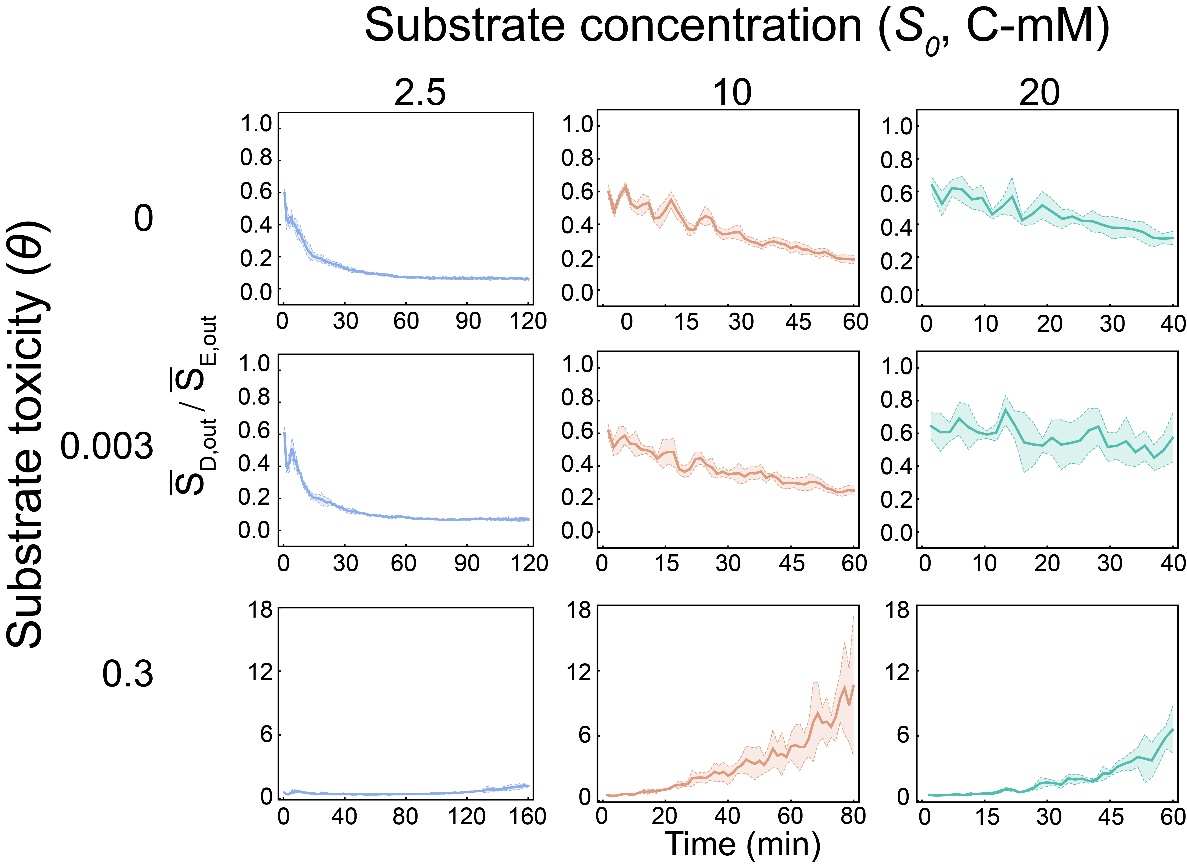

**Supplementary Figure 26** Dynamics of local extracellular substrate concentration the cells from of both populations in our individual-based simulation considering different conditions of substrate concentration and toxicity. $\bar{\text{S}_{\text{D,out}}}\text{/}\bar{\text{S}_{\text{E,out}}}$ was defined as the ratio of average the local extracellular substrate concentration (that is, the concentration in the grid containing the cell) of all Detoxifier cells to that of the Embezzler population. The ratio lower than 1 means local substrate concentration around Detoxifier cells is higher than Embezzler. The plot indicates that Detoxifier cells possess lower local extracellular substrate concentration than that of the Embezzler cells when substrate toxicity is absent or at a relative lower level. However, when strength of substrate toxicity is sufficient high (e.g., *θ* = 0.3), the local extracellular substrate concentration around Detoxifier cells becomes higher than that of the Embezzler cells, due to the development of the specific ‘Detoxifier-leading’ pattern. Shown are the results using the same parameters as in Figure 5A.

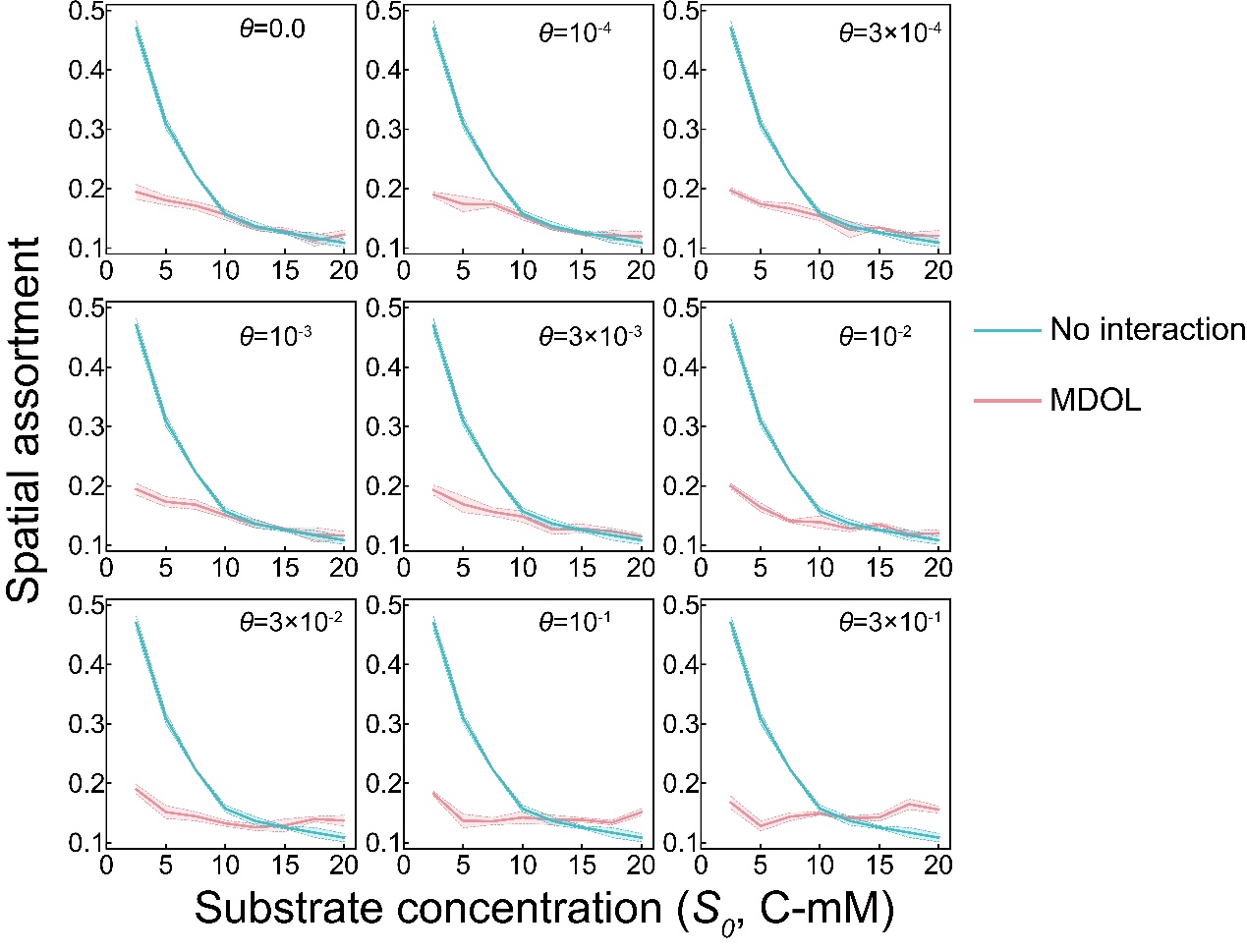

**Supplementary Figure 27** Substrate traits affect the intermixing level of the spatial pattern developed by microbial community engaged in metabolic division of labor (MDOL). As a control, we built an additional model to simulate the scenarios where the two populations just competed for the limited resource (without MDOL interaction). We did not include S and I in this model, while supplied P at the beginning which was evenly distributed across the environments. Values of the parameters used in this model were also totally same as before. Spatial assortment was defined to assess the level of spatial mixing between the populations involved in the community, which is calculated using the same method described previously [34, 35] and the distance of 4 μm was used as the neighborhood distance. Higher value of spatial assortment means the pattern possess lower intermixing level. These simulations suggest that, compared with the non-interaction scenario, the patterns developed by MDOL community possess higher intermixing level when both substrate concentration and its toxic strength are relatively low. The intermixing level increased with the increase of substrate concentration when substrate toxicity is relatively (e.g., *θ* < 0.03). However, when substrate toxicity is high (e.g., *θ* > 0.03), the intermixing level rarely shifted with substrate concentration, which is related to the formation of the Detoxifier-leading pattern (Supplementary Figure 28).

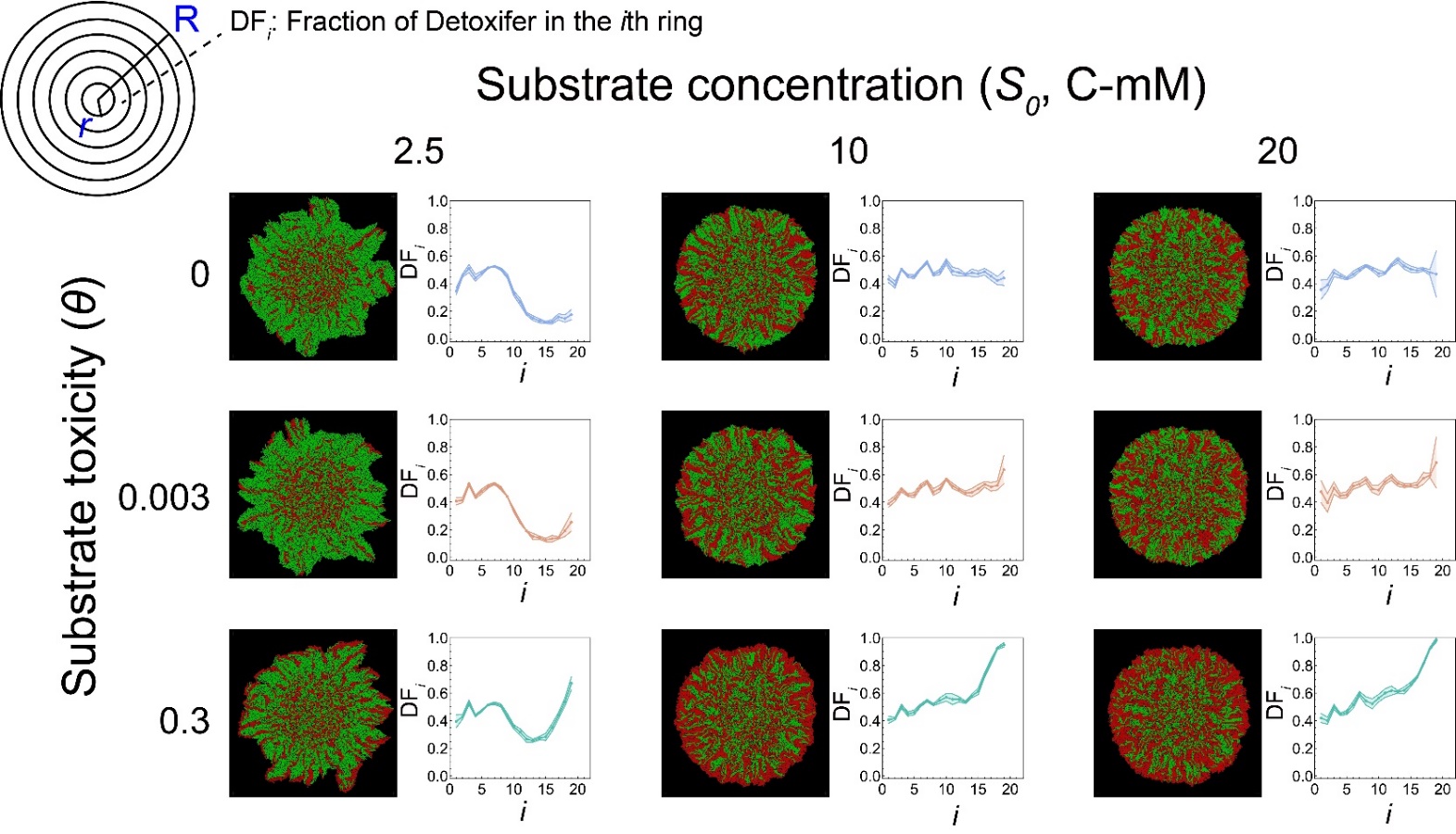

**Supplementary Figure 28** Analysis of the community structures along the radius of the colonies developed in our individual-based simulations. As shown in the top-left corner, each colony possessing a radius of *R* was divided into *i* concentric circles using a step of *r* (here, we set *r* = R/40). The community structures in each concentric circle were analyzed, and plotted as a function of the order of *i*. In the colony images, Detoxifier cells are shown in red, while Embezzler cells are shown in green.

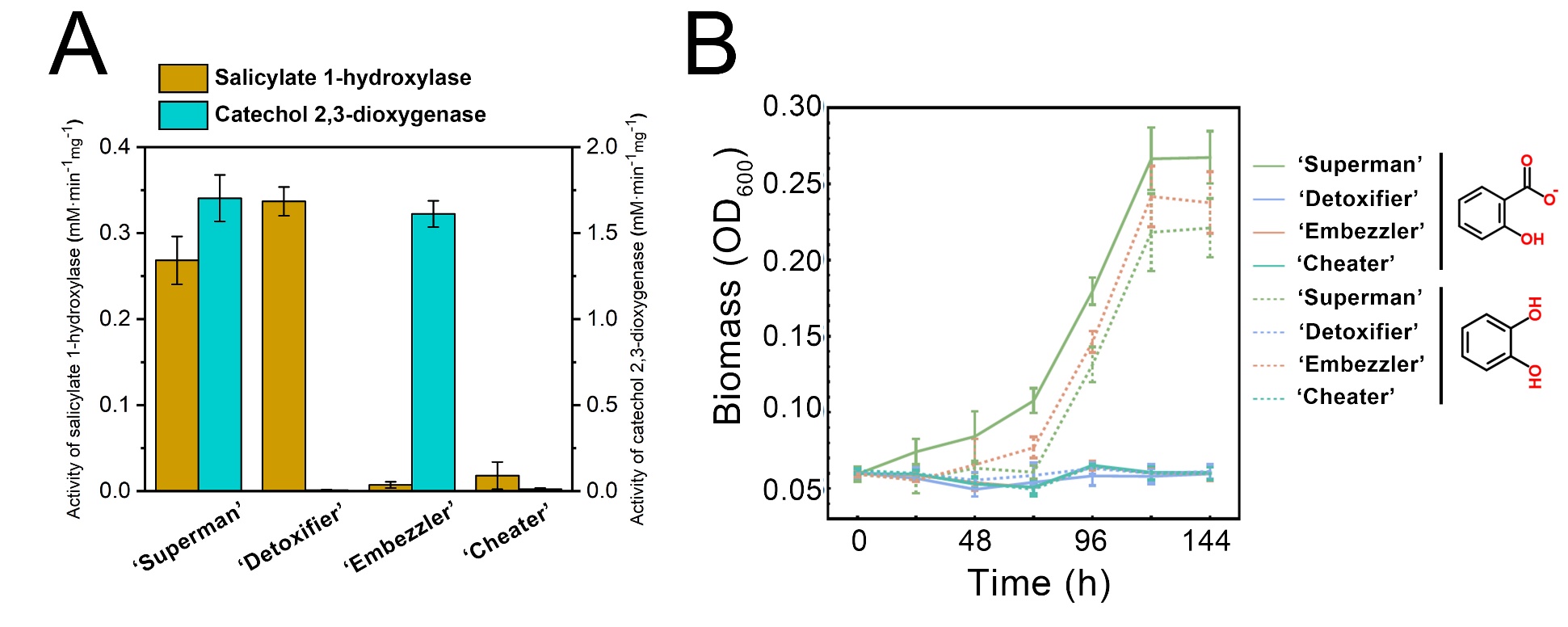

**Supplementary Figure 29** Identification of the strains used in this study. (A) Enzymic activity assays of salicylate 1-hydroxylase and catechol 2,3-dioxygenase were performed to verify the metabolism of the substrate, salicylate, and the intermediate, catechol, of the four strains. Three replicates were performed for each assay. (B) Growth dynamics from the mono-culture of four strains using salicylate (solid line) or catechol (dash line) as the sole carbon source. Five replicates were performed for each treatment. 10 C-mmol/L salicylate or catechol were used in the culture experiments.

**Supplementary video 1** A representative simulated dynamics with no substrate toxicity and low initial substrate concentration. (A) Dynamics of colony growth. Red cells are the from Detoxifier population, while green cells are the from Embezzler population. In this case, the initial substrate concentration is *S* = 2.5 C-mmol/L, while toxic strength *θ* = 0. Embezzler cells (Green) largely dominated the space in this case with a final fraction of 0.737 (4426 of 6000 cells). The values of other parameters are assigned with dimensional values listed in Supplementary Table 2**.** (B-D) Dynamics of the distributions of S (B), I (C), and P (D).

**Supplementary video 2** A representative simulated dynamics with no substrate toxicity and high initial substrate concentration. (A) Dynamics of colony growth. Red cells are the from Detoxifier population, while green cells are the from Embezzler population. In this case, the initial substrate concentration is *S* = 20 C-mmol/L, while toxic strength *θ* = 0. Detoxifier population (Red) possess a higher frequency than the case in Supplementary video 1, with a final fraction of 0.489 (2933 of 6000 cells). The values of other parameters are assigned with dimensional values listed in Supplementary Table 2**.** (B-D) Dynamics of the distributions of S (B), I (C), and P (D).

**Supplementary video 3** A representative simulated dynamics with high substrate toxicity and low initial substrate concentration. (A) Dynamics of colony growth. Red cells are the from Detoxifier population, while green cells are the from Embezzler population. In this case, the initial substrate concentration is *S* = 2.5 C-mmol/L, while toxic strength *θ* = 0.3. Embezzler cells (Green) dominated the space in this case with a final fraction of 0.605 (3629 of 6000 cells). The values of other parameters are assigned with dimensional values listed in Supplementary Table 2**.** (B-D) Dynamics of the distributions of S (B), I (C), and P (D).

**Supplementary video 4** A representative simulated dynamics with high substrate toxicity and high initial substrate concentration. (A) Dynamics of colony growth. Red cells are the from Detoxifier population, while green cells are the from Embezzler population. In this case, the initial substrate concentration is *S* = 20 C-mmol/L, while toxic strength *θ* = 0.3. Detoxifier cells (Red) dominated the space in this case with a final fraction of 0.621 (3726 of 6000 cells). The values of other parameters are assigned with dimensional values listed in Supplementary Table 2**.** (B-D) Dynamics of the distributions of S (B), I (C), and P (D).
